# Evolutionary branching under slowly evolving constraints

**DOI:** 10.64898/2026.09.21.753169

**Authors:** Hiroshi C. Ito

## Abstract

Biological populations have been evolving in multi-dimensional trait spaces. Ecological interactions can induce the evolutionary diversification of a biological population into distinct multiple populations, which is called evolutionary branching in adaptive dynamics theory. If possible mutations are restricted to certain directions due to developmental, physiological, or physical constraints, the resulting evolutionary dynamics may be restricted to subspaces with fewer dimensions than the original trait spaces. In general, such constraints are also evolving slowly, due to rare and slight mutational deviations from the constraint subspaces. In this sense, constraints are not strict. However, the method for describing non-strict constraints that is globally consistent has yet to be developed. This study proposes a general and flexible method for constructing a non-strict constraint subspace from the corresponding strict constraint subspace. This method enables efficient analysis of adaptive evolution including evolutionary branching under non-strict constraints in the framework of adaptive dynamics theory. Applying this method to a resource competition model shows that the shape of the carrying capacity distribution can be treated as a non-strict constraint curve, and that the likelihood of evolutionary branching is affected not only by the shape of the constraint curve but also by its slow evolution.

## 1. Introduction

Ecological interactions induce adaptive evolution of biological populations in multi-dimensional trait spaces, i.e., individual organisms are evolving in many traits simultaneously (Lande, 1979). Simultaneously, evolutionary constraints (i.e., trade-offs) often exist, such that a mutation improving one trait inevitably makes another trait worse (Tilman, 1982). Hence, adaptive evolution of populations in multi-dimensional trait spaces may be restricted to “constraint subspaces” with fewer dimensions (Schluter, 1996; deMazancourt and Dieckmann, 2004; Kisdi, 2006; Kisdi, 2015; Ito and Sasaki, 2016). The shapes of constraint subspaces can affect the likelihood of evolutionary diversification through ecological interaction (deMazancourt and Dieckmann, 2004; Kisdi, 2006; Kisdi, 2015; Ito and Sasaki, 2016). In general, such constraint subspaces are also evolving slowly, due to rare and slight mutational deviations from the constraint subspaces (Ito and Sasaki, 2023). In this sense, constraints are not strict. Ito and Sasaki (2020) described the local shapes of non-strict constraint subspaces as parabolic functions of traits and derived the conditions for evolutionary branching. However, the method for describing non-strict constraint subspaces that is globally consistent has yet to be developed. This study proposes a general and flexible method for describing non-strict constraint subspaces. Based on the method, we derive the conditions of evolutionary branching under non-strict constraint subspaces in the framework of adaptive dynamics theory. As an application example, we analyze a resource competition model under a non-strict constraint. The analysis shows that the carrying capacity distribution can be treated as a non-strict constraint subspace, and that the likelihood of evolutionary branching is affected not only by the shape of constraint subspace but also by its slow evolution. Numerically simulated evolution in this model typically shows repeated evolutionary branching and extinction (analogously to Ito and Dieckmann, 2007; Ito and Sasaki, 2023). For simplicity, we restrict our analysis in the main text to two-dimensional trait spaces, which can readily be extended to trait spaces and constraint subspaces of arbitrary dimensions (Appendix B).

## 2. Description method for non-strict constraints

For an arbitrary two-dimensional trait space **u** = (*X, Y*)^*T*^, we start with a strict constraint subspace expressed as a function *H*(**u**) = *Z*_0_ with *Z*_0_ being a constant, such that any mutant **u**′ = (*X* ′, *Y* ′) that emerges during evolutionary dynamics starting from a phenotype **u**_0_ = (*X*_0_, *Y*_0_)^*T*^ is restricted to a curve *H*(**u**′) = *H*(**u**_0_). To allow slight mutational deviations from the constraint curve, we assume a three-dimensional hyper space 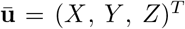 by adding the *Z*-axis to **u** = (*X, Y*)^*T*^, and assume a two-dimensional surface in it, defined by

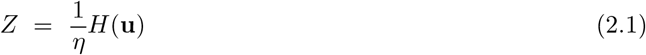

with a positive constant *η*, referred to as a “mutation mountain.” We also assume that the mutational covariance matrix for the hyper trait space is isotropic and constant, given by *σ*^2^**I**, inverse of which gives the mutational metric *σ*^-2^**I**. Then, the mutational square distance d*l*^2^ from 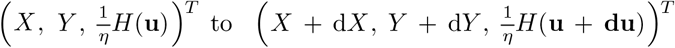 along the surface with an infinitesimal **du** = (d*X*, d*Y*) is given by

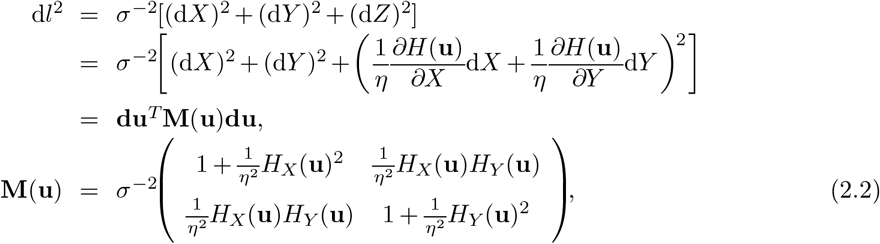

where **M**(**u**) is referred to as the mutational metric. From Eq. (2.2), we get the mutational covariance matrix as **V**(**u**) = **M**(**u**)^*-*1^,

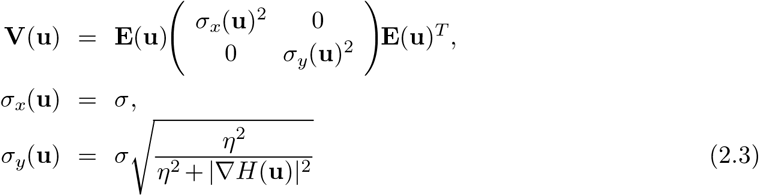

with

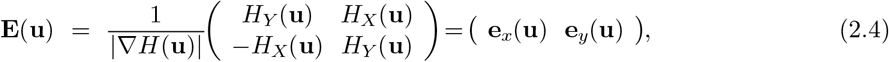

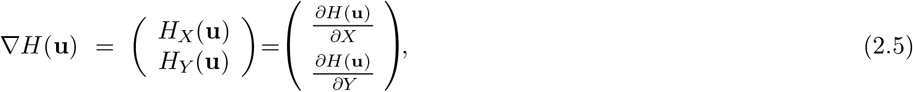

provided that ∣∇*H*∣ *≠* 0. As shown in Fig. 1c, a smaller *η* makes the constraint more strict, where *dl*^2^ is large for **du** orthogonal to the constraint curve 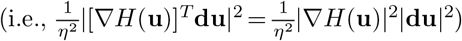 compared to **du** parallel to the constraint curve 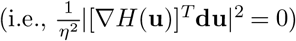. Hence, the exactly strict constraint is realized when *η* → 0.

**Figure 1.**
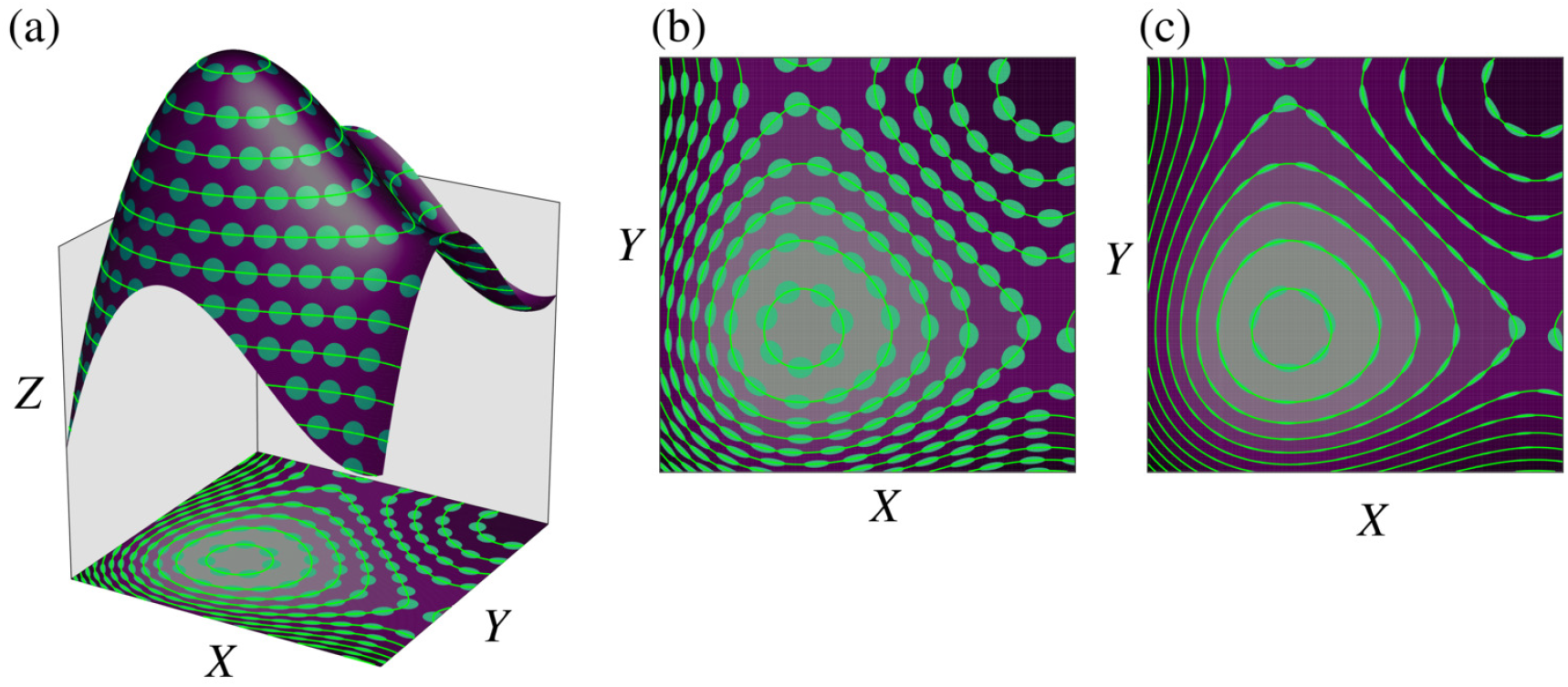
Describing a non-strict constraint by mutation mountain mapping

## 3. Conditions for evolutionary branching

### 3.1. Invasion fitness function

We define the population dynamics for arbitrary coexisting phenotypes **u**_1_,*…*, **u**_*L*_ with their population sizes denoted by *n*_1_,*…, n*_*L*_ as

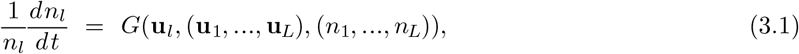

where *G*(**u**_*5*_, (**u**_1_, …, **u**_*L*_), (*n*_1_, …, *n*_*L*_)) describes the fitness for **u**_*5*_. By assuming their equilibrium population sizes denoted by 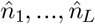 we define the invasion fitness for a mutant phenotype **u**′ as

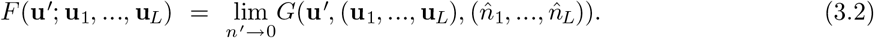

### 3.2. Local normalization of invasion fitness function

As shown by Ito and Sasaki (2020), the linear dependency of **V**(**u**) = **M**(**u**)^*−*1^ on **u** around an arbitrary point û = (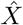, *Ŷ*)^*T*^ can be canceled by local coordinate normalization, composed of coordinate rotation into coordinates **s** = (*x, y*)^*T*^, defined by

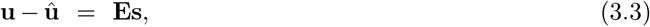

with **E** = **E**(**û**) and a nonlinear coordinate transformation into coordinates 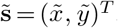, defined by,

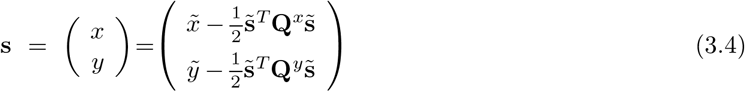

with

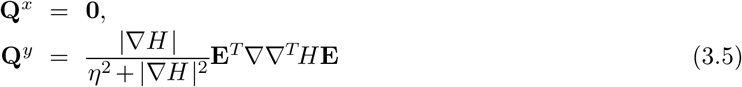

with

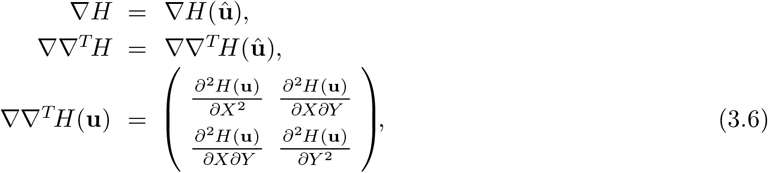

where **Q**^*x*^ and **Q**^*y*^ are referred to as distortion matrices” (see Appendix A).

We denote by *F* (**u**′, **u**) an invasion fitness function of mutant phenotype **u**′ under existence of a monomorphic resident with phenotype **u**. By using Eq. (2.5), we express *F* (**u**′, **u**) in the rotated coordinates **s** centered at an arbitrary point **û**, as

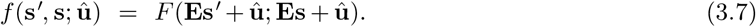

Since *f* (**s, s**; **û**) = 0 holds for any **s**, Eq. (2.19) can be expanded as

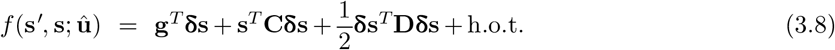

with

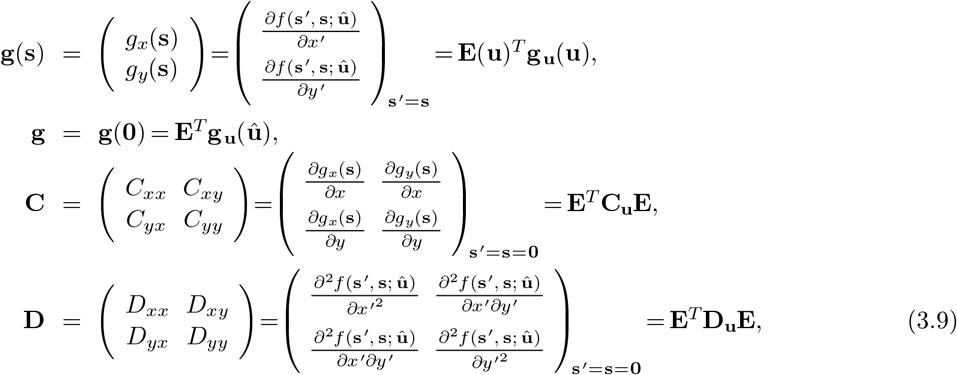

and

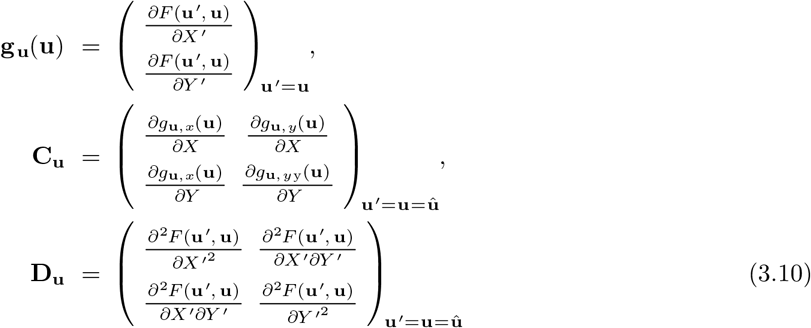

(Ito and Dieckmann, 2014; Ito and Sasaki, 2016). Substituting Eq. (2.6) into Eq. (2.20), we get the normalized invasion fitness function in the rotated-geodesic coordinates 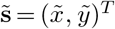.

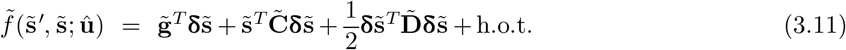

with

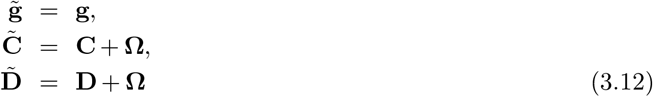

where

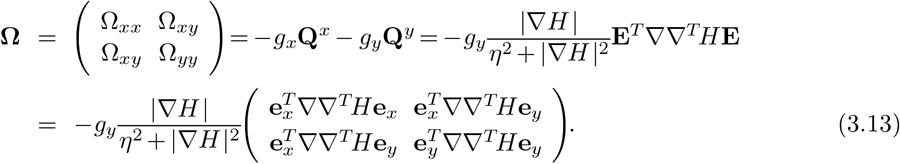

In the rotated-geodesic coordinates ~**s**, the mutation distribution can be locally approximated as a constant Gaussian mutation distribution with its covariance matrix given by **M**_**s**_(**0**; û) ^*−*1^. Thus, existing conditions for evolutionary branching points, lines, and areas can be applied to 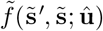 as shown by Ito and Sasaki (2020).

### 3.3. Conditions for evolutionary branching under a strict constraint

When *η* = 0 holds, the constraint is strict. Then, from Eqs. (3.10-3.12), we obtain the evolutionary branching point conditions under a strict constraint (Ito and Sasaki, 2016) as

i. **û** is evolutionary singular along 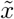, satisfying

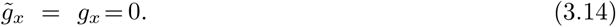
ii. **û**is convergence stable along 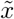, satisfying

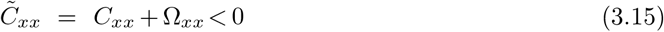

with 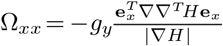.
iii. **û** is evolutionary unstable along 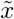, satisfying

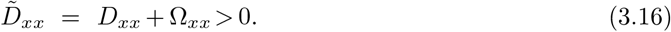

These conditions are illustrated in Fig. 2a-c, which are equivalent to the conditions developed by Ito and Sasaki (2016).

**Figure 2.**
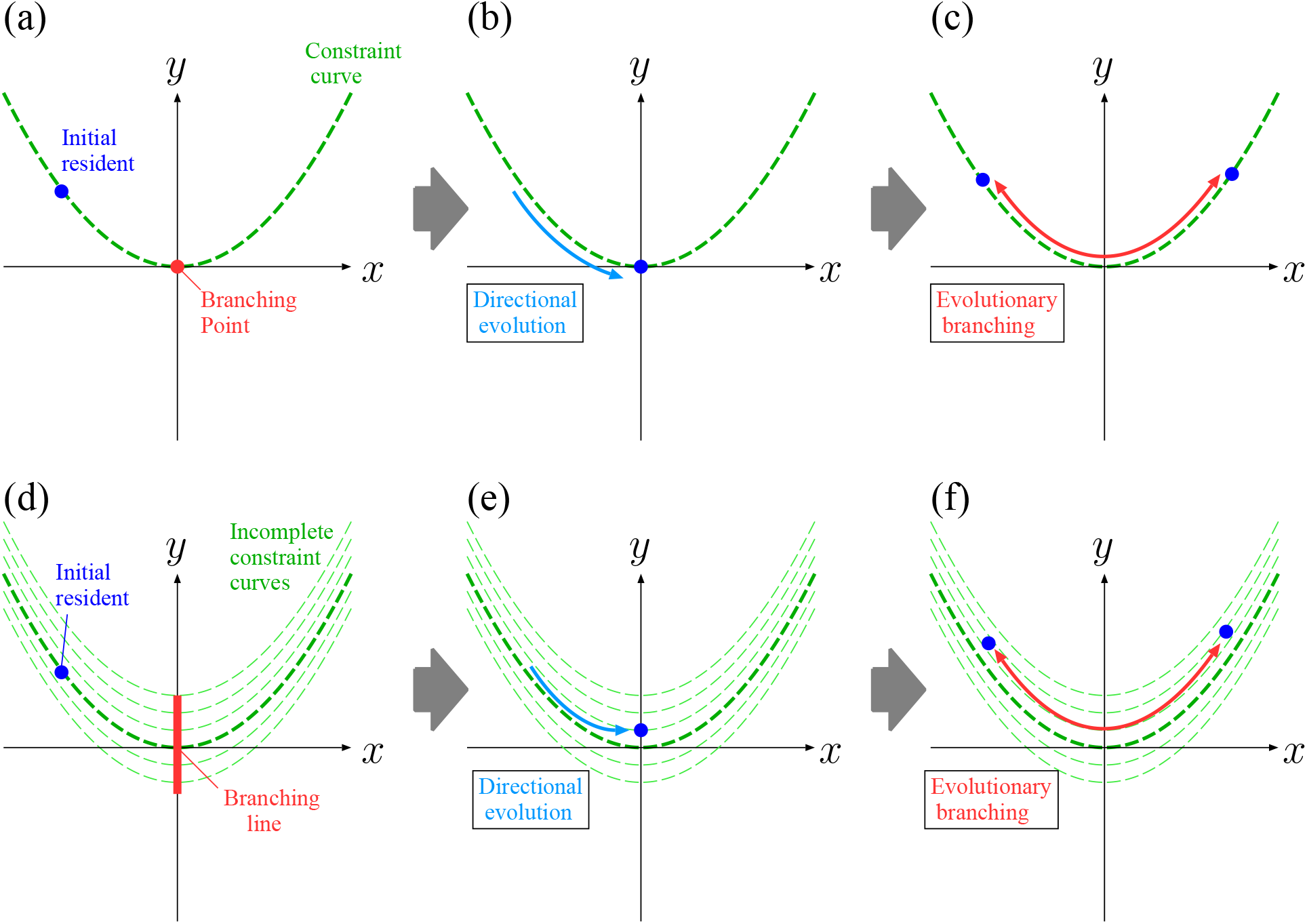
Evolutionary branching under strict and non-strict constraints in the locally rotated coordinate system **s** = (*x, y*)^*T*^

**Figure 3.**
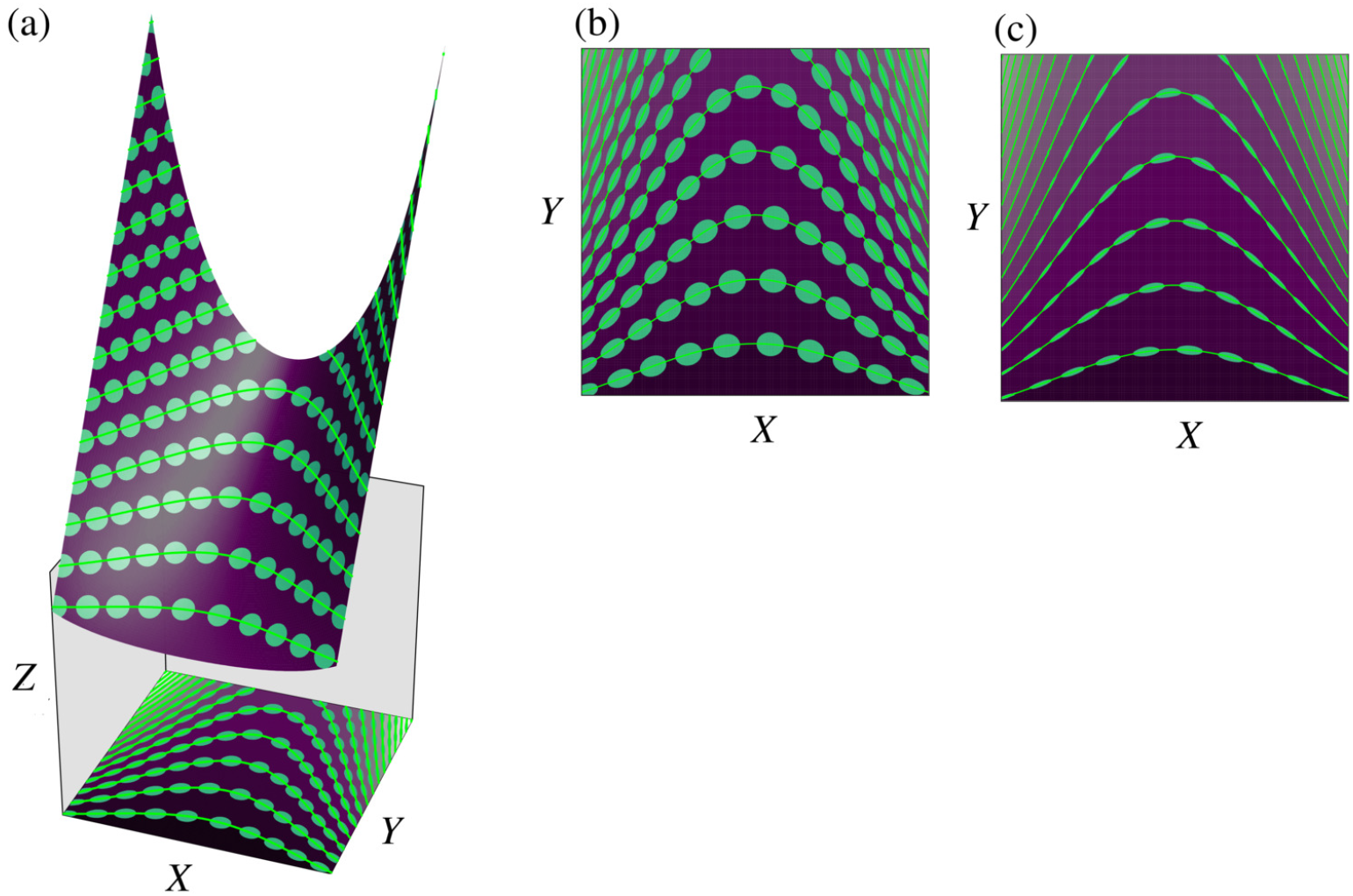
Describing a non-strict constraint by mutation mountain mapping in the application example

### 3.4. Conditions for evolutionary branching under a non-strict constraint

From Eqs. (3.10-3.12), we obtain the evolutionary branching line conditions under a non-strict constraint (Ito and Sasaki, 2020) as

i. the sensitivity of 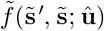 to single mutational changes of 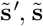 is significantly lower in 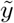 than in 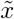, satisfying

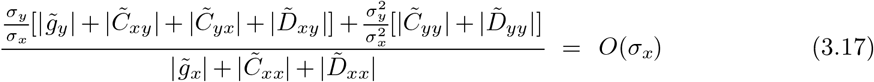

with 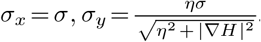.
ii. **û** is evolutionary singular along 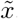, satisfying

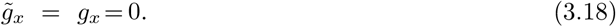
iii. **û** is convergence stable along 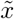, satisfying

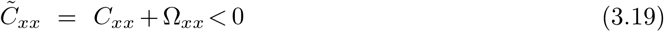

with 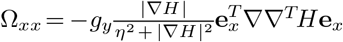.
iv. **û** is evolutionary unstable along *x*~, satisfying

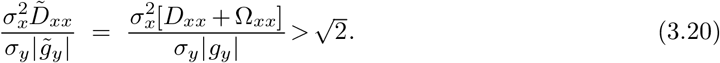

These conditions are illustrated in Fig. 2d-f. Note these conditions under *η* = 0 are equivalent to Eqs. (3.13-3.15).

## 4. Application example

We consider a classical MacArthur-Levins type resource competition model, defined on a one-dimensional trait space *X*:

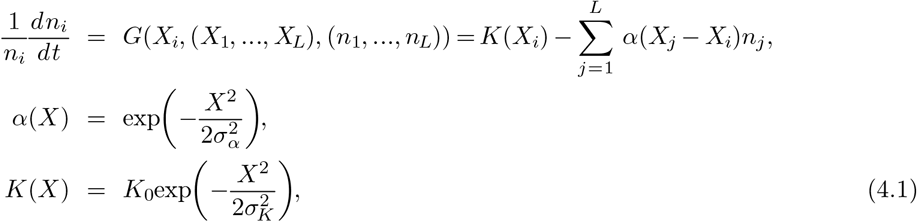

where *α*(*X*) and *K*(*X*) describe the competition kernel and carrying capacity distribution, respectively. To see how our method works, we modify this model into one defined on a two-dimensional trait space **u** = (*X, Y*)^*T*^ :

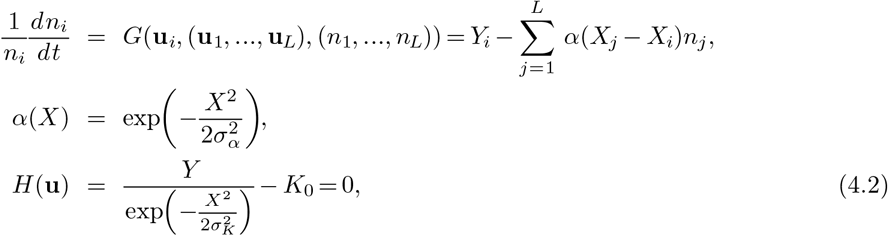

with *Y >* 0. Then we can analyze adaptive evolution in this model under a non-strict constraint by assuming a mutation mountain

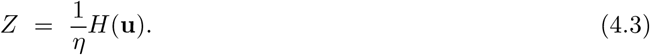

### 4.1. Adaptive evolution under a non-strict constraint

### 4.2. Adaptive evolution under a strict constraint

By using Eq. (3.2) we obtain from Eq. (4.1) the invasion fitness function:

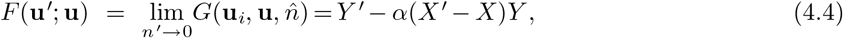

from which we get

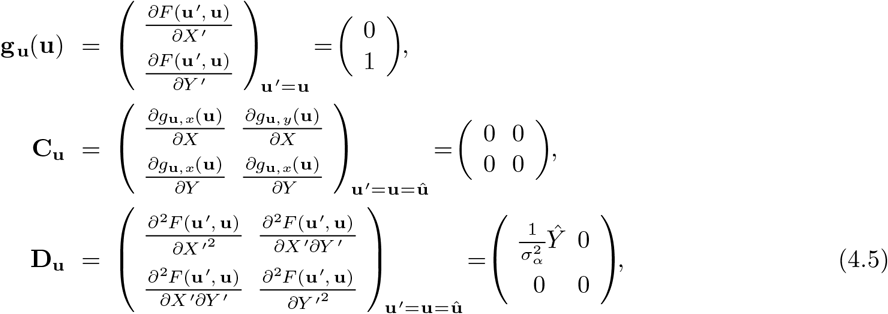

Also we see from Eq. (4.2) that

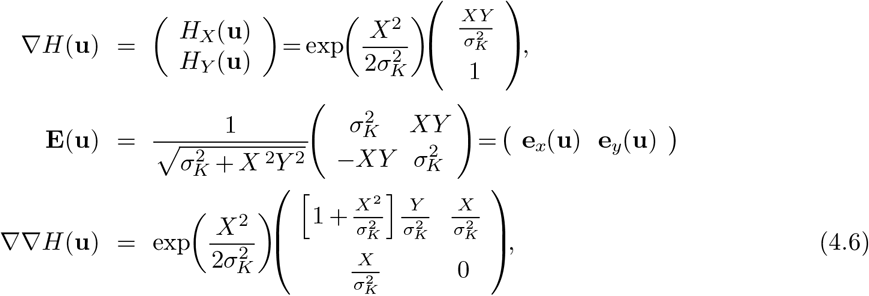

from which we get

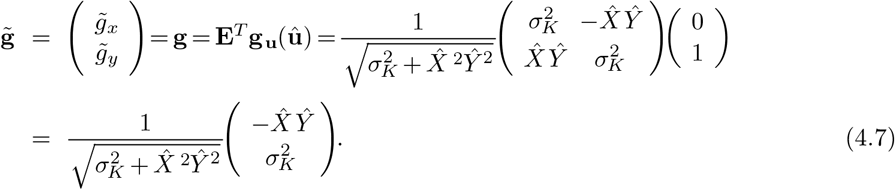

Hence *g*_*x*_ = 0 holds only when 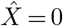. In this case, we see from 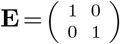 that

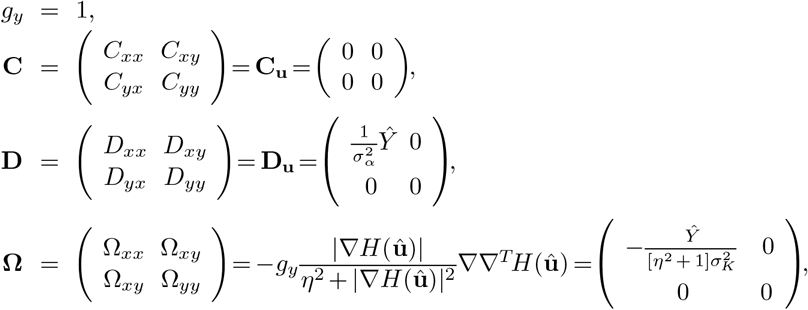

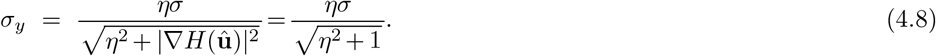

Therefore, conditions i,iii, and iv for branching lines are obtained as

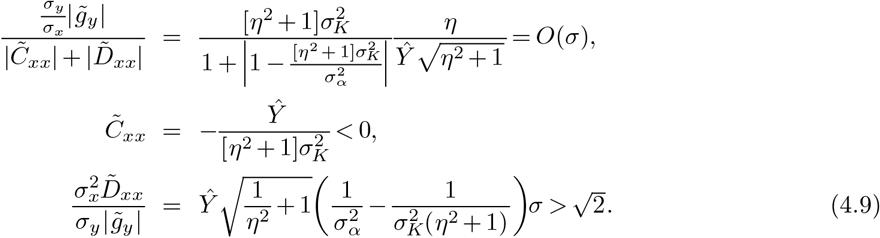

The occurrence of evolutionary branching in numerically simulated evolutions under rare and small mutations are effectively predicted by Eq. (4.9), as shown in Fig. 4 (see Ito and Sasaki (2020) for the simulation algorithm).

**Figure 4.**
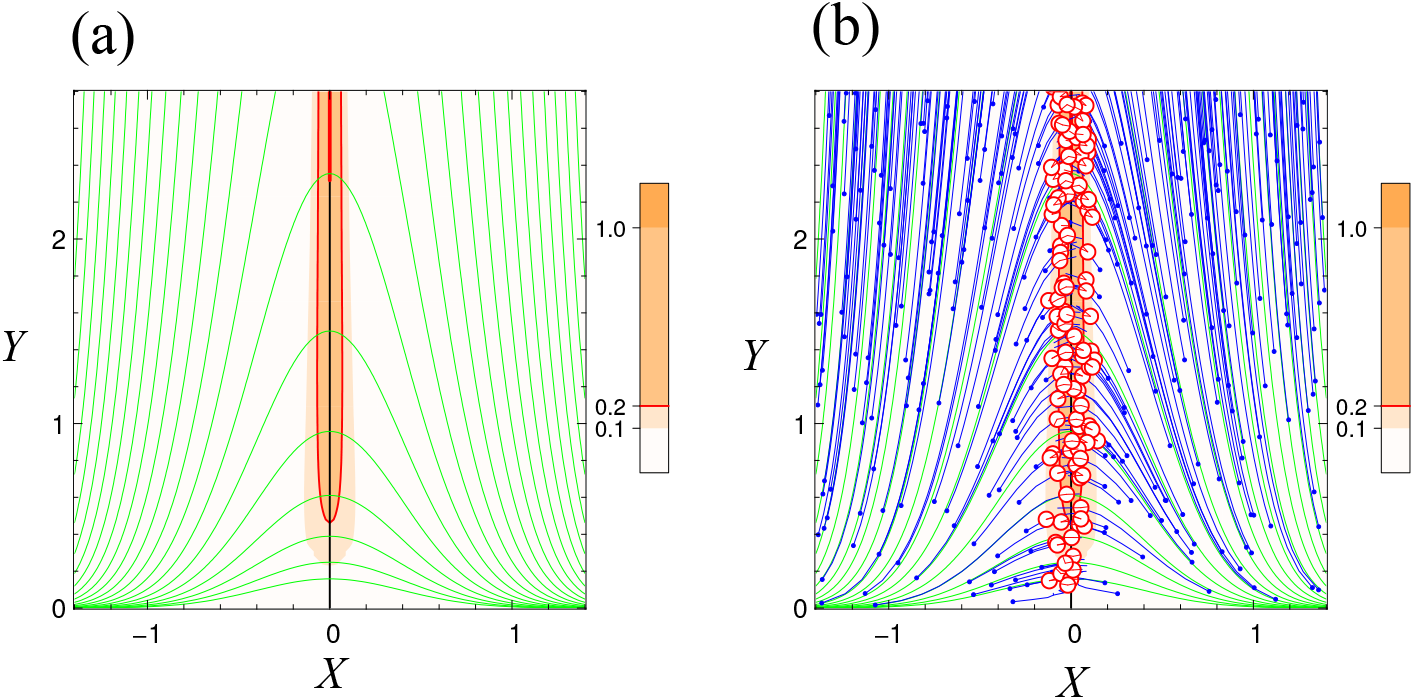
(a) Predicted evolutionary branching lines and areas under non-strict constraint. (b) Numerically simulated evolutionary branching. *σ*_*K*=0.4_, *σ*_*a*=0.2_, *σ* = 0.01, *K*_0_ = 1, *η*= 0.1.

The initial evolutionary branching is typically followed by repeated branching and extinction as shown in Fig. 5. When the constraint is strict, i.e., *η* = 0 holds, Eq. (4.9) is reduced to

**Figure 5.**
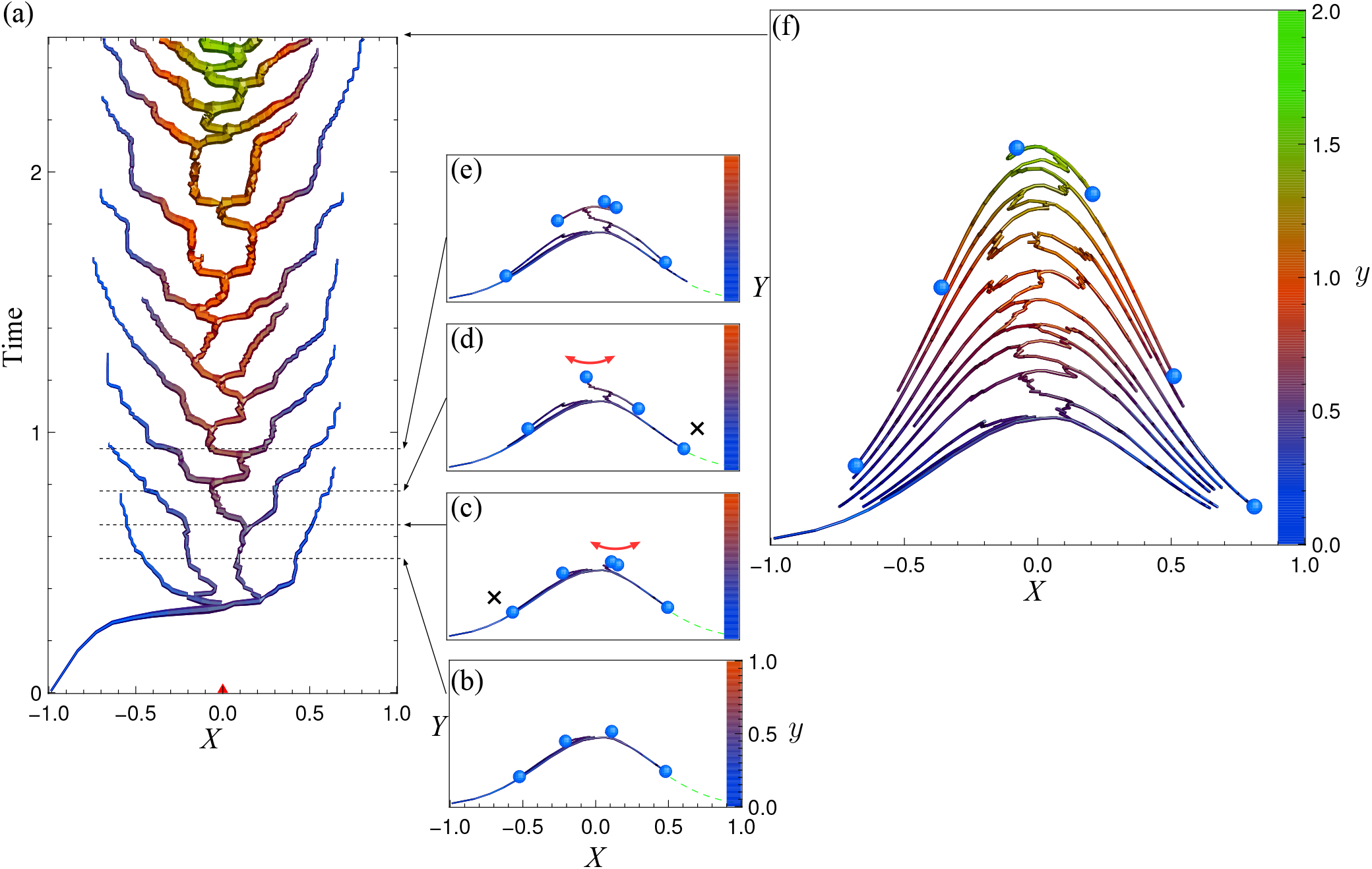
Evolutionary dynamics in example under a non-strict constraint.

**Figure 6.**
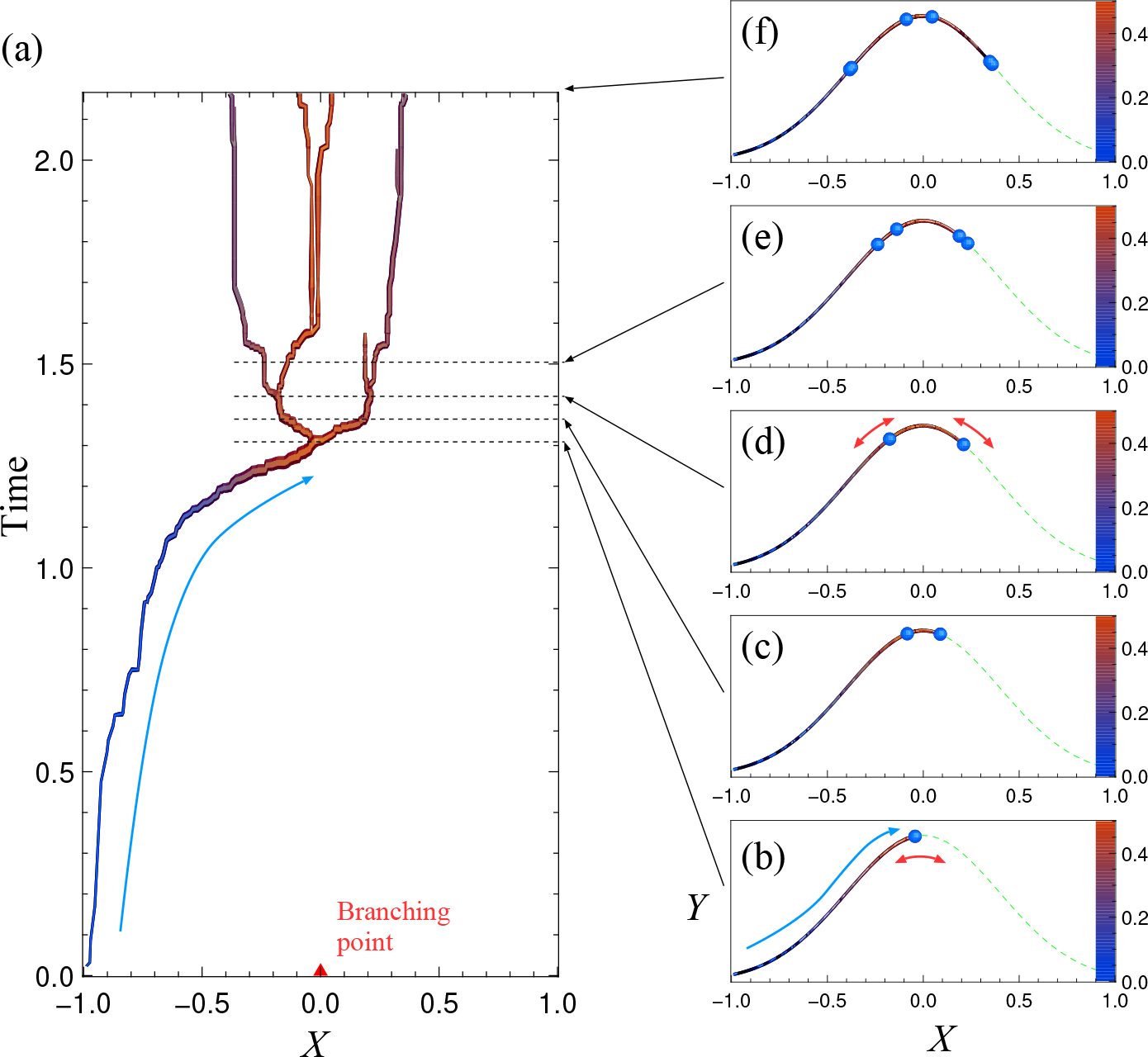
Early evolutionary dynamics in example under a strict constraint

**Figure 7.**
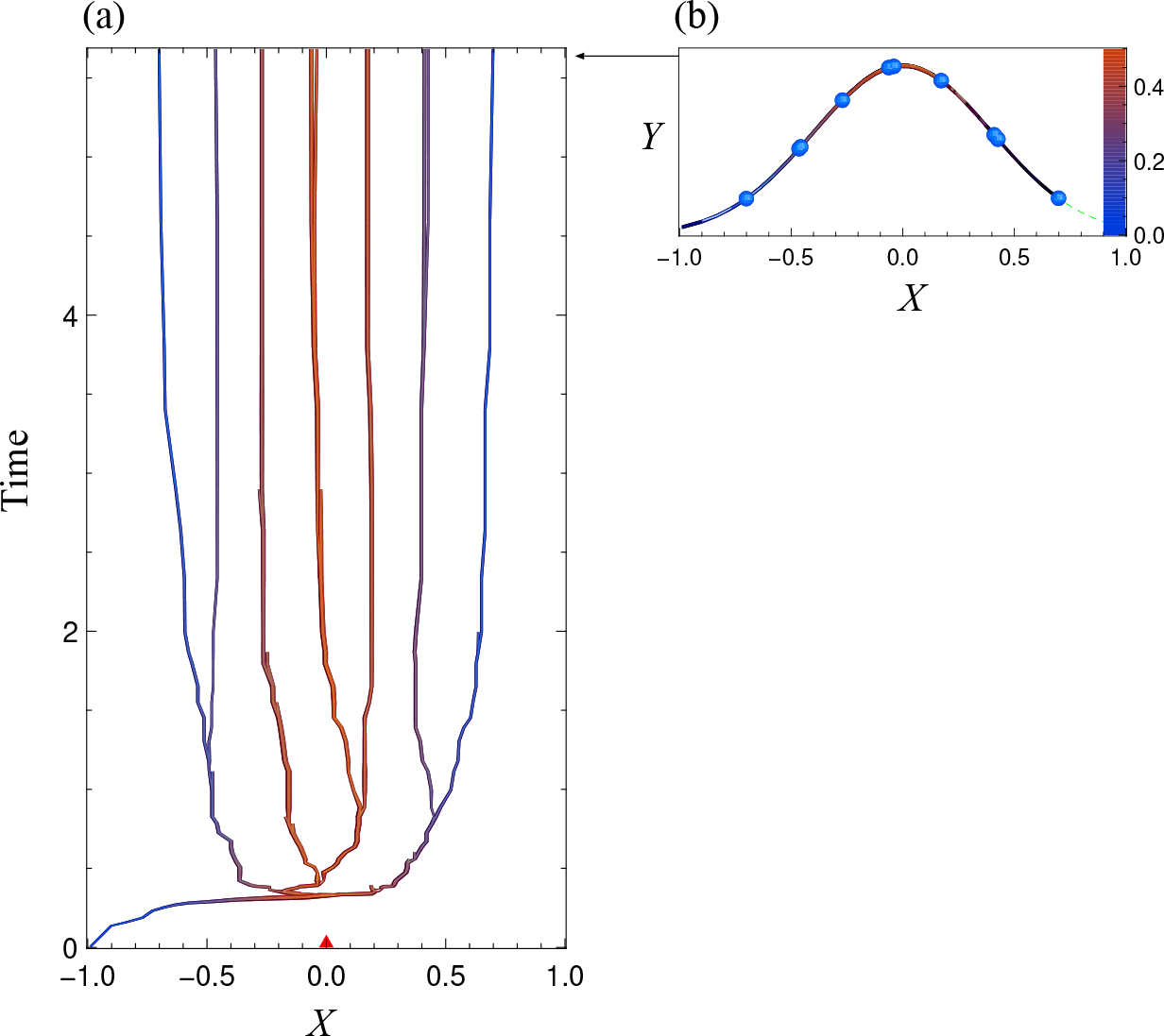
Long-term evolutionary dynamics in example under a strict constraint

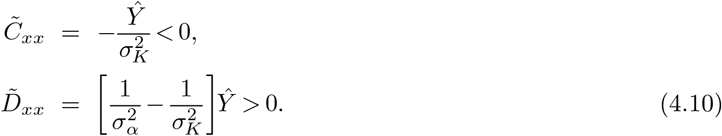

Note that Eq. (4.10) is identical to the branching point condition in a one-dimensional trait space analyzed by Dieckmann and Doebeli (1999). The numerically simulated evolutionary branching under this strict constraint is shown in Figs. 4 and 5 (see Ito and Sasaki (2020) for the simulation algorithm).

## 5. Non-strict constraints in higher-dimensional trait spaces

Our method for describing non-strict constraints are readily extended to trait spaces of higher dimensions (Appendix B). Notably, the adaptive evolution in trait spaces under non-strict constraints can be analyzed not only by local nonlinear coordinate normalization shown here (Ito and Sasaki, 2020), but also by applying a Lagrange multiplier method directly to mutation mountain surfaces (Ito and Sasaki, 2016). Both analyses are mathematically equivalent.

## 6. Discussion

Adaptive evolution of biological populations in multi-dimensional trait spaces may be restricted to constraint subspaces with fewer dimensions (Schluter, 1996; deMazancourt and Dieckmann, 2004; Kisdi, 2006; Kisdi, 2015; Ito and Sasaki, 2016). Such constraints are non-strict constraints in general, which are evolving slowly (Ito and Sasaki, 2020; Ito and Sasaki, 2023). Although such evolutionary changes of constraints may affect evolutionary branching, the method for describing non-strict constraints that is globally consistent has yet to be developed. The present study developed a general and flexible method for constructing a non-strict constraint subspace from the corresponding strict constraint subspace, called the mutation mountain mapping. This method (1) constructs a hyper trait space by adding pseudo axes to a focal trait space, (2) assumes a hyper constraint surface (i.e., a mutation mountain) in the hyper trait space, to which all possible mutants are restricted, but along which the mutational metric is constant and isotropic, and (3) projects the hyper constraint surface orthogonally onto the trait space. As shown by the application example, our method allows efficient analysis of adaptive evolution including evolutionary branching in the framework of adaptive dynamics theory. The application example to a resource competition model shows that the shape of the carrying capacity distribution can be treated as a non-strict constraint curve, and that the likelihood of evolutionary branching is affected not only by the shape of the constraint curve but also by its slow evolution.

