## Appendices for "Evolutionary branching under slowly evolving constraints"

### Appendices for
### **Evolutionary branching under slowly evolving constraints**

Hiroshi C. Ito

Research Center for Integrative Evolutionary Science, SOKENDAI, Hayama, Kanagawa 240-0193, Japan

### S1. Appendix A: 2-Dim trait spaces with one constraint

#### S1.1. Non-strict constraint

For an arbitrary two-dimensional trait space $\mathbf{u}=(X,Y)^T$, we assume a three-dimensional hyper space $\bar{\mathbf{u}}=(X,Y,Z)^T$ by adding the $Z$-axis to $\mathbf{u}=(X,Y)^T$, and assume a two-dimensional surface in it, defined by $(X,Y,H(\mathbf{u}))^T$, referred to as a "mutation mountain." We also assume that the mutational metric for the hyper trait space is isotropic and constant, given by $\sigma^{-2}\mathbf{I}$. Then we can express the mutational square distance $\mathrm{d}l^2$ from $(X,Y,H(\mathbf{u}))^T$ to $(X+\mathrm{d}X,Y+\mathrm{d}Y,H(\mathbf{u}+\mathbf{du}))^T$ along the surface with an infinitesimal $\mathbf{du}=(\mathrm{d}X,\mathrm{d}Y)$ as

$$\begin{aligned}
\mathrm{d}l^2 &= \sigma^{-2}[(\mathrm{d}X)^2+(\mathrm{d}Y)^2+(\mathrm{d}Z)^2] \\
&= \sigma^{-2}\left[(\mathrm{d}X)^2+(\mathrm{d}Y)^2+\left(\frac{\partial H(\mathbf{u})}{\partial X}\mathrm{d}X+\frac{\partial H(\mathbf{u})}{\partial Y}\mathrm{d}Y\right)^2\right] \\
&= \sigma^{-2}\begin{pmatrix} \mathrm{d}X \\ \mathrm{d}Y \end{pmatrix}\begin{pmatrix} 1+H_X(\mathbf{u})^2 & H_X(\mathbf{u})H_Y(\mathbf{u}) \\ H_X(\mathbf{u})H_Y(\mathbf{u}) & 1+H_Y(\mathbf{u})^2 \end{pmatrix}\begin{pmatrix} \mathrm{d}X \\ \mathrm{d}Y \end{pmatrix} \\
&= \mathbf{du}^T\mathbf{M}(\mathbf{u})\mathbf{du},
\end{aligned} \tag{S1.1}$$

from which we get

$$\begin{aligned}
\mathbf{V}(\mathbf{u}) &= \mathbf{M}(\mathbf{u})^{-1}=\mathbf{E}(\mathbf{u})\begin{pmatrix} \sigma_x^2 & 0 \\ 0 & \sigma_y^2 \end{pmatrix}\mathbf{E}(\mathbf{u})^T, \\
\sigma_x &= \sigma, \\
\sigma_y &= \sigma\sqrt{\frac{\eta^2}{\eta^2+|\nabla H(\mathbf{u})|^2}}
\end{aligned} \tag{S1.2}$$

with

$$\mathbf{E}(\mathbf{u}) = \frac{1}{|\nabla H(\mathbf{u})|}\begin{pmatrix} H_Y(\mathbf{u}) & H_X(\mathbf{u}) \\ -H_X(\mathbf{u}) & H_Y(\mathbf{u}) \end{pmatrix} = \begin{pmatrix} \mathbf{e}_x(\mathbf{u}) & \mathbf{e}_y(\mathbf{u}) \end{pmatrix},$$
$$\nabla H(\mathbf{u}) = \begin{pmatrix} H_X(\mathbf{u}) \\ H_Y(\mathbf{u}) \end{pmatrix} = \begin{pmatrix} \frac{\partial H(\mathbf{u})}{\partial X} \\ \frac{\partial H(\mathbf{u})}{\partial Y} \end{pmatrix}. \tag{S1.3}$$

As shown by Ito and Sasaki(2020), the linear dependency of $\mathbf{M}(\mathbf{u})$ on $\mathbf{u}$ at $\hat{\mathbf{u}}$ can be canceled by local coordinate normalization, composed of coordinate rotation into coordinates $\mathbf{s}$, defined by

$$\mathbf{u} - \hat{\mathbf{u}} = \mathbf{E}\mathbf{s}, \tag{S1.4}$$

and a nonlinear coordinate transformation into coordinates $\tilde{\mathbf{s}}$, defined by,

$$\mathbf{s} = \tilde{\mathbf{s}} - \frac{1}{2}\tilde{\mathbf{s}}\left\{ \begin{matrix} \mathbf{Q}^x \\ \mathbf{Q}^y \end{matrix} \right\}\tilde{\mathbf{s}}.$$
$$= \tilde{\mathbf{s}} - \frac{1}{2}\tilde{\mathbf{s}}\mathsf{Q}\tilde{\mathbf{s}}. \tag{S1.5}$$

where $\mathsf{Q}$ is a column vector with its components given by symmetric matrices, called a matrix-vector (see Appendix C), expressed as

$$\mathsf{Q} = \left\{ \begin{matrix} \mathbf{Q}^x \\ \mathbf{Q}^y \end{matrix} \right\} = \left\{ \begin{matrix} \begin{pmatrix} Q^x_{xx} & Q^x_{xy} \\ Q^x_{xy} & Q^x_{yy} \end{pmatrix} \\ \begin{pmatrix} Q^y_{xx} & Q^y_{xy} \\ Q^y_{xy} & Q^y_{yy} \end{pmatrix} \end{matrix} \right\}, \tag{S1.6}$$

where $\mathbf{Q}^x$ and $\mathbf{Q}^y$ are referred to as “distortion matrices”.

#### S1.2. Distortion matrices

The $Q^\gamma_{\alpha\beta}$ for $\alpha, \beta, \gamma \in \{x, y\}$ are given by

$$Q^\gamma_{\alpha\beta} = \frac{1}{2}\hat{V}_{\mathbf{s}\gamma x}[A^\beta_{x\alpha} + A^\alpha_{x\beta} - A^x_{\alpha\beta}] + \frac{1}{2}\hat{V}_{\mathbf{s}\gamma y}[A^\beta_{y\alpha} + A^\alpha_{y\beta} - A^y_{\alpha\beta}] \tag{S1.7}$$

with

$$\begin{pmatrix} \hat{V}_{\mathbf{s}xx} & \hat{V}_{\mathbf{s}xy} \\ \hat{V}_{\mathbf{s}xy} & \hat{V}_{\mathbf{s}yy} \end{pmatrix} = \mathbf{V}_\mathbf{s}(\mathbf{0};\hat{\mathbf{u}}) = \mathbf{M}_\mathbf{s}(\mathbf{0};\hat{\mathbf{u}})^{-1},$$
$$\begin{pmatrix} A^x_{xx} & A^x_{xy} \\ A^x_{xy} & A^x_{yy} \end{pmatrix} = \mathbf{A}^x = \left[\frac{\partial \mathbf{M}_\mathbf{s}(\mathbf{s};\hat{\mathbf{u}})}{\partial x}\right]_{\mathbf{s}=\mathbf{0}},$$
$$\begin{pmatrix} A^y_{xx} & A^y_{xy} \\ A^y_{xy} & A^y_{yy} \end{pmatrix} = \mathbf{A}^y = \left[\frac{\partial \mathbf{M}_\mathbf{s}(\mathbf{s};\hat{\mathbf{u}})}{\partial y}\right]_{\mathbf{s}=\mathbf{0}}. \tag{S1.8}$$

As shown in Appendix D, Eq. (S1.8) can alternatively be expressed as

$$\mathsf{Q} = \frac{1}{2}\langle \mathbf{M}_\mathbf{s}(\mathbf{0};\hat{\mathbf{u}})^{-1}, \mathsf{A}^r + \mathsf{A}^c - \mathsf{A}\rangle \tag{S1.9}$$

where $\mathsf{A}$ and its transpositions $\mathsf{A}^r$ and $\mathsf{A}^c$ are defined by

$$\mathsf{A} = \left\{ \begin{array}{c} \mathbf{A}^x \\ \mathbf{A}^y \end{array} \right\} = \left\{ \begin{array}{c} \begin{pmatrix} A^x_{xx} & A^x_{xy} \\ A^x_{xy} & A^x_{yy} \end{pmatrix} \\ \begin{pmatrix} A^y_{xx} & A^y_{xy} \\ A^y_{xy} & A^y_{yy} \end{pmatrix} \end{array} \right\},$$

$$\mathsf{A}^r = \left\{ \begin{array}{c} \begin{pmatrix} A^x_{xx} & A^x_{xy} \\ A^y_{xx} & A^y_{xy} \end{pmatrix} \\ \begin{pmatrix} A^x_{xy} & A^x_{yy} \\ A^y_{xy} & A^y_{yy} \end{pmatrix} \end{array} \right\}, \quad \mathsf{A}^c = \left\{ \begin{array}{c} \begin{pmatrix} A^x_{xx} & A^y_{xx} \\ A^x_{xy} & A^y_{xy} \end{pmatrix} \\ \begin{pmatrix} A^x_{xy} & A^y_{xy} \\ A^x_{yy} & A^y_{yy} \end{pmatrix} \end{array} \right\} \tag{S1.10}$$

(see Appendix C).

For convenience, we express the mutation mountain in the rotated coordinates $\mathbf{s}$ as

$$h(\mathbf{s}) = \frac{1}{\eta} H(\mathbf{E}\mathbf{s} + \hat{\mathbf{u}}), \tag{S1.11}$$

where its derivatives are denoted by

$$\begin{aligned}
\nabla h(\mathbf{s}) &= \begin{pmatrix} \frac{\partial h(\mathbf{s})}{\partial x} \\ \frac{\partial h(\mathbf{s})}{\partial y} \end{pmatrix} = \begin{pmatrix} h_x(\mathbf{s}) \\ h_y(\mathbf{s}) \end{pmatrix}, \\
\nabla h(\mathbf{0}) &= \nabla h = \begin{pmatrix} h_x \\ h_y \end{pmatrix} = \begin{pmatrix} 0 \\ h_y \end{pmatrix}, \\
\nabla\nabla^T h(\mathbf{s}) &= \begin{pmatrix} \frac{\partial^2 h(\mathbf{s})}{\partial x^2} & \frac{\partial^2 h(\mathbf{s})}{\partial x \partial y} \\ \frac{\partial^2 h(\mathbf{s})}{\partial x \partial y} & \frac{\partial^2 h(\mathbf{s})}{\partial y^2} \end{pmatrix} = \begin{pmatrix} h_{\mathrm{xx}}(\mathbf{s}) & h_{\mathrm{xy}}(\mathbf{s}) \\ h_{\mathrm{xy}}(\mathbf{s}) & h_{\mathrm{yy}}(\mathbf{s}) \end{pmatrix}, \\
\nabla\nabla^T h(\mathbf{0}) &= \nabla\nabla^T h = \begin{pmatrix} h_{\mathrm{xx}} & h_{\mathrm{xy}} \\ h_{\mathrm{xy}} & h_{\mathrm{yy}} \end{pmatrix}.
\end{aligned} \tag{S1.12}$$

Then we express the mutational metric in the rotated coordinates as

$$\begin{aligned}
\mathbf{M}_{\mathbf{s}}(\mathbf{s}; \hat{\mathbf{u}}) &= \sigma^{-2} \begin{pmatrix} 1 + [h_x(\mathbf{s})]^2 & [h_x(\mathbf{s})][h_y(\mathbf{s})] \\ [h_x(\mathbf{s})][h_y(\mathbf{s})] & 1 + [h_y(\mathbf{s})]^2 \end{pmatrix}, \\
\mathbf{M}_{\mathbf{s}}(\mathbf{0}; \hat{\mathbf{u}}) &= \begin{pmatrix} \sigma^{-2} & 0 \\ 0 & \sigma^{-2}[1 + h_y^2] \end{pmatrix}, \\
\mathbf{M}_{\mathbf{s}}(\mathbf{0}; \hat{\mathbf{u}})^{-1} &= \begin{pmatrix} \sigma^2 & 0 \\ 0 & \frac{\sigma^2}{1 + |h_y|^2} \end{pmatrix}.
\end{aligned} \tag{S1.13}$$

We calculate the first derivatives of $\mathbf{M}_{\mathbf{s}}(\mathbf{s}; \hat{\mathbf{u}})$ as

$$\begin{aligned}
\mathbf{A}^x &= \left[ \frac{\partial \mathbf{M}_{\mathbf{s}}(\mathbf{s}; \hat{\mathbf{u}})}{\partial x} \right]_{\mathbf{s}=\mathbf{0}} = \sigma^{-2} \begin{pmatrix} 2h_x h_{\mathrm{xx}} & h_y h_{\mathrm{xx}} \\ h_x h_{\mathrm{xy}} + h_y h_{\mathrm{xx}} & 2h_y h_{\mathrm{xy}} \end{pmatrix} = \sigma^{-2} \begin{pmatrix} 0 & h_y h_{\mathrm{xx}} \\ h_y h_{\mathrm{xx}} & 2h_y h_{\mathrm{xy}} \end{pmatrix}, \\
\mathbf{A}^y &= \left[ \frac{\partial \mathbf{M}_{\mathbf{s}}(\mathbf{s}; \hat{\mathbf{u}})}{\partial y} \right]_{\mathbf{s}=\mathbf{0}} = \sigma^{-2} \begin{pmatrix} 2h_x h_{\mathrm{xy}} & h_x h_{\mathrm{yy}} + h_y h_{\mathrm{xy}} \\ h_x h_{\mathrm{yy}} + h_y h_{\mathrm{xy}} & 2h_y h_{\mathrm{yy}} \end{pmatrix} = \sigma^{-2} \begin{pmatrix} 0 & h_y h_{\mathrm{xy}} \\ h_y h_{\mathrm{xy}} & 2h_y h_{\mathrm{yy}} \end{pmatrix},
\end{aligned} \tag{S1.14}$$

from which we get

$$\mathsf{A} = \sigma^{-2}\left\{\begin{matrix}\begin{pmatrix}0 & h_y h_{\mathrm{xx}}\\ h_y h_{\mathrm{xx}} & 2h_y h_{\mathrm{xy}}\end{pmatrix}\\ \begin{pmatrix}0 & h_y h_{\mathrm{xy}}\\ h_y h_{\mathrm{xy}} & 2h_y h_{\mathrm{yy}}\end{pmatrix}\end{matrix}\right\} = \left\{\begin{matrix}\begin{pmatrix}a & b\\ b & c\end{pmatrix}\\ \begin{pmatrix}d & e\\ e & f\end{pmatrix}\end{matrix}\right\},$$

$$\begin{aligned}\mathsf{A}^r{+}\mathsf{A}^c{-}\mathsf{A} &= \left\{\begin{matrix}\begin{pmatrix}a & b\\ d & e\end{pmatrix}\\ \begin{pmatrix}b & c\\ e & f\end{pmatrix}\end{matrix}\right\} + \left\{\begin{matrix}\begin{pmatrix}a & d\\ b & e\end{pmatrix}\\ \begin{pmatrix}b & e\\ c & f\end{pmatrix}\end{matrix}\right\} - \left\{\begin{matrix}\begin{pmatrix}a & b\\ b & c\end{pmatrix}\\ \begin{pmatrix}d & e\\ e & f\end{pmatrix}\end{matrix}\right\} = \left\{\begin{matrix}\begin{pmatrix}a & d\\ d & 2e-c\end{pmatrix}\\ \begin{pmatrix}2b-d & c\\ c & f\end{pmatrix}\end{matrix}\right\}\\ &= 2\sigma^{-2}\left\{\begin{matrix}\begin{pmatrix}0 & 0\\ 0 & 0\end{pmatrix}\\ h_y\begin{pmatrix}h_{\mathrm{xx}} & h_{\mathrm{xy}}\\ h_{\mathrm{xy}} & h_{\mathrm{yy}}\end{pmatrix}\end{matrix}\right\} = 2\sigma^{-2}\left\{\begin{matrix}\mathbf{0}\\ h_y\nabla\nabla^T h\end{matrix}\right\}\end{aligned} \tag{S1.15}$$

Thus, substituting Eq. (S1.15) into Eq. (S1.9), we get distortion matrices as

$$\begin{aligned}\mathsf{Q} &= \frac{1}{2}\langle \mathbf{M}_{\mathbf{s}}(\mathbf{0};\hat{\mathbf{u}})^{-1}, \mathsf{A}^r{+}\mathsf{A}^c{-}\mathsf{A}\rangle\\ &= \frac{1}{2}\left\langle\begin{pmatrix}\sigma^2 & 0\\ 0 & \frac{\sigma^2}{1+h_y^2}\end{pmatrix}, 2\sigma^{-2}\left\{\begin{matrix}\mathbf{0}\\ h_y\nabla\nabla^T h\end{matrix}\right\}\right\rangle\\ &= \left\{\begin{matrix}\mathbf{0}\\ \dfrac{h_y}{1+h_y^2}\nabla\nabla^T h\end{matrix}\right\}.\end{aligned} \tag{S1.16}$$

Finally, we transform Eq. (S1.16) into

$$\mathsf{Q} = \left\{\begin{matrix}\mathbf{0}\\ \dfrac{|\nabla H(\hat{\mathbf{u}})|}{\eta^2+|\nabla H(\hat{\mathbf{u}})|^2}\mathbf{E}^T\nabla\nabla^T H(\hat{\mathbf{u}})\mathbf{E}\end{matrix}\right\} \tag{S1.17}$$

with

$$\nabla\nabla^T H(\mathbf{u}) = \begin{pmatrix}\frac{\partial^2 H(\mathbf{u})}{\partial X^2} & \frac{\partial^2 H(\mathbf{u})}{\partial X\partial Y}\\ \frac{\partial^2 H(\mathbf{u})}{\partial X\partial Y} & \frac{\partial^2 H(\mathbf{u})}{\partial Y^2}\end{pmatrix}. \tag{S1.18}$$

#### S1.3. Analysis with local coordinate normalization

We denote by $F(\mathbf{u}', \mathbf{u}^\circ)$ an invasion fitness function of mutant phenotype $\mathbf{u}'$ under existence of a monomorphic resident with phenotype $\mathbf{u}^\circ$. We express $F(\mathbf{u}', \mathbf{u}^\circ)$ in the rotated coordinates $\mathbf{s}$ centered at an arbitrary point $\hat{\mathbf{u}}$, as

$$f(\mathbf{s}', \mathbf{s}^\circ; \hat{\mathbf{u}}) = F(\mathbf{E}\mathbf{s}' + \hat{\mathbf{u}}, \mathbf{E}\mathbf{s}^\circ + \hat{\mathbf{u}}). \tag{S1.19}$$

Since $f(\mathbf{s}^\circ, \mathbf{s}^\circ; \hat{\mathbf{u}}) = 0$ holds for any $\mathbf{s}^\circ$, Eq. (S1.19) can be expanded as

$$f(\mathbf{s}', \mathbf{s}^\circ; \hat{\mathbf{u}}) = \mathbf{g}^T\boldsymbol{\delta}\mathbf{s} + \mathbf{s}^{\circ T}\mathbf{C}\boldsymbol{\delta}\mathbf{s} + \frac{1}{2}\boldsymbol{\delta}\mathbf{s}^T\mathbf{D}\boldsymbol{\delta}\mathbf{s} + \text{h.o.t.} \tag{S1.20}$$

with

$$\begin{aligned}\mathbf{g} &= \begin{pmatrix} g_x \\ g_y \end{pmatrix}, \\ \mathbf{C} &= \begin{pmatrix} C_{\mathrm{xx}} & C_{\mathrm{xy}} \\ C_{\mathrm{yx}} & C_{\mathrm{yy}} \end{pmatrix}, \\ \mathbf{D} &= \begin{pmatrix} D_{\mathrm{xx}} & D_{\mathrm{xy}} \\ D_{\mathrm{yx}} & D_{\mathrm{yy}} \end{pmatrix} \end{aligned} \tag{S1.21}$$

(Ito and Dieckmann, 2014; Ito and Sasaki, 2016). Substituting Eq. (S1.6) into Eq. (S1.20), we get the normalized invasion fitness function in the rotated-geodesic coordinates $\tilde{\mathbf{s}} = (\tilde{x}, \tilde{y})^T$.

$$\tilde{f}(\tilde{\mathbf{s}}', \tilde{\mathbf{s}}^\circ; \hat{\mathbf{u}}) = \tilde{\mathbf{g}}^T \boldsymbol{\delta}\tilde{\mathbf{s}} + \tilde{\mathbf{s}}^{\circ T} \tilde{\mathbf{C}} \boldsymbol{\delta}\tilde{\mathbf{s}} + \frac{1}{2} \boldsymbol{\delta}\tilde{\mathbf{s}}^T \tilde{\mathbf{D}} \boldsymbol{\delta}\tilde{\mathbf{s}} + \mathrm{h.o.t.} \tag{S1.22}$$

with

$$\begin{aligned}\tilde{\mathbf{g}} &= \mathbf{g}, \\ \tilde{\mathbf{C}} &= \mathbf{C} + \boldsymbol{\Omega}, \\ \tilde{\mathbf{D}} &= \mathbf{D} + \boldsymbol{\Omega}\end{aligned} \tag{S1.23}$$

where

$$\boldsymbol{\Omega} = -\langle \mathbf{g}^T, \mathrm{Q} \rangle = -\mathbf{g}_x \mathbf{Q}^x - \mathbf{g}_y \mathbf{Q}^y. \tag{S1.24}$$

In the rotated-geodesic coordinates $\tilde{\mathbf{s}}$, the mutation distribution can be locally approximated as a constant Gaussian mutation distribution with its covariance matrix given by $\mathbf{M}_{\mathbf{s}}(\mathbf{0}; \hat{\mathbf{u}})^{-1}$. Thus, existing conditions for evolutionary branching points, lines, and areas can be applied to $\tilde{f}(\tilde{\mathbf{s}}', \tilde{\mathbf{s}}^\circ; \hat{\mathbf{u}})$ as shown by Ito and Sasaki (2020).

#### S1.4. Analysis with Lagrange multiplier method

We show that the same invasion fitness function with the normalized invasion fitness function, given by Eq. (S1.22), can be derived by the Lagrange multiplier method (Ito and Sasaki, 2016).We form rotated hyper coordinates $\bar{\mathbf{s}} = (x, y, z)$ by rotating the coordinates of the hyper trait space $\bar{\mathbf{u}} = (X, Y, Z)^T$ as

$$\bar{\mathbf{s}} = \begin{pmatrix} x \\ y \\ z \end{pmatrix} = \begin{pmatrix} \mathbf{s} \\ Z \end{pmatrix} = \begin{pmatrix} \mathbf{E}^T[\mathbf{u} - \hat{\mathbf{u}}] \\ Z \end{pmatrix}. \tag{S1.25}$$

In the rotated hyper coordinates $\bar{\mathbf{s}}$, we define the invasion fitness function as

$$\begin{aligned}\bar{f}(\bar{\mathbf{s}}', \bar{\mathbf{s}}^\circ; \hat{\mathbf{u}}) &= f(\mathbf{s}', \mathbf{s}^\circ; \hat{\mathbf{u}}) \\ &= \bar{\mathbf{g}}^T \boldsymbol{\delta}\bar{\mathbf{s}} + \bar{\mathbf{s}}^{\circ T} \bar{\mathbf{C}} \boldsymbol{\delta}\bar{\mathbf{s}} + \frac{1}{2} \boldsymbol{\delta}\bar{\mathbf{s}}^T \bar{\mathbf{D}} \boldsymbol{\delta}\bar{\mathbf{s}} + \mathrm{h.o.t.}\end{aligned} \tag{S1.26}$$

with

$$\begin{aligned}\bar{\mathbf{g}} &= \begin{pmatrix} g_x \\ g_y \\ 0 \end{pmatrix}, \\ \bar{C} &= \begin{pmatrix} C_{\mathrm{xx}} & C_{\mathrm{xy}} & 0 \\ C_{\mathrm{yx}} & C_{\mathrm{yy}} & 0 \\ 0 & 0 & 0 \end{pmatrix}, \\ \bar{D} &= \begin{pmatrix} D_{\mathrm{xx}} & D_{\mathrm{xy}} & 0 \\ D_{\mathrm{yx}} & D_{\mathrm{yy}} & 0 \\ 0 & 0 & 0 \end{pmatrix}, \end{aligned} \tag{S1.27}$$

where $z$ has no contribution. In addition we define a hyper-constraint surface $z = h(\mathbf{s})$ as

$$\bar{h}(\bar{\mathbf{s}}) = h(\mathbf{s}) - z = 0. \tag{S1.28}$$

where

$$\begin{aligned}
\nabla\bar{h}(\bar{\mathbf{s}}) &= \begin{pmatrix} \nabla h(\mathbf{s}) \\ -1 \end{pmatrix}, \nabla\bar{h} = \nabla\bar{h}(\mathbf{0}) = \begin{pmatrix} \nabla h \\ -1 \end{pmatrix} = \begin{pmatrix} 0 \\ h_y \\ -1 \end{pmatrix}, \\
\nabla\nabla^T\bar{h}(\bar{\mathbf{s}}) &= \begin{pmatrix} \nabla\nabla^T h(\mathbf{s}) & 0 \\ 0 & 0 \end{pmatrix}, \nabla\nabla^T\bar{h} = \nabla\nabla^T\bar{h}(\mathbf{0}) = \begin{pmatrix} \nabla\nabla^T h & 0 \\ 0 & 0 \end{pmatrix} = \begin{pmatrix} h_{\mathrm{xx}} & h_{\mathrm{xy}} & 0 \\ h_{\mathrm{xy}} & h_{\mathrm{yy}} & 0 \\ 0 & 0 & 0 \end{pmatrix}
\end{aligned} \tag{S1.29}$$

Then the evolutionary dynamics in the rotated coordinates $\mathbf{s} = (x, y)^T$ can be translated into the evolutionary dynamics along the hyper-constraint surface $\bar{h}(\bar{\mathbf{s}})$ in the rotated hyper coordinates $\bar{\mathbf{s}}$ that has constant and isotropic mutational metric. Following Ito and Sasaki (2016), we project the invasion fitness function along the hyper-constraint surface on its tangent plane at $\mathbf{s} = (0, 0)^T$. We choose $\bar{\mathbf{e}}_1$ and $\bar{\mathbf{e}}_2$ in orthogonal directions to $\nabla\bar{h}(\bar{\mathbf{s}})$, and describe the tangent plane as

$$\begin{aligned}
\bar{\mathbf{s}}_E &= x\bar{\mathbf{e}}_1 + y\bar{\mathbf{e}}_2 = \begin{pmatrix} \bar{\mathbf{e}}_1 & \bar{\mathbf{e}}_2 \end{pmatrix}\begin{pmatrix} x \\ y \end{pmatrix} = \bar{\mathbf{E}}\mathbf{s}, \\
\bar{\mathbf{e}}_1 &= \begin{pmatrix} 1 \\ 0 \\ 0 \end{pmatrix}, \bar{\mathbf{e}}_2 = \begin{pmatrix} 0 \\ 1 \\ h_y \end{pmatrix}, \bar{\mathbf{E}} = \begin{pmatrix} \bar{\mathbf{e}}_1 & \bar{\mathbf{e}}_2 \end{pmatrix} = \begin{pmatrix} 1 & 0 \\ 0 & 1 \\ 0 & h_y \end{pmatrix}
\end{aligned} \tag{S1.30}$$

On this basis, we can define the mutational square distance at $\hat{\mathbf{u}}$ along the surface as

$$\begin{aligned}
[dl^2]_{\mathbf{u}=\hat{\mathbf{u}}} &= \sigma^{-2}\mathbf{d}\bar{\mathbf{s}}_E^T\mathbf{d}\bar{\mathbf{s}}_E \\
&= \sigma^{-2}\mathbf{ds}^T\bar{\mathbf{E}}^T\bar{\mathbf{E}}\mathbf{ds} \\
&= \sigma^{-2}\mathbf{ds}^T\begin{pmatrix} 1 & 0 & 0 \\ 0 & 1 & h_y \end{pmatrix}\begin{pmatrix} 1 & 0 \\ 0 & 1 \\ 0 & h_y \end{pmatrix}\mathbf{ds} = \sigma^{-2}\mathbf{ds}^T\begin{pmatrix} 1 & 0 \\ 0 & 1 + h_y^2 \end{pmatrix}\mathbf{ds}, \\
&= \mathbf{ds}^T\begin{pmatrix} \sigma^{-2} & 0 \\ 0 & \sigma^{-2}[1 + h_y^2] \end{pmatrix}\mathbf{ds},
\end{aligned} \tag{S1.31}$$

which gives the same mutation metric with $\mathbf{M}_{\mathbf{s}}(\mathbf{0}; \hat{\mathbf{u}})$ in Eqs. (S1.13).

According to Ito and Sasaki (2016), the projected invasion fitness function onto the tangent plane can be expressed as

$$\begin{aligned}
f_h(\mathbf{s}', \mathbf{s}^\circ; \hat{\mathbf{u}}) &= \bar{f}_h(\bar{\mathbf{s}}'_E, \bar{\mathbf{s}}^\circ_E; \hat{\mathbf{u}}) \\
&= \bar{f}(\bar{\mathbf{s}}'_E, \bar{\mathbf{s}}^\circ_E; \hat{\mathbf{u}}) - \lambda[\bar{h}(\bar{\mathbf{s}}'_E) - \bar{h}(\bar{\mathbf{s}}^\circ_E)] \\
&= \bar{\mathbf{g}}^T\boldsymbol{\delta}\bar{\mathbf{s}}_E + \bar{\mathbf{s}}_E^{\circ T}[\bar{\mathbf{C}} - \lambda\nabla\nabla^T\bar{h}]\boldsymbol{\delta}\bar{\mathbf{s}}_E + \frac{1}{2}\boldsymbol{\delta}\bar{\mathbf{s}}_E^T[\bar{\mathbf{D}} - \lambda\nabla\nabla^T\bar{h}]\boldsymbol{\delta}\bar{\mathbf{s}}_E + \text{h.o.t.} \\
&= \bar{\mathbf{g}}^T\bar{\mathbf{E}}\boldsymbol{\delta}\mathbf{s} + \mathbf{s}^{\circ T}\bar{\mathbf{E}}^T[\bar{\mathbf{C}} - \lambda\nabla\nabla^T\bar{h}]\bar{\mathbf{E}}\boldsymbol{\delta}\mathbf{s} + \frac{1}{2}\boldsymbol{\delta}\mathbf{s}^T\bar{\mathbf{E}}^T[\mathbf{D} - \lambda\nabla\nabla^T\bar{h}]\bar{\mathbf{E}}\boldsymbol{\delta}\mathbf{s} + \text{h.o.t.} \\
&= \mathbf{g}^T\boldsymbol{\delta}\mathbf{s} + \mathbf{s}^{\circ T}[\mathbf{C} + \boldsymbol{\Omega}_L]\boldsymbol{\delta}\mathbf{s} + \frac{1}{2}\boldsymbol{\delta}\mathbf{s}^T[\mathbf{D} + \boldsymbol{\Omega}_L]\boldsymbol{\delta}\mathbf{s} + \text{h.o.t.}
\end{aligned} \tag{S1.32}$$

with

$$\begin{aligned}
\boldsymbol{\Omega}_L &= -\lambda\bar{\mathbf{E}}^T\nabla\nabla^T\bar{h}\bar{\mathbf{E}}, \\
\lambda &= \frac{\bar{\mathbf{g}}^T\nabla\bar{h}}{|\nabla\bar{h}|^2}.
\end{aligned} \tag{S1.33}$$

We transform Eqs. (S1.33) further into

$$\begin{aligned}\boldsymbol{\Omega}_L &= -\frac{\begin{pmatrix} g_x & g_y & 0 \end{pmatrix}\begin{pmatrix} 0 \\ h_y \\ -1 \end{pmatrix}}{\left|\begin{pmatrix} 0 \\ h_y \\ -1 \end{pmatrix}\right|^2}\begin{pmatrix} 1 & 0 & 0 \\ 0 & 1 & h_y \end{pmatrix}\begin{pmatrix} h_{\mathrm{xx}} & h_{\mathrm{xy}} & 0 \\ h_{\mathrm{xy}} & h_{\mathrm{yy}} & 0 \\ 0 & 0 & 0 \end{pmatrix}\begin{pmatrix} 1 & 0 \\ 0 & 1 \\ 0 & h_y \end{pmatrix} \\ &= -\frac{g_y h_y}{1+h_y^2}\nabla\nabla^T h = -\left\langle \begin{pmatrix} g_x \\ g_y \end{pmatrix}^T, \left\{ \begin{matrix} \mathbf{0} \\ \frac{h_y}{1+|h_y|^2}\nabla\nabla^T h \end{matrix} \right\} \right\rangle \\ &= -\langle \mathbf{g}^T, \mathsf{Q}_L \rangle,\end{aligned} \tag{S1.34}$$

with

$$\mathsf{Q}_L = \left\{ \begin{matrix} \mathbf{0} \\ \frac{h_y}{1+|h_y|^2}\nabla\nabla^T h \end{matrix} \right\}. \tag{S1.35}$$

Note that Eq. (S1.32) is identical to the normalized invasion fitness function, Eq. (S1.22), because $\mathsf{Q}_L{=}\mathsf{Q}$ holds.

### S2. Appendix B: $N$-Dim trait spaces with $M$-constraints

We extend the analysis in the previous section for arbitrary $N$-dimensional trait spaces with arbitrary $M$ constraints satisfying $M \leqslant N$.

#### S2.1. Mutation mountain mapping

For an arbitrary trait space $\mathbf{u} = (X_1, ...., X_N)^T$, we assume an $(N+M)$-dimensional hyper space $\mathbf{u} = (X_1, ...., X_N, Z_1, ..., Z_M)^T$ and an $N$-dimensional hyper-constraint surface in it, defined by $\left(X_1, ..., X_N, \overset{1}{H}(\mathbf{u}), ..., \overset{M}{H}(\mathbf{u})\right)^T$. We also assume that the mutational metric for the hyper trait space is constant and isotropic, given by $\sigma^{-2}\mathbf{I}$. Then, the square distance $\mathrm{d}l^2$ along the hyper-constraint surface is expressed as

$$\begin{aligned}\mathrm{d}l^2 &= \sigma^{-2}[(\mathrm{d}X_1)^2 + ... + (\mathrm{d}X_N)^2 + (\mathrm{d}Z_1)^2 + \cdots + (\mathrm{d}Z_N)^2] \\ &= \sigma^{-2}\left[(\mathrm{d}X_1)^2 + ... + (\mathrm{d}X_N)^2 + \sum_{m=1}^{M}\left(\frac{\partial \overset{m}{H}(\mathbf{u})}{\partial X_1}\mathrm{d}X_1 + ... + \frac{\partial \overset{m}{H}(\mathbf{u})}{\partial X_N}\mathrm{d}X_N\right)^2\right] \\ &= \sigma^{-2}\begin{pmatrix} \mathrm{d}X_1 \\ \vdots \\ \mathrm{d}X_N \end{pmatrix}^T\left[\mathbf{I} + \sum_{m=1}^{M}\begin{pmatrix} \frac{\partial \overset{m}{H}(\mathbf{u})}{\partial X_1}\frac{\partial \overset{m}{H}(\mathbf{u})}{\partial X_1} & \cdots & \frac{\partial \overset{m}{H}(\mathbf{u})}{\partial X_1}\frac{\partial \overset{m}{H}(\mathbf{u})}{\partial X_N} \\ \vdots & \ddots & \vdots \\ \frac{\partial \overset{m}{H}(\mathbf{u})}{\partial X_N}\frac{\partial \overset{m}{H}(\mathbf{u})}{\partial X_1} & \cdots & \frac{\partial \overset{m}{H}(\mathbf{u})}{\partial X_N}\frac{\partial \overset{m}{H}(\mathbf{u})}{\partial X_N} \end{pmatrix}\right]\begin{pmatrix} \mathrm{d}X_1 \\ \vdots \\ \mathrm{d}X_N \end{pmatrix} \\ &= \sigma^{-2}\mathbf{du}^T\left[\mathbf{I} + \sum_{m=1}^{M}\nabla\overset{m}{H}(\mathbf{u})\nabla^T\overset{m}{H}(\mathbf{u})\right]\mathbf{du} \\ &= \mathbf{du}^T\mathbf{M}(\mathbf{u})\mathbf{du},\end{aligned} \tag{S2.1}$$

with infinitesimal $\mathbf{du}=(\mathrm{d}X_1,....,\mathrm{d}X_N)$, from which we get the mutational metric for the trait space,

$$\mathbf{M}(\mathbf{u}) \;=\; \sigma^{-2}\left[\mathbf{I}+\sum_{m=1}^{M}\nabla\overset{m}{H}(\mathbf{u})\nabla^T\overset{m}{H}(\mathbf{u})\right]. \tag{S2.2}$$

For efficient derivation of distortion matrices, we introduce coordinates $\mathbf{s}=(x_1,...,x_N)^T$,

$$\mathbf{u}-\hat{\mathbf{u}} \;=\; \mathbf{s} \tag{S2.3}$$

Then the mutation mountains $\overset{1}{H}(\mathbf{u}),...,\overset{M}{H}(\mathbf{u})$ are expressed in the rotated coordinates as

$$\overset{m}{h}(\mathbf{s}) \;=\; \overset{m}{H}(\mathbf{s}+\hat{\mathbf{u}}), \tag{S2.4}$$

for $m=1,...,M$, with

$$\nabla\overset{m}{h}(\mathbf{s}) \;=\; \begin{pmatrix}\overset{m}{h}_1(\mathbf{s})\\ \vdots \\ \overset{m}{h}_N(\mathbf{s})\end{pmatrix}, \nabla\overset{m}{h}(\mathbf{0})=\begin{pmatrix}\overset{m}{h}_1\\ \vdots \\ \overset{m}{h}_N\end{pmatrix}. \tag{S2.5}$$

The mutational metric in the rotated coordinates is expressed as

$$\begin{aligned}
\mathbf{M}_{\mathbf{s}}(\mathbf{s};\hat{\mathbf{u}}) \;&=\; \sigma^{-2}\left[\mathbf{I}+\sum_{m=1}^{M}\nabla\overset{m}{h}(\mathbf{s})\nabla^T\overset{m}{h}(\mathbf{s})\right]\\
&=\; \sigma^{-2}\left[\mathbf{I}+\begin{pmatrix}\nabla\overset{1}{h}(\mathbf{s}) & \cdots & \nabla\overset{M}{h}(\mathbf{s})\end{pmatrix}\begin{pmatrix}\nabla^T\overset{1}{h}(\mathbf{s})\\ \vdots \\ \nabla^T\overset{M}{h}(\mathbf{s})\end{pmatrix}\right],\\
\mathbf{M}_{\mathbf{s}}(\mathbf{0};\hat{\mathbf{u}}) \;&=\; \sigma^{-2}[\mathbf{I}+\mathbf{B}\mathbf{B}^T],\\
\mathbf{B} \;&=\; \begin{pmatrix}\nabla\overset{1}{h} & \cdots & \nabla\overset{M}{h}\end{pmatrix}=\nabla\mathbf{h}^T.
\end{aligned} \tag{S2.6}$$

and

$$M_{\mathbf{s}ij}(\mathbf{s}) \;=\; \sigma^{-2}\left[\delta(i-j)+\sum_{m=1}^{M}\overset{m}{h}_i(\mathbf{s})\overset{m}{h}_j(\mathbf{s})\right], \tag{S2.7}$$

which gives

$$\mathbf{M}_{\mathbf{s}}(\mathbf{0};\hat{\mathbf{u}})^{-1} \;=\; \sigma^2[\mathbf{I}+\mathbf{B}\mathbf{B}^T]^{-1}. \tag{S2.8}$$

#### S2.2. Distortion matrices

We see

$$\sigma^2 A_{ij}^k \;=\; \sigma^2\left[\frac{\partial M_{\mathbf{s}ij}(\mathbf{s})}{\partial x_k}\right]_{\mathbf{s}=\mathbf{0}}=\left[\frac{\partial}{\partial x_k}\sum_{m=1}^{M}\overset{m}{h}_i(\mathbf{s})\overset{m}{h}_j(\mathbf{s})\right]_{\mathbf{s}=\mathbf{0}}=\sum_{m=1}^{M}\left[\overset{m}{h}_i\overset{m}{h}_{jk}+\overset{m}{h}_j\overset{m}{h}_{ik}\right] \tag{S2.9}$$

and

$$\begin{aligned}
\sigma^2[A_{ij}^k]^r \;&=\; \sigma^2 A_{kj}^i=\sum_{m=1}^{M}\left[\overset{m}{h}_k\overset{m}{h}_{ji}+\overset{m}{h}_j\overset{m}{h}_{ki}\right]\\
\sigma^2[A_{ij}^k]^c \;&=\; \sigma^2 A_{ik}^j=\sum_{m=1}^{M}\left[\overset{m}{h}_i\overset{m}{h}_{kj}+\overset{m}{h}_k\overset{m}{h}_{ij}\right].
\end{aligned} \tag{S2.10}$$

Thus, from

$$
\begin{aligned}
[A^k_{ij}]^r + [A^k_{ij}]^c - A^k_{ij} &= 2\sigma^{-2} \sum_{m=1}^{M} \overset{m}{h}_{k} \overset{m}{h}_{ji}, \\
\mathsf{A}^r + \mathsf{A}^c - \mathsf{A} &= 2\sigma^{-2} \left\{ \begin{array}{c} \sum_{m=1}^{M} \overset{m}{h}_{1} \nabla\nabla^T \overset{m}{h} \\ \vdots \\ \sum_{m=1}^{M} \overset{m}{h}_{N} \nabla\nabla^T \overset{m}{h} \end{array} \right\}, \\
&= 2\sigma^{-2} \left\langle \begin{pmatrix} \overset{1}{h}_{1} & \cdots & \overset{M}{h}_{1} \\ \vdots & \ddots & \vdots \\ \overset{1}{h}_{N} & \cdots & \overset{M}{h}_{N} \end{pmatrix}, \left\{ \begin{array}{c} \nabla\nabla^T \overset{1}{h} \\ \vdots \\ \nabla\nabla^T \overset{M}{h} \end{array} \right\} \right\rangle \\
&= 2\sigma^{-2} \left\langle \begin{pmatrix} \nabla \overset{1}{h}_{1} & \cdots & \nabla \overset{M}{h}_{1} \end{pmatrix}, \nabla\nabla^T \mathbf{h} \right\rangle \\
&= 2\sigma^{-2} \langle \mathbf{B}, \nabla\nabla^T \mathbf{h} \rangle \qquad \text{(S2.11)}
\end{aligned}
$$

with $\nabla\nabla^T \overset{m}{h} = \left[ \nabla\nabla^T \overset{m}{h} \right]_{\mathbf{s}=\mathbf{0}}$, we get distortion matrices as

$$
\begin{aligned}
\mathsf{Q} &= \frac{1}{2} \langle \mathbf{M}_{\mathbf{s}}(\mathbf{0}; \hat{\mathbf{u}})^{-1}, \mathsf{A}^r + \mathsf{A}^c - \mathsf{A} \rangle \\
&= \frac{1}{2} \langle \sigma^2 [\mathbf{I}_N + \mathbf{B}\mathbf{B}^T]^{-1}, 2\sigma^{-2} \langle \mathbf{B}, \nabla\nabla^T \mathbf{h} \rangle \rangle \\
&= \langle [\mathbf{I}_N + \mathbf{B}\mathbf{B}^T]^{-1}, \langle \mathbf{B}, \nabla\nabla^T \mathbf{h} \rangle \rangle \\
&= \langle [\mathbf{I}_N + \mathbf{B}\mathbf{B}^T]^{-1} \mathbf{B}, \nabla\nabla^T \mathbf{h} \rangle \qquad \text{(S2.12)}
\end{aligned}
$$

(see Appendix C for operations for matrix-vectors).

#### S2.3. Analysis with local coordinate normalization

We derive the normalized invasion fitness function in the same manner as in the two-dimensional case in Section 2.3. Specifically, for an invasion fitness function in the rotated coordinates,

$$
f(\mathbf{s}', \mathbf{s}^\circ; \hat{\mathbf{u}}) = \mathbf{g}^T \boldsymbol{\delta}\mathbf{s} + \mathbf{s}^{\circ T} \mathbf{C} \boldsymbol{\delta}\mathbf{s} + \frac{1}{2} \boldsymbol{\delta}\mathbf{s}^T \mathbf{D} \boldsymbol{\delta}\mathbf{s} + \text{h.o.t.} \qquad \text{(S2.13)}
$$

with

$$
\begin{aligned}
\mathbf{g} &= \begin{pmatrix} g_1 \\ \vdots \\ g_N \end{pmatrix}, \\
\mathbf{C} &= \begin{pmatrix} C_{11} & \cdots & C_{1N} \\ \vdots & \ddots & \vdots \\ C_{N1} & \cdots & C_{NN} \end{pmatrix}, \\
\mathbf{D} &= \begin{pmatrix} D_{11} & \cdots & D_{1N} \\ \vdots & \ddots & \vdots \\ D_{N1} & \cdots & D_{NN} \end{pmatrix}, \qquad \text{(S2.14)}
\end{aligned}
$$

we get the normalized invasion fitness function in the rotated-geodesic coordinates $\tilde{\mathbf{s}} = (\tilde{x}_1, ..., \tilde{x}_N)^T$,

$$
\tilde{f}(\tilde{\mathbf{s}}', \tilde{\mathbf{s}}^\circ; \hat{\mathbf{u}}) = \tilde{\mathbf{g}}^T \boldsymbol{\delta}\tilde{\mathbf{s}} + \tilde{\mathbf{s}}^{\circ T} \tilde{\mathbf{C}} \boldsymbol{\delta}\tilde{\mathbf{s}} + \frac{1}{2} \boldsymbol{\delta}\tilde{\mathbf{s}}^T \tilde{\mathbf{D}} \boldsymbol{\delta}\tilde{\mathbf{s}} + \text{h.o.t.} \qquad \text{(S2.15)}
$$

with

$$\begin{aligned}\tilde{\mathbf{g}} &= \mathbf{g},\\ \tilde{\mathbf{C}} &= \mathbf{C}-\langle\mathbf{g}^T,\mathsf{Q}\rangle,\\ \tilde{\mathbf{D}} &= \mathbf{D}-\langle\mathbf{g}^T,\mathsf{Q}\rangle,\end{aligned} \tag{S2.16}$$

where

$$\mathbf{s} = \tilde{\mathbf{s}}-\frac{1}{2}\tilde{\mathbf{s}}\mathsf{Q}\tilde{\mathbf{s}}. \tag{S2.17}$$

#### S2.4. Analysis with Lagrange multiplier method

We show that the same invasion fitness function with the normalized invasion fitness function, given by Eqs. (S2.15), can be derived by the Lagrange multiplier method (Ito and Sasaki, 2016). We form the rotated hyper coordinates $\bar{\mathbf{s}}=(x_1,...,x_N,z_1,...,z_M)$ by coordinate rotation of the hyper trait space $\bar{u}=(X_1,...,X_N,Z_1,...,Z_M)^T$,

$$\bar{\mathbf{s}} = \begin{pmatrix} x_1\\ \vdots\\ x_N\\ z_1\\ \vdots\\ z_M\end{pmatrix}=\begin{pmatrix}\mathbf{s}\\ \mathbf{z}\end{pmatrix}=\begin{pmatrix}[\mathbf{u}-\hat{\mathbf{u}}]\\ Z_1\\ \vdots\\ Z_M\end{pmatrix}. \tag{S2.18}$$

In the rotated hyper coordinates $\bar{\mathbf{s}}$, we express the invasion fitness function as

$$\begin{aligned}\bar{f}(\bar{\mathbf{s}}',\bar{\mathbf{s}}^\circ;\hat{\mathbf{u}}) &= f(\mathbf{s}',\mathbf{s}^\circ;\hat{\mathbf{u}})\\ &= \bar{\mathbf{g}}^T\boldsymbol{\delta}\bar{\mathbf{s}}+\bar{\mathbf{s}}^{\circ T}\bar{\mathbf{C}}\boldsymbol{\delta}\bar{\mathbf{s}}+\frac{1}{2}\boldsymbol{\delta}\bar{\mathbf{s}}^T\bar{\mathbf{D}}\boldsymbol{\delta}\bar{\mathbf{s}}+\text{h.o.t.}\end{aligned} \tag{S2.19}$$

with

$$\begin{aligned}\bar{\mathbf{g}} &= \begin{pmatrix}\mathbf{g}\\ \mathbf{0}\end{pmatrix},\\ \bar{\mathbf{C}} &= \begin{pmatrix}\mathbf{C} & \mathbf{0}\\ \mathbf{0} & \mathbf{0}\end{pmatrix},\\ \bar{\mathbf{D}} &= \begin{pmatrix}\mathbf{D} & \mathbf{0}\\ \mathbf{0} & \mathbf{0}\end{pmatrix},\end{aligned} \tag{S2.20}$$

where $\mathbf{z}$ has no contribution. In addition, we express the hyper-constraint surface $z_m=\overset{m}{h}(\mathbf{s})$ as

$$\overset{m}{\bar{h}}(\bar{\mathbf{s}}) = \overset{m}{h}(\mathbf{s})-z_m=0. \tag{S2.21}$$

where

$$\begin{aligned}\nabla\overset{1}{\bar{h}}(\bar{\mathbf{s}}) &= \begin{pmatrix}\nabla\overset{1}{h}(\mathbf{s})\\ -1\\ \vdots\\ 0\end{pmatrix},\nabla\overset{m}{\bar{h}}(\bar{\mathbf{s}})=\begin{pmatrix}\nabla\overset{m}{h}(\mathbf{s})\\ -\delta(1-m)\\ \vdots\\ -\delta(M-m)\end{pmatrix},\\ \nabla\overset{1}{\bar{h}}(\mathbf{0}) &= \nabla\overset{1}{\bar{h}}=\begin{pmatrix}\nabla\overset{1}{h}\\ -1\\ \vdots\\ 0\end{pmatrix},\nabla\overset{m}{\bar{h}}(\mathbf{0})=\nabla\overset{m}{\bar{h}}=\begin{pmatrix}\nabla\overset{m}{h}\\ -\delta(1-m)\\ \vdots\\ -\delta(M-m)\end{pmatrix},\\ \bar{\mathbf{B}} &= \begin{pmatrix}\begin{pmatrix}\nabla\overset{1}{h} & \cdots & \nabla\overset{M}{h}\end{pmatrix}\\ -\mathbf{I}_M\end{pmatrix}=\begin{pmatrix}\mathbf{B}\\ -\mathbf{I}_M\end{pmatrix}.\end{aligned} \tag{S2.22}$$

Then the evolutionary dynamics in the rotated coordinates $\mathbf{s}$ can be translated into the evolutionary dynamics along the hyper-constraint surface in the rotated hyper coordinates $\bar{\mathbf{s}}$ that has a constant and isotropic mutation. Following Ito and Sasaki(2016), we project the invasion fitness function along the hyper-constraint surface on its tangent plane at $\mathbf{s}=\mathbf{0}$. We choose $\bar{\mathbf{e}}_1, ..., \bar{\mathbf{e}}_N$ in orthogonal directions to $\nabla\overset{1}{\bar{h}}, ..., \nabla\overset{m}{\bar{h}}$, and describe the tangent plane as

$$\bar{\mathbf{e}}_1 = \begin{pmatrix} 1 \\ \vdots \\ 0 \\ \overset{1}{h}_{1} \\ \vdots \\ \overset{M}{h}_{1} \end{pmatrix}, \bar{\mathbf{e}}_n = \begin{pmatrix} \delta(1-n) \\ \vdots \\ \delta(N-n) \\ \overset{1}{h}_{n} \\ \vdots \\ \overset{M}{h}_{n} \end{pmatrix},$$

$$\bar{\mathbf{E}} = \begin{pmatrix} \bar{\mathbf{e}}_1 & \cdots & \bar{\mathbf{e}}_N \end{pmatrix} = \begin{pmatrix} \mathbf{I}_N \\ \begin{pmatrix} \nabla^T\overset{1}{h} \\ \vdots \\ \nabla^T\overset{M}{h} \end{pmatrix} \end{pmatrix} = \begin{pmatrix} \mathbf{I}_N \\ \mathbf{B}^T \end{pmatrix}, \tag{S2.23}$$

where $\left[\nabla\overset{m}{\bar{h}}\right]^T \bar{\mathbf{e}}_n = 0$ holds good, because

$$\left[\nabla\overset{m}{\bar{h}}\right]^T \bar{\mathbf{e}}_n = \begin{pmatrix} \overset{m}{h}_{1} \\ \vdots \\ \overset{m}{h}_{N} \\ -\delta(1-m) \\ \vdots \\ -\delta(M-m) \end{pmatrix}^T \begin{pmatrix} \delta(1-n) \\ \vdots \\ \delta(N-n) \\ \overset{1}{h}_{n} \\ \vdots \\ \overset{M}{h}_{n} \end{pmatrix} = \overset{m}{\bar{h}}_{n} - \overset{m}{\bar{h}}_{n} = 0. \tag{S2.24}$$

Here, the tangent plane is given by

$$\bar{\mathbf{s}}_E = \bar{\mathbf{E}}\mathbf{s}, \tag{S2.25}$$

Since the mutational metric for the hyper trait space is assumed to be constant and isotropic, we can express the mutational square distance at $\hat{\mathbf{u}}$ along the surface as

$$\begin{aligned} [dl^2]_{\mathbf{u}=\hat{\mathbf{u}}} &= \sigma^{-2}\mathbf{d}\bar{\mathbf{s}}_E^T\mathbf{d}\bar{\mathbf{s}}_E \\ &= \sigma^{-2}\mathbf{ds}^T\bar{\mathbf{E}}^T\bar{\mathbf{E}}\mathbf{ds} \\ &= \sigma^{-2}\mathbf{ds}^T\begin{pmatrix} \mathbf{I}_N & \mathbf{B} \end{pmatrix}\begin{pmatrix} \mathbf{I}_N \\ \mathbf{B}^T \end{pmatrix}\mathbf{ds} = \sigma^{-2}\mathbf{ds}^T[\mathbf{I}_N + \mathbf{B}\mathbf{B}^T]\mathbf{ds}, \end{aligned} \tag{S2.26}$$

which gives the same mutation metric with $\mathbf{M}_{\mathbf{s}}(\mathbf{0};\hat{\mathbf{u}})$ in Eq. (S2.6).

According to Ito and Sasaki(2016), the projected invasion fitness function is expressed as

$$\begin{aligned} f_h(\mathbf{s}', \mathbf{s}^\circ; \hat{\mathbf{u}}) &= \bar{f}_h(\bar{\mathbf{s}}'_E, \bar{\mathbf{s}}^\circ_E; \hat{\mathbf{u}}) \\ &= \bar{f}(\bar{\mathbf{s}}'_E, \bar{\mathbf{s}}^\circ_E; \hat{\mathbf{u}}) - \sum_{m=1}^{M} \lambda_m \left[\overset{m}{\bar{h}}(\bar{\mathbf{s}}'_E) - \overset{m}{\bar{h}}(\bar{\mathbf{s}}^\circ_E)\right] \\ &= \mathbf{g}^T\bar{\mathbf{E}}\boldsymbol{\delta}\mathbf{s} + \mathbf{s}^{\circ T}\bar{\mathbf{E}}^T\left[\bar{\mathbf{C}} - \sum_{m=1}^{M} \lambda_m \nabla\nabla^T\overset{m}{\bar{h}}\right]\bar{\mathbf{E}}\boldsymbol{\delta}\mathbf{s} \\ &\quad + \frac{1}{2}\boldsymbol{\delta}\mathbf{s}^T\bar{\mathbf{E}}^T\left[\bar{\mathbf{D}} - \sum_{m=1}^{M} \lambda_m \nabla\nabla^T\overset{m}{\bar{h}}\right]\bar{\mathbf{E}}\boldsymbol{\delta}\mathbf{s} + \text{h.o.t.} \\ &= \mathbf{g}^T\boldsymbol{\delta}\mathbf{s} + \mathbf{s}^{\circ T}[\mathbf{C} + \boldsymbol{\Omega}_L]\boldsymbol{\delta}\mathbf{s} + \frac{1}{2}\boldsymbol{\delta}\mathbf{s}^T[\mathbf{D} + \boldsymbol{\Omega}_L]\boldsymbol{\delta}\mathbf{s} + \text{h.o.t.} \end{aligned} \tag{S2.27}$$

with

$$
\begin{aligned}
-\mathbf{\Omega}_L &= \sum_{m=1}^{M} \lambda_m \bar{\mathbf{E}}^T \nabla\nabla^T \overset{m}{\bar{h}}\, \bar{\mathbf{E}} = \left\langle \boldsymbol{\lambda}^T, \bar{\mathbf{E}}^T \left\{ \begin{array}{c} \nabla\nabla^T \overset{1}{\bar{h}} \\ \vdots \\ \nabla\nabla^T \overset{M}{\bar{h}} \end{array} \right\} \bar{\mathbf{E}} \right\rangle, \\
\boldsymbol{\lambda}^T &= \bar{\mathbf{g}}^T \bar{\mathbf{B}}^{+T}, \\
\bar{\mathbf{B}}^+ &= [\bar{\mathbf{B}}^T \bar{\mathbf{B}}]^{-1} \bar{\mathbf{B}}^T.
\end{aligned} \tag{S2.28}
$$

Exploiting Eq. (S2.22), we see

$$
\begin{aligned}
\bar{\mathbf{B}}^+ &= \left[ \begin{pmatrix} \mathbf{B}^T & -\mathbf{I}_M \end{pmatrix} \begin{pmatrix} \mathbf{B} \\ -\mathbf{I}_M \end{pmatrix} \right]^{-1} \begin{pmatrix} \mathbf{B}^T & -\mathbf{I}_M \end{pmatrix} = [\mathbf{B}^T\mathbf{B} + \mathbf{I}_M]^{-1} \begin{pmatrix} \mathbf{B}^T & -\mathbf{I}_M \end{pmatrix}, \\
\boldsymbol{\lambda}^T &= \bar{\mathbf{g}}^T \bar{\mathbf{B}}^{+T} = \begin{pmatrix} \mathbf{g}^T & \mathbf{0}_M \end{pmatrix} \begin{pmatrix} \mathbf{B} \\ -\mathbf{I}_M \end{pmatrix} [\mathbf{B}^T\mathbf{B} + \mathbf{I}_M]^{-1} = \mathbf{g}^T \mathbf{B} [\mathbf{B}^T\mathbf{B} + \mathbf{I}_M]^{-1},
\end{aligned} \tag{S2.29}
$$

and thus

$$
\begin{aligned}
-\mathbf{\Omega} &= \left\langle \boldsymbol{\lambda}^T, \bar{\mathbf{E}}^T \left\{ \begin{array}{c} \nabla\nabla^T \overset{1}{\bar{h}} \\ \vdots \\ \nabla\nabla^T \overset{M}{\bar{h}} \end{array} \right\} \bar{\mathbf{E}} \right\rangle = \left\langle \boldsymbol{\lambda}^T, \begin{pmatrix} \mathbf{I}_N & \mathbf{B} \end{pmatrix} \left\{ \begin{array}{c} \begin{pmatrix} \nabla\nabla^T \overset{1}{h} & \mathbf{0} \\ \mathbf{0} & \mathbf{0} \end{pmatrix} \\ \vdots \\ \begin{pmatrix} \nabla\nabla^T \overset{M}{h} & \mathbf{0} \\ \mathbf{0} & \mathbf{0} \end{pmatrix} \end{array} \right\} \begin{pmatrix} \mathbf{I}_N \\ \mathbf{B}^T \end{pmatrix} \right\rangle \\
&= \left\langle \boldsymbol{\lambda}^T, \left\{ \begin{array}{c} \nabla\nabla^T \overset{1}{h} \\ \vdots \\ \nabla\nabla^T \overset{M}{h} \end{array} \right\} \right\rangle = \left\langle \mathbf{g}^T \mathbf{B} [\mathbf{B}^T\mathbf{B} + \mathbf{I}_M]^{-1}, \left\{ \begin{array}{c} \nabla\nabla^T \overset{1}{h} \\ \vdots \\ \nabla\nabla^T \overset{M}{h} \end{array} \right\} \right\rangle \\
&= \langle \mathbf{g}^T \mathbf{B} [\mathbf{B}^T\mathbf{B} + \mathbf{I}_M]^{-1}, \nabla\nabla^T \mathbf{h} \rangle \\
&= \langle \mathbf{g}^T, \langle \mathbf{B} [\mathbf{B}^T\mathbf{B} + \mathbf{I}_M]^{-1}, \nabla\nabla^T \mathbf{h} \rangle \rangle \\
&= \langle \mathbf{g}^T, \mathsf{Q}_L \rangle
\end{aligned} \tag{S2.30}
$$

where

$$
\mathsf{Q}_L = \langle \mathbf{B} [\mathbf{B}^T\mathbf{B} + \mathbf{I}_M]^{-1}, \nabla\nabla^T \mathbf{h} \rangle. \tag{S2.31}
$$

From $\mathbf{B}[\mathbf{B}^T\mathbf{B} + \mathbf{I}_M] = \mathbf{B}\mathbf{B}^T\mathbf{B} + \mathbf{B} = [\mathbf{I}_N + \mathbf{B}\mathbf{B}^T]\mathbf{B}$, we see that

$$
\mathbf{B}[\mathbf{B}^T\mathbf{B} + \mathbf{I}_M]^{-1} = [\mathbf{I}_N + \mathbf{B}\mathbf{B}^T]^{-1}\mathbf{B}. \tag{S2.32}
$$

Substituting Eq. (S2.32) into Eq. (S2.31), we find

$$
\begin{aligned}
\mathsf{Q}_L &= \langle [\mathbf{I}_N + \mathbf{B}\mathbf{B}^T]^{-1}\mathbf{B}, \nabla\nabla^T \mathbf{h} \rangle \\
&= \mathsf{Q}.
\end{aligned} \tag{S2.33}
$$

Therefore Eq. (S2.27) is identical to the normalized invasion fitness function, Eq. (S2.15), because $\mathsf{Q}_L$=$\mathsf{Q}$ holds.

#### S2.5. Coordinate transformation corresponding to Lagrange multiplier method

As described in Lemma 2 in Ito and Sasaki(2016), the above Lagrange multiplier method corresponds to a coordinate transformation of a point on the constraint surface, denoted by

$$\bar{\mathbf{s}}_h \;=\; (x_1,...,x_N,h_1(\mathbf{s}),...,h_M(\mathbf{s}))^T = \begin{pmatrix} \mathbf{s} \\ \mathbf{h}(\mathbf{s}) \end{pmatrix}, \tag{S2.34}$$

into a point on the tangent plane, denoted by

$$\bar{\mathbf{s}}_E \;=\; \bar{\mathbf{E}}\tilde{\mathbf{s}}, \tag{S2.35}$$

where the transformation is given by

$$\bar{\mathbf{s}}_h \;=\; \bar{\mathbf{s}}_E - \bar{\mathbf{B}}^{+T}\frac{1}{2}[\bar{\mathbf{s}}_E]^T \left\{ \begin{matrix} \nabla\nabla^T \bar{h}^{1} \\ \vdots \\ \nabla\nabla^T \bar{h}^{M} \end{matrix} \right\} [\bar{\mathbf{s}}_E] + h.o.t.. \tag{S2.36}$$

Substituting Eq. (3.23) $\bar{\mathbf{E}} = \begin{pmatrix} \mathbf{I}_N \\ \mathbf{B}^T \end{pmatrix}$, Eq. (3.29) $\bar{\mathbf{B}}^+ = [\mathbf{B}^T\mathbf{B} + \mathbf{I}_M]^{-1}(\ \mathbf{B}^T\ \ -\mathbf{I}_M\ )$, and Eqs. (3.34), (3.35) into Eq. (3.36) gives

$$\begin{pmatrix} \mathbf{s} \\ \mathbf{h}(\mathbf{s}) \end{pmatrix} = \begin{pmatrix} \tilde{\mathbf{s}} \\ \mathbf{B}^T\tilde{\mathbf{s}} \end{pmatrix} - \frac{1}{2}\begin{pmatrix} \mathbf{B} \\ -\mathbf{I}_M \end{pmatrix}[\mathbf{B}^T\mathbf{B} + \mathbf{I}_M]^{-1}\begin{pmatrix} \tilde{\mathbf{s}} \\ \mathbf{B}^T\tilde{\mathbf{s}} \end{pmatrix}^T \left\{ \begin{matrix} \nabla\nabla^T \bar{h}^{1} \\ \vdots \\ \nabla\nabla^T \bar{h}^{M} \end{matrix} \right\} \begin{pmatrix} \tilde{\mathbf{s}} \\ \mathbf{B}^T\tilde{\mathbf{s}} \end{pmatrix}.$$

Multiplying this equation by $(\ \mathbf{I}_N\ \ \mathbf{0}_M\ )$ from the left-side gives

$$\begin{aligned}
\mathbf{s} \;=&\; (\ \mathbf{I}_N\ \ \mathbf{0}_M\ )\begin{pmatrix} \tilde{\mathbf{s}} \\ \mathbf{B}^T\tilde{\mathbf{s}} \end{pmatrix} \\
&- \frac{1}{2}(\ \mathbf{I}_N\ \ \mathbf{0}_M\ )\begin{pmatrix} \mathbf{B} \\ -\mathbf{I}_M \end{pmatrix}[\mathbf{B}^T\mathbf{B} + \mathbf{I}_M]^{-1}\begin{pmatrix} \tilde{\mathbf{s}} \\ \mathbf{B}^T\tilde{\mathbf{s}} \end{pmatrix}^T \left\{ \begin{matrix} \nabla\nabla^T \bar{h}^{1} \\ \vdots \\ \nabla\nabla^T \bar{h}^{M} \end{matrix} \right\} \begin{pmatrix} \tilde{\mathbf{s}} \\ \mathbf{B}^T\tilde{\mathbf{s}} \end{pmatrix} \\
=&\; \tilde{\mathbf{s}} - \frac{1}{2}\mathbf{B}[\mathbf{B}^T\mathbf{B} + \mathbf{I}_M]^{-1}\begin{pmatrix} \tilde{\mathbf{s}} \\ \mathbf{B}^T\tilde{\mathbf{s}} \end{pmatrix}^T \left\{ \begin{matrix} \nabla\nabla^T \bar{h}^{1} \\ \vdots \\ \nabla\nabla^T \bar{h}^{M} \end{matrix} \right\} \begin{pmatrix} \tilde{\mathbf{s}} \\ \mathbf{B}^T\tilde{\mathbf{s}} \end{pmatrix} \\
=&\; \tilde{\mathbf{s}} - \frac{1}{2}\mathbf{B}[\mathbf{B}^T\mathbf{B} + \mathbf{I}_M]^{-1}\begin{pmatrix} \tilde{\mathbf{s}} \\ \mathbf{B}^T\tilde{\mathbf{s}} \end{pmatrix}^T \left\{ \begin{matrix} \begin{pmatrix} \nabla\nabla^T h^{1} & \mathbf{0} \\ \mathbf{0} & \mathbf{0} \end{pmatrix} \\ \vdots \\ \begin{pmatrix} \nabla\nabla^T h^{M} & \mathbf{0} \\ \mathbf{0} & \mathbf{0} \end{pmatrix} \end{matrix} \right\} \begin{pmatrix} \tilde{\mathbf{s}} \\ \mathbf{B}^T\tilde{\mathbf{s}} \end{pmatrix} \\
=&\; \tilde{\mathbf{s}} - \frac{1}{2}\mathbf{B}[\mathbf{B}^T\mathbf{B} + \mathbf{I}_M]^{-1}\tilde{\mathbf{s}}^T \left\{ \begin{matrix} \nabla\nabla^T h^{1} \\ \vdots \\ \nabla\nabla^T h^{M} \end{matrix} \right\} \tilde{\mathbf{s}} \\
=&\; \tilde{\mathbf{s}} - \frac{1}{2}\tilde{\mathbf{s}}^T \langle \mathbf{B}[\mathbf{B}^T\mathbf{B} + \mathbf{I}_M]^{-1}, \nabla\nabla^T\mathbf{h}\rangle \tilde{\mathbf{s}}.{\circ}
\end{aligned} \tag{S2.37}$$

Then by using Eqs. (S2.31) and (S2.33), we transform Eq. (S2.37) as

$$\begin{aligned}\mathbf{s} &= \tilde{\mathbf{s}} - \frac{1}{2}\tilde{\mathbf{s}}^T\langle \mathbf{B}[\mathbf{B}^T\mathbf{B}+\mathbf{I}_M]^{-1}, \nabla\nabla^T\mathbf{h}\rangle\tilde{\mathbf{s}} \\ &= \tilde{\mathbf{s}} - \frac{1}{2}\tilde{\mathbf{s}}^T\mathsf{Q}_L\tilde{\mathbf{s}} \\ &= \tilde{\mathbf{s}} - \frac{1}{2}\tilde{\mathbf{s}}^T\mathsf{Q}\tilde{\mathbf{s}},\end{aligned} \tag{S2.38}$$

which is identical to Eq. (S2.17).

### S3. Appendix C: Operations for vectors of matrices

#### S3.1. Structure

For efficient handling of nonlinear coordinate transformations, we introduce vectors having matrices as their components, referred to as "matrix-vectors", and some operations for them. We consider an arbitrary $N$-dimensional space $\mathbf{u}=(u_1, ..., u_N)^T$. We write a matrix-vector in the Sans-serif font $\mathsf{Q}$, and express its component matrices as

$$\mathsf{Q} = \begin{Bmatrix} \mathbf{Q}^1 \\ \vdots \\ \mathbf{Q}^N \end{Bmatrix}, \tag{S3.1}$$

with braces $\{\}$, where its $i$th component is given by an $N \times N$ matrix

$$\mathbf{Q}^i = \begin{pmatrix} Q^i_{11} & \cdots & Q^i_{1N} \\ \vdots & \ddots & \vdots \\ Q^i_{N1} & \cdots & Q^i_{NN} \end{pmatrix}. \tag{S3.2}$$

#### S3.2. Multiplication

We define multiplications of $\mathsf{Q}$ by a vector $\mathbf{u}=(u_1, ..., u_N)^T$ and by an $N \times N$ matrix $\mathbf{B}$ as

$$\mathbf{u}^T\mathsf{Q} = \begin{Bmatrix} \mathbf{u}^T\mathbf{Q}^1 \\ \vdots \\ \mathbf{u}^T\mathbf{Q}^N \end{Bmatrix}, \tag{S3.3}$$

$$\mathsf{Q}\mathbf{u} = \begin{Bmatrix} \mathbf{Q}^1\mathbf{u} \\ \vdots \\ \mathbf{Q}^N\mathbf{u} \end{Bmatrix}, \tag{S3.4}$$

$$\mathbf{B}\mathsf{Q} = \begin{Bmatrix} \mathbf{B}\mathbf{Q}^1 \\ \vdots \\ \mathbf{B}\mathbf{Q}^N \end{Bmatrix}, \tag{S3.5}$$

$$\mathsf{Q}\mathbf{B} = \begin{Bmatrix} \mathbf{Q}^1\mathbf{B} \\ \vdots \\ \mathbf{Q}^N\mathbf{B} \end{Bmatrix}. \tag{S3.6}$$

Here, $\mathbf{Q}^i$ and $\mathbf{B}$ are not necessarily square matrices, as long as their matrix-level multiplications in above formulas are valid. When a column matrix-vector has component matrices of dimension $1 \times N$ (or $1 \times 1$), it is treated as a normal matrix (or normal column vector), and a analogous rule is applied for a row matrix-vector. Specifically, the following rule is applied for arbitrary $M$ and $L$,

$$\begin{aligned}
\left\{\begin{matrix} \left( Q^1_{11} \cdots Q^1_{1L} \right) \\ \vdots \\ \left( Q^M_{11} \cdots Q^M_{1L} \right) \end{matrix}\right\} &= \begin{pmatrix} Q^1_{11} & \cdots & Q^1_{1L} \\ \vdots & \ddots & \vdots \\ Q^M_{11} & \cdots & Q^M_{1L} \end{pmatrix}, \\
\left\{\begin{matrix} \left( Q^1_{11} \right) \\ \vdots \\ \left( Q^M_{11} \right) \end{matrix}\right\} &= \begin{pmatrix} Q^1_{11} \\ \vdots \\ Q^M_{11} \end{pmatrix}, \\
\left\{ \begin{pmatrix} Q^1_{11} \\ \vdots \\ Q^1_{L1} \end{pmatrix} \cdots \begin{pmatrix} Q^M_{11} \\ \vdots \\ Q^M_{L1} \end{pmatrix} \right\} &= \begin{pmatrix} Q^1_{11} & \cdots & Q^M_{11} \\ \vdots & \ddots & \vdots \\ Q^1_{L1} & \cdots & Q^M_{L1} \end{pmatrix}, \\
\left\{ \left( Q^1_{11} \right) \cdots \left( Q^M_{11} \right) \right\} &= \left( Q^1_{11} \cdots Q^M_{11} \right).
\end{aligned} \tag{S3.7}$$

For example, Eq. (S3.3) is treated as an $N \times N$ matrix because each component $\mathbf{u}^T\mathbf{Q}^i$ becomes row vector of length $N$, i.e.,

$$\begin{aligned}
\mathbf{u}^T \left\{\begin{matrix} \mathbf{Q}^1 \\ \vdots \\ \mathbf{Q}^N \end{matrix}\right\} &= \left\{\begin{matrix} \mathbf{u}^T\mathbf{Q}^1 \\ \vdots \\ \mathbf{u}^T\mathbf{Q}^N \end{matrix}\right\} = \begin{pmatrix} \mathbf{u}^T\mathbf{Q}^1 \\ \vdots \\ \mathbf{u}^T\mathbf{Q}^N \end{pmatrix} \\
&= \begin{pmatrix} \sum_j u_j Q^1_{j1} & \cdots & \sum_j u_j Q^1_{jN} \\ \vdots & \ddots & \vdots \\ \sum_j u_j Q^N_{j1} & \cdots & \sum_j u_j Q^N_{jN} \end{pmatrix},
\end{aligned} \tag{S3.8}$$

while Eq. (5.4) gives a matrix-vector consisting of column vectors of length $N$,

$$\left\{\begin{matrix} \mathbf{Q}^1 \\ \vdots \\ \mathbf{Q}_N \end{matrix}\right\} \mathbf{u} = \left\{\begin{matrix} \mathbf{Q}^1\mathbf{u} \\ \vdots \\ \mathbf{Q}^N\mathbf{u} \end{matrix}\right\} = \left\{\begin{matrix} \begin{pmatrix} \sum_k u_k Q^1_{1k} \\ \vdots \\ \sum_k u_k Q^1_{N1} \end{pmatrix} \\ \vdots \\ \begin{pmatrix} \sum_k u_k Q^N_{1k} \\ \vdots \\ \sum_k u_k Q^N_{N1} \end{pmatrix} \end{matrix}\right\}. \tag{S3.9}$$

Eqs. (5.5) and (5.6) give matrix-vectors of the same shape with $\mathsf{Q}$.

$$\left\{ \mathbf{P}^1 \cdots \mathbf{P}^N \right\} \left\{\begin{matrix} \mathbf{Q}^1 \\ \vdots \\ \mathbf{Q}^N \end{matrix}\right\} = \sum_{i=1}^{N} \mathbf{P}^i\mathbf{Q}^i \tag{S3.10}$$

As long as calculable, multiplication defined by Eqs. (S3.3-3.6) satisfy the associative law, i.e.,

$$\begin{aligned}
\mathbf{u}^T(\mathsf{Q}\mathbf{u}) &= (\mathbf{u}^T\mathsf{Q})\mathbf{u}, \\
\mathbf{u}^T(\mathsf{Q}\mathbf{A}) &= (\mathbf{u}^T\mathsf{Q})\mathbf{A}, \\
\mathbf{A}^T(\mathsf{Q}\mathbf{u}) &= (\mathbf{A}^T\mathsf{Q})\mathbf{u}, \\
\mathbf{A}^T(\mathsf{Q}\mathbf{B}) &= (\mathbf{A}^T\mathsf{Q})\mathbf{B}, \\
\mathbf{u}^T(\mathbf{A}^T\mathsf{Q}) &= (\mathbf{u}^T\mathbf{A}^T)\mathsf{Q},
\end{aligned}$$

$$\begin{aligned}
\mathbf{A}^T(\mathbf{u}^T\mathsf{Q}) &= (\mathbf{A}^T\mathbf{u}^T)\mathsf{Q}, \\
\mathbf{A}^T(\mathbf{B}^T\mathsf{Q}) &= (\mathbf{A}^T\mathbf{B}^T)\mathsf{Q}, \\
\mathsf{Q}(\mathbf{Au}) &= (\mathsf{Q}\mathbf{A})\mathbf{u}, \\
\mathsf{Q}(\mathbf{uA}) &= (\mathsf{Q}\mathbf{u})\mathbf{A}, \\
\mathsf{Q}(\mathbf{AB}) &= (\mathsf{Q}\mathbf{A})\mathbf{B}.
\end{aligned} \tag{S3.11}$$

Note that, for example, the left sides of $(\mathsf{Q}\mathbf{u})\mathbf{u} = \mathsf{Q}(\mathbf{uu})$ and $\mathbf{u}^T(\mathbf{u}^T\mathsf{Q}) = (\mathbf{u}^T\mathbf{u}^T)\mathsf{Q}$ are calculable but their right sides are not, because the multiplications $\mathbf{uu}$ and $\mathbf{u}^T\mathbf{u}^T$ are not defined, i.e., not calculable, although such multiplications do not appear in the analysis in this paper.

As for the distributive law, the following relationships hold good,

$$\begin{aligned}
\mathsf{Q}[\mathbf{a}+\mathbf{b}] &= \mathsf{Q}\mathbf{a}+\mathsf{Q}\mathbf{b}, \\
[\mathbf{a}+\mathbf{b}]^T\mathsf{Q} &= \mathbf{a}^T\mathsf{Q}+\mathbf{b}^T\mathsf{Q}, \\
\mathsf{Q}[\mathbf{A}+\mathbf{B}] &= \mathsf{Q}\mathbf{A}+\mathsf{Q}\mathbf{B}, \\
[\mathbf{A}+\mathbf{B}]^T\mathsf{Q} &= \mathbf{A}^T\mathsf{Q}+\mathbf{B}^T\mathsf{Q},
\end{aligned} \tag{S3.12}$$

where $\mathbf{A}$ and $\mathbf{B}$ are $N \times N$ matrices.

#### S3.3. Another multiplication

We define another multiplication between $\mathbf{u}$ and $\mathsf{Q}$ as

$$\langle \mathbf{u}^T, \mathsf{Q} \rangle = u_1\mathbf{Q}^1 + ... + u_N\mathbf{Q}^N, \tag{S3.13}$$

which is extended for a matrix $\mathbf{B} = (\mathbf{b}_1, ..., \mathbf{b}_N) = \begin{pmatrix} \tilde{\mathbf{b}}_1^T \\ \vdots \\ \tilde{\mathbf{b}}_N^T \end{pmatrix}$ as

$$\begin{aligned}
\langle \mathbf{B}^T, \mathsf{Q} \rangle &= \left\langle \begin{pmatrix} \mathbf{b}_1^T \\ \vdots \\ \mathbf{b}_N^T \end{pmatrix}, \mathsf{Q} \right\rangle = \left\{ \begin{array}{c} \langle \mathbf{b}_1^T, \mathsf{Q} \rangle \\ \vdots \\ \langle \mathbf{b}_N^T, \mathsf{Q} \rangle \end{array} \right\}, \\
\langle \mathbf{B}, \mathsf{Q} \rangle &= \left\langle \begin{pmatrix} \tilde{\mathbf{b}}_1^T \\ \vdots \\ \tilde{\mathbf{b}}_N^T \end{pmatrix}, \mathsf{Q} \right\rangle = \left\{ \begin{array}{c} \langle \tilde{\mathbf{b}}_1^T, \mathsf{Q} \rangle \\ \vdots \\ \langle \tilde{\mathbf{b}}_N^T, \mathsf{Q} \rangle \end{array} \right\}.
\end{aligned} \tag{S3.14}$$

Note that $\langle , \rangle$ is not defined for the arguments in the reverse order; its first argument must be a usual vector or matrix.

As long as calculable, multiplication defined by Eqs. (S3.13) and (S3.14) satisfy the distributive law, i.e.,

$$\begin{aligned}
\langle [\mathbf{a}+\mathbf{b}]^T, \mathsf{Q} \rangle &= \langle \mathbf{a}^T, \mathsf{Q} \rangle + \langle \mathbf{b}^T, \mathsf{Q} \rangle \\
\langle \mathbf{a}^T, \mathsf{P}+\mathsf{Q} \rangle &= \langle \mathbf{a}^T, \mathsf{P} \rangle + \langle \mathbf{a}^T, \mathsf{Q} \rangle \\
\langle [\mathbf{a}+\mathbf{b}]^T, \mathsf{Q} \rangle &= \langle \mathbf{a}^T, \mathsf{Q} \rangle + \langle \mathbf{b}^T, \mathsf{Q} \rangle \\
\langle \mathbf{a}^T, \mathsf{P}+\mathsf{Q} \rangle &= \langle \mathbf{a}^T, \mathsf{P} \rangle + \langle \mathbf{a}^T, \mathsf{Q} \rangle
\end{aligned}$$

where $\mathbf{a}$ and $\mathbf{b}$ are column vectors of length $N$, and $\mathsf{P}$ is a matrix-vector of the same shape with $\mathsf{Q}$.

#### S3.4. Transposition

We define the transpose $\mathsf{Q}^T$ as a row vector with its component matrices transposed,

$$\mathsf{Q}^T = \{ \ \mathbf{Q}^{1T} \ \cdots \ \mathbf{Q}^{NT} \ \}. \tag{S3.15}$$

We also define three kinds of additional transposes of $\mathsf{Q}$ that keep its shape. For its each scalar component $Q_{i,jk}$, the first transpose $\mathsf{Q}^t$ swaps $j$ and $k$, the second transpose $\mathsf{Q}^c$ swaps $i$ and $k$, and the third transpose $\mathsf{Q}^r$ swaps $i$ and $j$. More intuitively, in the two-dimensional case for example, $\mathsf{Q}^t$ transposes each of its component matrices

$$\mathsf{Q}^t = \begin{pmatrix} \mathbf{Q}^{1T} \\ \mathbf{Q}^{2T} \end{pmatrix}, \tag{S3.16}$$

$\mathsf{Q}^c$ swaps the columns highlighted with gray color between the component matrices,

$$\begin{aligned} \mathsf{Q}^c &= \left\{ \begin{matrix} \begin{pmatrix} Q^1_{11} & Q^1_{12} \\ Q^1_{21} & Q^1_{22} \end{pmatrix} \\ \begin{pmatrix} Q^2_{11} & Q^2_{12} \\ Q^2_{21} & Q^2_{22} \end{pmatrix} \end{matrix} \right\}^c \\ &= \left\{ \begin{matrix} \begin{pmatrix} Q^1_{11} & Q^2_{11} \\ Q^1_{21} & Q^2_{21} \end{pmatrix} \\ \begin{pmatrix} Q^1_{12} & Q^2_{12} \\ Q^1_{22} & Q^2_{22} \end{pmatrix} \end{matrix} \right\}, \end{aligned} \tag{S3.17}$$

and $\mathsf{Q}^r$ swaps the rows highlighted with gray color between the component matrices

$$\begin{aligned} \mathsf{Q}^r &= \left\{ \begin{matrix} \begin{pmatrix} Q^1_{11} & Q^1_{12} \\ Q^1_{21} & Q^1_{22} \end{pmatrix} \\ \begin{pmatrix} Q^2_{11} & Q^2_{12} \\ Q^2_{21} & Q^2_{22} \end{pmatrix} \end{matrix} \right\}^r \\ &= \left\{ \begin{matrix} \begin{pmatrix} Q^1_{11} & Q^1_{12} \\ Q^2_{11} & Q^2_{12} \end{pmatrix} \\ \begin{pmatrix} Q^1_{21} & Q^1_{22} \\ Q^2_{21} & Q^2_{22} \end{pmatrix} \end{matrix} \right\}. \end{aligned} \tag{S3.18}$$

Clearly, $[\mathsf{Q}^t]^t = \mathsf{Q}$, $[\mathsf{Q}^c]^c = \mathsf{Q}$, and $[\mathsf{Q}^r]^r = \mathsf{Q}$ hold good.

#### S3.5. Transformation rules

On the basis of the above definitions, the following transformation rules are derived in Appendix E,

$$\mathbf{b}^T\mathbf{u}^T\mathsf{Q} = \mathbf{u}^T\langle\mathbf{b}^T, \mathsf{Q}\rangle, \tag{S3.19}$$

$$\mathbf{B}\mathbf{u}^T\mathsf{Q} = \mathbf{u}^T\langle\mathbf{B}, \mathsf{Q}\rangle, \tag{S3.20}$$

$$\mathbf{u}^T\mathsf{Q}^c = [\mathbf{u}^T\mathsf{Q}]^T, \tag{S3.21}$$

$$\mathbf{u}^T\mathsf{Q}^r = \langle\mathbf{u}^T, \mathsf{Q}\rangle, \tag{S3.22}$$

$$\mathbf{B}\mathsf{Q}^c = (\mathbf{B}\mathsf{Q})^c, \tag{S3.23}$$

$$\mathsf{Q}^r\mathbf{B} = (\mathsf{Q}\mathbf{B})^r, \tag{S3.24}$$

$$\mathbf{B}\mathsf{Q}^r = \langle\mathbf{B}, \mathsf{Q}\rangle^r, \tag{S3.25}$$

$$(\mathsf{Q}\mathbf{B})^c = \langle\mathbf{B}^T, \mathsf{Q}^c\rangle, \tag{S3.26}$$

$$\mathbf{A}\langle\mathbf{u}^T, \mathsf{Q}\rangle = \langle\mathbf{u}^T, \mathbf{A}\mathsf{Q}\rangle \quad , \quad \langle\mathbf{u}^T, \mathsf{Q}\rangle\mathbf{A} = \langle\mathbf{u}^T, \mathsf{Q}\mathbf{A}\rangle \tag{S3.27}$$

$$\mathbf{A}\langle\mathbf{B}, \mathsf{Q}\rangle = \langle\mathbf{B}, \mathbf{A}\mathsf{Q}\rangle \quad , \quad \langle\mathbf{B}, \mathsf{Q}\rangle\mathbf{A} = \langle\mathbf{B}, \mathsf{Q}\mathbf{A}\rangle \tag{S3.28}$$

$$\langle\mathbf{B}, \langle\mathbf{A}, \mathsf{Q}\rangle\rangle = \langle\mathbf{B}\mathbf{A}, \mathsf{Q}\rangle \tag{S3.29}$$

$$\mathsf{Q}^{rcr} = \mathsf{Q}^{\mathrm{crc}} = \mathsf{Q}^t \tag{S3.30}$$

When the component matrices of Q are symmetric, i.e., $\mathsf{Q}^t$=Q, then Q trivially satisfies

$$\begin{aligned}\mathsf{Q}^{cr} &= \mathsf{Q}^{c},\\ \mathsf{Q}^{rc} &= \mathsf{Q}^{r},\\ \mathsf{Q}^{crc} &= \mathsf{Q}^{rcr}{=}\mathsf{Q}.\end{aligned} \tag{S3.31}$$

### S4. Appendix D: Derivation of Christoffel symbols

Here we derive Eq. (S1.9), as a usage example of the matrix-vector defined in Appendix C. We denote the geodesic coordinates $\tilde{\mathbf{u}}$ for a point $\hat{\mathbf{u}}$ by

$$\mathbf{u} = \tilde{\mathbf{u}} - \frac{1}{2}[\tilde{\mathbf{u}}-\hat{\mathbf{u}}]^T\Gamma[\tilde{\mathbf{u}}-\hat{\mathbf{u}}] \tag{S4.1}$$

where "^" is omitted from $\hat{\Gamma}$ for convenience, and $\hat{\mathbf{u}}=\mathbf{0}$ is assumed without loss of generality. From Eq. (S1.1),

$$\begin{aligned}\delta\mathbf{u} &= [\mathbf{u}+\delta\mathbf{u}]-\mathbf{u}\\ &= \left\{[\tilde{\mathbf{u}}+\delta\tilde{\mathbf{u}}]-\frac{1}{2}[\tilde{\mathbf{u}}+\delta\tilde{\mathbf{u}}]^T\Gamma[\tilde{\mathbf{u}}+\delta\tilde{\mathbf{u}}]\right\}-\left\{\tilde{\mathbf{u}}-\frac{1}{2}\tilde{\mathbf{u}}^T\Gamma\tilde{\mathbf{u}}\right\}\\ &= \delta\tilde{\mathbf{u}}-\frac{1}{2}\tilde{\mathbf{u}}^T\Gamma\tilde{\mathbf{u}}-\frac{1}{2}\tilde{\mathbf{u}}^T\Gamma\delta\tilde{\mathbf{u}}-\frac{1}{2}\delta\tilde{\mathbf{u}}^T\Gamma\tilde{\mathbf{u}}-\frac{1}{2}\delta\tilde{\mathbf{u}}^T\Gamma\delta\tilde{\mathbf{u}}+\frac{1}{2}\tilde{\mathbf{u}}^T\Gamma\tilde{\mathbf{u}}\\ &= \delta\tilde{\mathbf{u}}-\frac{1}{2}\tilde{\mathbf{u}}^T\Gamma\delta\tilde{\mathbf{u}}-\frac{1}{2}\delta\tilde{\mathbf{u}}^T\Gamma\tilde{\mathbf{u}}-\frac{1}{2}\delta\tilde{\mathbf{u}}^T\Gamma\delta\tilde{\mathbf{u}}\\ &= \delta\tilde{\mathbf{u}}-\tilde{\mathbf{u}}^T\Gamma\delta\tilde{\mathbf{u}}-\frac{1}{2}\delta\tilde{\mathbf{u}}^T\Gamma\delta\tilde{\mathbf{u}}\\ &= [\mathbf{I}-\tilde{\mathbf{u}}^T\Gamma]\delta\tilde{\mathbf{u}}-\frac{1}{2}\delta\tilde{\mathbf{u}}^T\Gamma\delta\tilde{\mathbf{u}},\\ d\mathbf{u} &= [\mathbf{I}-\tilde{\mathbf{u}}^T\Gamma]d\tilde{\mathbf{u}},\\ dl^2 &= d\mathbf{u}^T\mathbf{M}(\mathbf{u})d\mathbf{u}\\ &= d\tilde{\mathbf{u}}^T[\mathbf{I}-\tilde{\mathbf{u}}^T\Gamma]^T\mathbf{M}(\mathbf{u})[\mathbf{I}-\tilde{\mathbf{u}}^T\Gamma]d\tilde{\mathbf{u}}=d\tilde{\mathbf{u}}^T\tilde{\mathbf{M}}(\tilde{\mathbf{u}})d\tilde{\mathbf{u}},\end{aligned}$$

where $\delta\tilde{\mathbf{u}}^T\Gamma\tilde{\mathbf{u}}$ is transformed into $\tilde{\mathbf{u}}^T\Gamma\delta\tilde{\mathbf{u}}$ by using $[\hat{\mathbf{\Gamma}}^i]^T=\hat{\mathbf{\Gamma}}^i$ for all $i=1,...,N$, we can express the mutation metric as

$$\tilde{\mathbf{M}}(\tilde{\mathbf{u}}) = [I-\tilde{\mathbf{u}}^T\Gamma]^T\mathbf{M}(\mathbf{u})[I-\tilde{\mathbf{u}}^T\Gamma], \tag{S4.2}$$

By using Eqs. (S3.19-S3.31),we transform Eq. (S4.2) into

$$\begin{aligned}\tilde{\mathbf{M}}(\tilde{\mathbf{u}}) &= [I-\langle\tilde{\mathbf{u}}^T,\Gamma^r\rangle]^T\mathbf{M}(\mathbf{u})[I-\langle\tilde{\mathbf{u}}^T,\Gamma^r\rangle]\\ &= [I-\langle\tilde{\mathbf{u}}^T,\Phi\rangle]^T\mathbf{M}(\mathbf{u})[I-\langle\tilde{\mathbf{u}}^T,\Phi\rangle],\\ \Phi &= \left\{\begin{array}{c}\mathbf{\Phi}^1\\ \vdots\\ \mathbf{\Phi}^N\end{array}\right\}=\Gamma^r.\end{aligned} \tag{S4.3}$$

Below, we derive $\Gamma$ by requiring

$$\frac{\partial\tilde{\mathbf{M}}(\hat{\mathbf{u}})}{\partial\tilde{u}_i} = \left[\frac{\partial\tilde{\mathbf{M}}(\tilde{\mathbf{u}})}{\partial\tilde{u}_i}\right]_{\tilde{\mathbf{u}}=\hat{\mathbf{u}}}=\mathbf{0} \tag{S4.4}$$

for all $i=1,...,N$.

First, we see

$$\begin{aligned}
\frac{\partial\tilde{\mathbf{M}}(\tilde{\mathbf{u}})}{\partial\tilde{u}_i} &= \frac{\partial[I-\langle\tilde{\mathbf{u}}^T,\Phi\rangle]^T}{\partial\tilde{u}_i}\mathbf{M}(\mathbf{u})[I-\langle\tilde{\mathbf{u}}^T,\Phi\rangle] \\
&\quad+[I-\langle\tilde{\mathbf{u}}^T,\Phi\rangle]^T\frac{\partial\mathbf{M}(\mathbf{u})}{\partial\tilde{u}_i}[I-\langle\tilde{\mathbf{u}}^T,\Phi\rangle] \\
&\quad+[I-\langle\tilde{\mathbf{u}}^T,\Phi\rangle]^T\mathbf{M}(\mathbf{u})\frac{\partial[I-\langle\tilde{\mathbf{u}}^T,\Phi\rangle]}{\partial\tilde{u}_i} \\
&= -\mathbf{\Phi}^{iT}\mathbf{M}(\mathbf{u})[I-\langle\tilde{\mathbf{u}}^T,\Phi\rangle] \\
&\quad+[I-\langle\tilde{\mathbf{u}}^T,\Phi\rangle]^T\frac{\partial\mathbf{M}(\mathbf{u})}{\partial\tilde{u}_i}[I-\langle\tilde{\mathbf{u}}^T,\Phi\rangle] \\
&\quad-[I-\langle\tilde{\mathbf{u}}^T,\Phi\rangle]^T\mathbf{M}(\mathbf{u})\mathbf{\Phi}^i. \qquad \text{(S4.5)}
\end{aligned}$$

Substitution of Eq. (S4.5) into Eq. (S4.4) gives

$$\frac{\partial\tilde{\mathbf{M}}(\hat{\mathbf{u}})}{\partial\tilde{u}_i} = \mathbf{\Phi}^{iT}\hat{\mathbf{M}}+\frac{\partial\mathbf{M}(\hat{\mathbf{u}})}{\partial\tilde{u}_i}-\hat{\mathbf{M}}\mathbf{\Phi}^i=\mathbf{0} \qquad \text{(S4.6)}$$

with $\hat{\mathbf{M}}=\mathbf{M}(\hat{\mathbf{u}})$ for all $i=1,...,N$, which are combined into

$$\begin{aligned}
\nabla\tilde{\mathbf{M}}(\hat{\mathbf{u}})=\begin{Bmatrix}\frac{\partial\tilde{\mathbf{M}}(\hat{\mathbf{u}})}{\partial\tilde{u}_1}\\ \vdots\\ \frac{\partial\tilde{\mathbf{M}}(\hat{\mathbf{u}})}{\partial\tilde{u}_N}\end{Bmatrix} &= \begin{Bmatrix}\frac{\partial\mathbf{M}(\hat{\mathbf{u}})}{\partial\tilde{u}_1}\\ \vdots\\ \frac{\partial\mathbf{M}(\hat{\mathbf{u}})}{\partial\tilde{u}_N}\end{Bmatrix}-\begin{Bmatrix}\mathbf{\Phi}^{1T}\hat{\mathbf{M}}\\ \vdots\\ \mathbf{\Phi}^{NT}\hat{\mathbf{M}}\end{Bmatrix}-\begin{Bmatrix}\hat{\mathbf{M}}\mathbf{\Phi}^1\\ \vdots\\ \hat{\mathbf{M}}\mathbf{\Phi}^N\end{Bmatrix} \\
&= \begin{Bmatrix}\sum_{j=1}^N\frac{\partial\mathbf{M}(\hat{\mathbf{u}})}{\partial u_j}\left[\frac{\partial u_j}{\partial\tilde{u}_1}\right]_{\mathbf{u}=\hat{\mathbf{u}}}\\ \vdots\\ \sum_{j=1}^N\frac{\partial\mathbf{M}(\hat{\mathbf{u}})}{\partial u_j}\left[\frac{\partial u_j}{\partial\tilde{u}_N}\right]_{\mathbf{u}=\hat{\mathbf{u}}}\end{Bmatrix}-\Phi^t\hat{\mathbf{M}}-\hat{\mathbf{M}}\Phi \\
&= \begin{Bmatrix}\frac{\partial\mathbf{M}(\hat{\mathbf{u}})}{\partial u_1}\\ \vdots\\ \frac{\partial\mathbf{M}(\hat{\mathbf{u}})}{\partial u_N}\end{Bmatrix}-\Phi^t\hat{\mathbf{M}}-\hat{\mathbf{M}}\Phi \\
&= \nabla\mathbf{M}(\hat{\mathbf{u}})-\Phi^t\hat{\mathbf{M}}-\hat{\mathbf{M}}\Phi \\
&= \mathbf{0}. \qquad \text{(S4.7)}
\end{aligned}$$

By using $\Phi$=$\Gamma^r$ and Eqs. (S3.19-S3.31), we transform Eq. (S4.7) into

$$\begin{aligned}
\nabla\mathbf{M}(\hat{\mathbf{u}}) &= \Phi^t\hat{\mathbf{M}}+\hat{\mathbf{M}}\Phi=[\Gamma^r]^{rcr}\hat{\mathbf{M}}+\hat{\mathbf{M}}\Gamma^r \\
&= \Gamma^{cr}\hat{\mathbf{M}}+\hat{\mathbf{M}}\Gamma^r \\
&= \Gamma^c\hat{\mathbf{M}}+\hat{\mathbf{M}}\Gamma^r, \qquad \text{(S4.8)}
\end{aligned}$$

and find

$$\begin{aligned}
[[\nabla\mathbf{M}(\hat{\mathbf{u}})]^r+[\nabla\mathbf{M}(\hat{\mathbf{u}})]^c-\nabla\mathbf{M}(\hat{\mathbf{u}})] &= [\Gamma^c\hat{\mathbf{M}}]^r+[\hat{\mathbf{M}}\Gamma^r]^r+[\Gamma^c\hat{\mathbf{M}}]^c+[\hat{\mathbf{M}}\Gamma^r]^c \\
&\quad-\Gamma^c\hat{\mathbf{M}}-\hat{\mathbf{M}}\Gamma^r \\
&= \Gamma^c\hat{\mathbf{M}}+\langle\hat{\mathbf{M}},\Gamma\rangle+\langle\hat{\mathbf{M}},\Gamma\rangle+\hat{\mathbf{M}}\Gamma^r \\
&\quad-\Gamma^c\hat{\mathbf{M}}-\hat{\mathbf{M}}\Gamma^r \\
&= 2\langle\hat{\mathbf{M}},\Gamma\rangle, \qquad \text{(S4.9)}
\end{aligned}$$

by using Eqs. (S3.19-S3.31), i.e.,

$$\begin{aligned}
[\Gamma^c\hat{\mathbf{M}}]^r &= \Gamma^{cr}\hat{\mathbf{M}} = \Gamma^c\hat{\mathbf{M}}, \\
[\hat{\mathbf{M}}\Gamma^r]^r &= \langle\hat{\mathbf{M}}, \Gamma\rangle, \\
[\Gamma^c\hat{\mathbf{M}}]^c &= \langle\hat{\mathbf{M}}, \Gamma^{cc}\rangle = \langle\hat{\mathbf{M}}, \Gamma\rangle, \\
[\hat{\mathbf{M}}\Gamma^r]^c &= \hat{\mathbf{M}}\Gamma^{rc} = \hat{\mathbf{M}}\Gamma^r.
\end{aligned} \tag{S4.10}$$

Since Eq. (S3.28) allows $\langle\hat{\mathbf{M}}^{-1}, \langle\hat{\mathbf{M}}, \Gamma\rangle\rangle = \Gamma$, we get

$$\begin{aligned}
\Gamma &= \frac{1}{2}\langle\hat{\mathbf{M}}^{-1}, [\nabla\mathbf{M}(\hat{\mathbf{u}})]^r + [\nabla\mathbf{M}(\hat{\mathbf{u}})]^c - \nabla\mathbf{M}(\hat{\mathbf{u}})\rangle \\
&= \frac{1}{2}\langle\mathbf{M}(\hat{\mathbf{u}})^{-1}, [\nabla\mathbf{M}(\hat{\mathbf{u}})]^r + [\nabla\mathbf{M}(\hat{\mathbf{u}})]^c - \nabla\mathbf{M}(\hat{\mathbf{u}})\rangle
\end{aligned} \tag{S4.11}$$

which is identical to Eqs. (S1.7) and (S1.9).

### S5. Appendix E: Derivation of Eq. (S2.32)

From

$$\mathbf{B}[\mathbf{B}^T\mathbf{B} + \mathbf{I}_M] = \mathbf{B}\mathbf{B}^T\mathbf{B} + \mathbf{B} = [\mathbf{I}_N + \mathbf{B}\mathbf{B}^T]\mathbf{B}, \tag{S5.1}$$

we get

$$\mathbf{B}[\mathbf{B}^T\mathbf{B} + \mathbf{I}_M]^{-1} = [\mathbf{I}_N + \mathbf{B}\mathbf{B}^T]^{-1}\mathbf{B}. \tag{S5.2}$$

### S6. Appendix F: Derivation of transformation rule for Christoffel symbols

#### S6.1. Transformation rule

We consider a coordinate system $\mathbf{z} = (z_1, ..., z_N)^T$, such that

$$\begin{aligned}
\mathbf{u} - \mathbf{u}_0 &= \mathbf{W}[\mathbf{z} - \mathbf{z}_0] \\
&\quad + \frac{1}{2}[\mathbf{z} - \mathbf{z}_0]^T \mathsf{H}\, [\mathbf{z} - \mathbf{z}_0]
\end{aligned} \tag{S6.1}$$

where

$$\mathbf{W} = \begin{pmatrix} \frac{\partial u_1}{\partial z_1} & \cdots & \frac{\partial u_1}{\partial z_N} \\ \vdots & \ddots & \vdots \\ \frac{\partial u_N}{\partial z_1} & \cdots & \frac{\partial u_N}{\partial z_N} \end{pmatrix}_{\mathbf{z}=\mathbf{z}_0},$$

$$\mathsf{H} = \begin{Bmatrix} \mathbf{H}^1 \\ \vdots \\ \mathbf{H}^N \end{Bmatrix} = \begin{Bmatrix} \begin{pmatrix} \frac{\partial^2 u_1}{\partial z_1 \partial z_1} & \cdots & \frac{\partial^2 u_1}{\partial z_1 \partial z_N} \\ \vdots & \ddots & \vdots \\ \frac{\partial^2 u_1}{\partial z_N \partial z_1} & \cdots & \frac{\partial^2 u_1}{\partial z_N \partial z_N} \end{pmatrix}_{\mathbf{z}=\mathbf{z}_0} \\ \vdots \\ \begin{pmatrix} \frac{\partial^2 u_N}{\partial z_1 \partial z_1} & \cdots & \frac{\partial^2 u_N}{\partial z_1 \partial z_N} \\ \vdots & \ddots & \vdots \\ \frac{\partial^2 u_N}{\partial z_N \partial z_1} & \cdots & \frac{\partial^2 u_N}{\partial z_N \partial z_N} \end{pmatrix}_{\mathbf{z}=\mathbf{z}_0} \end{Bmatrix}. \tag{S6.2}$$

where $\mathbf{H}^1, ..., \mathbf{H}^N$ are all symmetric, which gives $\mathsf{H}^t$=$\mathsf{H}$. In differential geometry, Christoffel symbols (of the second kind) at $\mathbf{z}_0$ in coordinates $\mathbf{z}$ are given by

$$[\Gamma_z]^i_{jk}(\mathbf{z}_0) = \sum_p \frac{\partial z_i}{\partial u_p} \sum_m \frac{\partial u_m}{\partial z_k} \sum_q \frac{\partial u_q}{\partial z_j} \Gamma^p_{qm}(\mathbf{u}_0) + \sum_r \frac{\partial z_i}{\partial u_r} \frac{\partial^2 u_r}{\partial z_j \partial z_k} \tag{S6.3}$$

(in pp64 in Hobson et al. 2007), where all derivatives are evaluated at $\mathbf{z} = \mathbf{z}_0$ (i.e., $\mathbf{u} = \mathbf{u}_0$). Eq. (S6.3) can be expressed in a vector-matrix form as

$$\Gamma_z(\mathbf{z}_0) = \langle \mathbf{W}^{-1}, \mathbf{W}^T \Gamma(\mathbf{u}_0) \mathbf{W} + \mathsf{H} \rangle. \tag{S6.4}$$

In the following subsections, we derive Eq. (S6.4).

#### S6.2. Derivation of transformation rule

##### S6.2.1 Decomposition of coordinate transformation

For convenience, we decompose Eq. (S6.1) into a linear transformation into coordinates $\mathbf{y} = (y_1, ..., y_N)^T$,

$$\mathbf{u} - \mathbf{u}_0 = \mathbf{W}[\mathbf{y} - \mathbf{y}_0] \tag{S6.5}$$

and a subsequent nonlinear transformation into coordinates $\mathbf{z}$,

$$\begin{aligned} \mathbf{y} - \mathbf{y}_0 &= \mathbf{z} - \mathbf{z}_0 + \frac{1}{2}[\mathbf{z} - \mathbf{z}_0]^T \mathsf{Q} [\mathbf{z} - \mathbf{z}_0], \\ \mathsf{Q} &= \langle \mathbf{W}^{-1}, \mathsf{H} \rangle, \end{aligned} \tag{S6.6}$$

where $\mathsf{Q}^t$=$\mathsf{Q}$ is satisfied because of $\mathsf{H}^t$=$\mathsf{H}$. Note that substituting Eqs. (S6.6) into Eq. (S6.5) recovers Eq. (S6.1):

$$\begin{aligned} \mathbf{u} - \mathbf{u}_0 &= \mathbf{W}[\mathbf{z} - \mathbf{z}_0] + \mathbf{W}\frac{1}{2}[\mathbf{z} - \mathbf{z}_0]^T \langle \mathbf{W}^{-1}, \mathsf{H} \rangle [\mathbf{z} - \mathbf{z}_0] \\ &= \mathbf{W}[\mathbf{z} - \mathbf{z}_0] + \frac{1}{2}[\mathbf{z} - \mathbf{z}_0]^T \langle \mathbf{W}, \langle \mathbf{W}^{-1}, \mathsf{H} \rangle \rangle [\mathbf{z} - \mathbf{z}_0] \\ &= \mathbf{W}[\mathbf{z} - \mathbf{z}_0] + \frac{1}{2}[\mathbf{z} - \mathbf{z}_0]^T \mathsf{H} [\mathbf{z} - \mathbf{z}_0]. \end{aligned} \tag{S6.7}$$

##### S6.2.2 Linear transformation

We denote by $\breve{\mathbf{u}}$ the geodesic coordinates for $\mathbf{u}_0$ in coordinates $\mathbf{u}$, expressed as

$$\mathbf{u} - \mathbf{u}_0 = \breve{\mathbf{u}} - \mathbf{u}_0 + \frac{1}{2}[\breve{\mathbf{u}} - \mathbf{u}_0]^T \Gamma(\mathbf{u}_0) [\breve{\mathbf{u}} - \mathbf{u}_0], \tag{S6.8}$$

where the first derivative of the mutation metric in $\breve{\mathbf{u}}$ is zero at $\breve{\mathbf{u}} = \mathbf{u}_0$, i.e., $\partial \breve{\mathbf{M}}(\breve{\mathbf{u}}) / \partial \breve{x}_i = \mathbf{0}$ at $\breve{\mathbf{u}} = \mathbf{u}_0$ holds good for all $i = 1, ..., N$.

We consider a linear transformation of $\breve{\mathbf{u}}$ into $\breve{\mathbf{y}}$, defined by

$$\breve{\mathbf{u}} - \mathbf{u}_0 = \mathbf{W}[\breve{\mathbf{y}} - \mathbf{y}_0]. \tag{S6.9}$$

Substituting Eqs. (S6.5) and (S6.9) into Eq. (S6.8) gives

$$\mathbf{W}[\mathbf{y} - \mathbf{y}_0] = \mathbf{W}[\breve{\mathbf{y}} - \mathbf{y}_0] + \frac{1}{2}[\breve{\mathbf{y}} - \mathbf{y}_0] \mathbf{W}^T \Gamma(\mathbf{u}_0) \mathbf{W} [\breve{\mathbf{y}} - \mathbf{y}_0]. \tag{S6.10}$$

Multiplying $\mathbf{W}^{-1}$ by left-side gives

$$\begin{aligned}\mathbf{y}-\mathbf{y}_0 &= \check{\mathbf{y}}-\mathbf{y}_0+\frac{1}{2}\mathbf{W}^{-1}[\check{\mathbf{y}}-\mathbf{y}_0]\mathbf{W}^T\Gamma(\mathbf{u}_0)\mathbf{W}[\check{\mathbf{y}}-\mathbf{y}_0] \\ &= \check{\mathbf{y}}-\mathbf{y}_0+\frac{1}{2}[\check{\mathbf{y}}-\mathbf{y}_0]\langle\mathbf{W}^{-1},\mathbf{W}^T\Gamma(\mathbf{u}_0)\mathbf{W}\rangle[\check{\mathbf{y}}-\mathbf{y}_0] \\ &= \check{\mathbf{y}}-\mathbf{y}_0+\frac{1}{2}[\check{\mathbf{y}}-\mathbf{y}_0]\Gamma_y(\mathbf{y}_0)[\check{\mathbf{y}}-\mathbf{y}_0], \qquad \text{(S6.11)}\end{aligned}$$

with

$$\Gamma_y(\mathbf{y}_0) = \langle\mathbf{W}^{-1},\mathbf{W}^T\Gamma(\mathbf{u}_0)\mathbf{W}\rangle. \qquad \text{(S6.12)}$$

Since $\check{\mathbf{y}}$ is a linear transformation of $\check{\mathbf{u}}$, we see

$$\begin{aligned}\left[\frac{\partial\check{\mathbf{M}}_y(\check{\mathbf{y}})}{\partial\check{y}_i}\right]_{\check{\mathbf{y}}=\mathbf{y}_0} &= \left[\mathbf{W}^T\frac{\partial\check{\mathbf{M}}(\check{\mathbf{u}})}{\partial\check{y}_i}\mathbf{W}\right]_{\check{\mathbf{y}}=\mathbf{y}_0} \\ &= \left[\mathbf{W}^T\sum_j^N\frac{\partial\check{\mathbf{M}}(\check{\mathbf{u}})}{\partial\check{x}_j}\frac{\partial x_j}{\partial y_i}\mathbf{W}\right]_{\check{\mathbf{y}}=\mathbf{y}_0} \\ &= \mathbf{W}^T\sum_j^N\left[\frac{\partial\check{\mathbf{M}}(\check{\mathbf{u}})}{\partial\check{x}_j}\right]_{\check{\mathbf{u}}=\mathbf{u}_0}W_{ji}\mathbf{W} \\ &= \mathbf{0} \qquad \text{(S6.13)}\end{aligned}$$

for all $i=1$, ..., $N$. Thus, coordinates $\check{\mathbf{y}}$ are the geodesic coordinates for $\mathbf{y}_0$ in coordinates $\mathbf{y}$. Therefore, Christoffel symbols at $\mathbf{y}_0$ in coordinates $\mathbf{y}$ are given by Eq. (S6.12).

##### S6.2.3 Nonlinear transformation

From

$$\begin{aligned}dl^2 &= d\mathbf{u}^T\mathbf{M}(\mathbf{u})d\mathbf{u} \\ &= d\mathbf{y}^T\mathbf{W}^T\mathbf{M}(\mathbf{u})\mathbf{W}d\mathbf{y} \\ &= d\mathbf{y}^T\mathbf{M}_y(\mathbf{y})d\mathbf{y} \qquad \text{(S6.14)}\end{aligned}$$

we get the mutation metric $\mathbf{M}_y(\mathbf{y})$ in coordinates $\mathbf{y}$

$$\mathbf{M}_y(\mathbf{y}) = \mathbf{W}^T\mathbf{M}(\mathbf{u})\mathbf{W}. \qquad \text{(S6.15)}$$

In addition, we see from Eqs. (S6.6) that

$$\mathbf{dy} = [\mathbf{I}+[\mathbf{z}-\mathbf{z}_0]^T\mathsf{Q}]\mathbf{dz},$$

which gives

$$\begin{aligned}dl^2 &= \mathbf{dz}^T[\mathbf{I}+[\mathbf{z}-\mathbf{z}_0]^T\mathsf{Q}]^T\mathbf{M}_y(\mathbf{y})[\mathbf{I}+[\mathbf{z}-\mathbf{z}_0]^T\mathsf{Q}]\mathbf{dz} \qquad \text{(S6.16)} \\ &= \mathbf{dz}^T\mathbf{M}_z(\mathbf{z})\mathbf{dz}\end{aligned}$$

and thus

$$\mathbf{M}_z(\mathbf{z}) = [\mathbf{I}+[\mathbf{z}-\mathbf{z}_0]^T\mathsf{Q}]^T\mathbf{M}_y(\mathbf{y})[\mathbf{I}+[\mathbf{z}-\mathbf{z}_0]^T\mathsf{Q}], \qquad \text{(S6.17)}$$

By using Eq. (S3.22) (i.e., $[\mathbf{z}-\mathbf{z}_0]^T\mathsf{Q}$=$\langle[\mathbf{z}-\mathbf{z}_0]^T,\mathsf{Q}^r\rangle$), we transform Eq. (S6.17) into

$$\begin{aligned}\mathbf{M}_z(\mathbf{z}) &= [I+\langle[\mathbf{z}-\mathbf{z}_0]^T,\Phi\rangle]^T\mathbf{M}_y(\mathbf{y})[I+\langle[\mathbf{z}-\mathbf{z}_0]^T,\Phi\rangle] \\ \Phi &= \left\{\begin{array}{c}\mathbf{\Phi}^1 \\ \vdots \\ \mathbf{\Phi}^N\end{array}\right\}=\mathsf{Q}^r. \qquad \text{(S6.18)}\end{aligned}$$

Then we calculate the first derivative of $\mathbf{M}_z(\mathbf{z})$ at $\mathbf{z}=\mathbf{z}_0$ as

$$\begin{aligned}
\frac{\partial \mathbf{M}_z(\mathbf{z}_0)}{\partial z_i} &= \frac{\partial [I+\langle[\mathbf{z}-\mathbf{z}_0]^T,\Phi\rangle]^T}{\partial z_i}\mathbf{M}_y(\mathbf{y}_0) \\
&\quad +\frac{\partial \mathbf{M}_y(\mathbf{y}_0)}{\partial z_i} \\
&\quad +\mathbf{M}_y(\mathbf{y}_0)\frac{\partial [I+\langle[\mathbf{z}-\mathbf{z}_0]^T,\Phi\rangle]^T}{\partial z_i} \\
&= \mathbf{\Phi}^{iT}\mathbf{M}_y(\mathbf{y}_0)+\left[\sum_j \frac{\partial \mathbf{M}_y(\mathbf{y})}{\partial y_j}\frac{\partial y_j}{\partial z_i}\right]_{\mathbf{y}=\mathbf{y}_0}+\mathbf{M}_y(\mathbf{y}_0)\mathbf{\Phi}^i \\
&= \mathbf{\Phi}^{iT}\mathbf{M}_y(\mathbf{y}_0)+\frac{\partial \mathbf{M}_y(\mathbf{y}_0)}{\partial y_i}+\mathbf{M}_y(\mathbf{y}_0)\mathbf{\Phi}^i
\end{aligned} \tag{S6.19}$$

for all $i=1,...,N$, which are combined by using Eq. (S3.16) (i.e., $\left\{\begin{array}{c}\mathbf{\Phi}^{1T}\\ \vdots \\ \mathbf{\Phi}^{NT}\end{array}\right\}=\Phi^t$) into

$$\begin{aligned}
\nabla \mathbf{M}_z(\mathbf{z}_0)=\left\{\begin{array}{c}\frac{\partial \mathbf{M}_z(\mathbf{z}_0)}{\partial z_1}\\ \vdots \\ \frac{\partial \mathbf{M}_z(\mathbf{z}_0)}{\partial z_N}\end{array}\right\} &= \left\{\begin{array}{c}\frac{\partial \mathbf{M}_y(\mathbf{y}_0)}{\partial y_1}\\ \vdots \\ \frac{\partial \mathbf{M}_y(\mathbf{y}_0)}{\partial y_N}\end{array}\right\}+\left\{\begin{array}{c}\mathbf{\Phi}^{1T}\mathbf{M}_y(\mathbf{y}_0)\\ \vdots \\ \mathbf{\Phi}^{NT}\mathbf{M}_y(\mathbf{y}_0)\end{array}\right\}+\left\{\begin{array}{c}\mathbf{M}_y(\mathbf{y}_0)\mathbf{\Phi}^{1}\\ \vdots \\ \mathbf{M}_y(\mathbf{y}_0)\mathbf{\Phi}^{N}\end{array}\right\} \\
&= \nabla\mathbf{M}_y(\mathbf{y}_0)+\Phi^t\mathbf{M}_y(\mathbf{y}_0)+\mathbf{M}_y(\mathbf{y}_0)\Phi. \\
&= \nabla\mathbf{M}_{y0}+\Phi^t\mathbf{M}_{y0}+\mathbf{M}_{y0}\Phi.
\end{aligned} \tag{S6.20}$$

with $\mathbf{M}_{y0}=\mathbf{M}_y(\mathbf{y}_0)$ and $\nabla\mathbf{M}_{y0}=\nabla\mathbf{M}_y(\mathbf{y}_0)$.

By using $\Phi$=$\mathsf{Q}^r$ and Eqs. (S3.30) and (S3.31) (i.e., $\Phi^t$=$\Phi^{rcr}$ and $\mathsf{Q}^{cr}$=$\mathsf{Q}^c$), we transform Eq. (S6.20) into

$$\begin{aligned}
\nabla\mathbf{M}_z(\mathbf{z}_0) &= \nabla\mathbf{M}_{y0}+[\mathsf{Q}^r]^{rcr}\mathbf{M}_{y0}+\mathbf{M}_{y0}\mathsf{Q}^r \\
&= \nabla\mathbf{M}_{y0}+\mathsf{Q}^{cr}\mathbf{M}_{y0}+\mathbf{M}_{y0}\mathsf{Q}^r \\
&= \nabla\mathbf{M}_{y0}+\mathsf{Q}^{c}\mathbf{M}_{y0}+\mathbf{M}_{y0}\mathsf{Q}^r.
\end{aligned} \tag{S6.21}$$

Then, we find

$$\begin{aligned}
[\nabla\mathbf{M}_z(\mathbf{z}_0)]^r+[\nabla\mathbf{M}_z(\mathbf{z}_0)]^c-\nabla\mathbf{M}_z(\mathbf{z}_0) &= [\nabla\mathbf{M}_{y0}]^r+[\nabla\mathbf{M}_{y0}]^c-\nabla\mathbf{M}_{y0} \\
&\quad +2\langle\mathbf{M}_{y0},\mathsf{Q}\rangle.
\end{aligned} \tag{S6.22}$$

By using Eq. (S3.11) and Eq. (S3.29) we get Christoffel symbols for $\mathbf{z}_0$ in coordinates $\mathbf{z}$,

$$\begin{aligned}
\Gamma_z(\mathbf{z}_0) &= \frac{1}{2}\langle\mathbf{M}_{y0}^{-1},[\nabla\mathbf{M}_z(\mathbf{z}_0)]^r+[\nabla\mathbf{M}_z(\mathbf{z}_0)]^c-\nabla\mathbf{M}_z(\mathbf{z}_0)\rangle \\
&= \frac{1}{2}\langle\mathbf{M}_{y0}^{-1},[\nabla\mathbf{M}_{y0}]^r+[\nabla\mathbf{M}_{y0}]^c-\nabla\mathbf{M}_{y0}+2\langle\mathbf{M}_{y0},\mathsf{Q}\rangle\rangle \\
&= \frac{1}{2}\langle\mathbf{M}_{y0}^{-1},[\nabla\mathbf{M}_{y0}]^r+[\nabla\mathbf{M}_{y0}]^c-\nabla\mathbf{M}_{y0}\rangle+\langle\mathbf{M}_{y0}^{-1},\langle\mathbf{M}_{y0},\mathsf{Q}\rangle\rangle \\
&= \Gamma_y(\mathbf{y}_0)+\mathsf{Q}.
\end{aligned} \tag{S6.23}$$

Finally, substituting Eq. (S6.12) into Eq. (S6.23), we get

$$\begin{aligned}
\Gamma_z(\mathbf{z}_0) &= \Gamma_y(\mathbf{y}_0)+\mathsf{Q} \\
&= \langle\mathbf{W}^{-1},\mathbf{W}^T\Gamma(\mathbf{u}_0)\mathbf{W}\rangle+\langle\mathbf{W}^{-1},\mathsf{H}\rangle \\
&= \langle\mathbf{W}^{-1},\mathbf{W}^T\Gamma(\mathbf{u}_0)\mathbf{W}+\mathsf{H}\rangle.
\end{aligned} \tag{S6.24}$$

### S7. Appendix G: Derivations of Eqs. (S3.19-S3.30) in Appendix C

$$
\begin{aligned}
\mathbf{u} &= \begin{pmatrix} u_1 \\ u_2 \\ u_3 \end{pmatrix}, \\
\mathbf{z} &= \begin{pmatrix} z_1 \\ z_2 \\ z_3 \end{pmatrix}, \\
\mathbf{B} &= \begin{pmatrix} \mathbf{b}_1^T \\ \mathbf{b}_2^T \\ \mathbf{b}_3^T \end{pmatrix} = \begin{pmatrix} B_{11} & B_{12} & B_{13} \\ B_{21} & B_{22} & B_{23} \\ B_{31} & B_{32} & B_{33} \end{pmatrix}, \\
\mathbf{A} &= \begin{pmatrix} \mathbf{a}_1^T \\ \mathbf{a}_2^T \\ \mathbf{a}_3^T \end{pmatrix} = \begin{pmatrix} \boldsymbol{\alpha}_1 & \boldsymbol{\alpha}_2 & \boldsymbol{\alpha}_3 \end{pmatrix} = \begin{pmatrix} A_{11} & A_{12} & A_{13} \\ A_{21} & A_{22} & A_{23} \\ A_{31} & A_{32} & A_{33} \end{pmatrix}, \\
\mathbf{Q}_i &= \begin{pmatrix} Q_{11}^i & Q_{12}^i & Q_{13}^i \\ Q_{21}^i & Q_{22}^i & Q_{23}^i \\ Q_{31}^i & Q_{32}^i & Q_{33}^i \end{pmatrix} = \begin{pmatrix} \mathbf{q}_{i1} & \mathbf{q}_{i2} & \mathbf{q}_{i3} \end{pmatrix} = \begin{pmatrix} \mathbf{p}_{i1}^T \\ \mathbf{p}_{i2}^T \\ \mathbf{p}_{i3}^T \end{pmatrix}, \\
\mathsf{Q} &= \begin{Bmatrix} \mathbf{Q}_1 \\ \mathbf{Q}_2 \\ \mathbf{Q}_3 \end{Bmatrix} = \begin{Bmatrix} \begin{pmatrix} \mathbf{q}_{11} & \mathbf{q}_{12} & \mathbf{q}_{13} \end{pmatrix} \\ \begin{pmatrix} \mathbf{q}_{21} & \mathbf{q}_{22} & \mathbf{q}_{23} \end{pmatrix} \\ \begin{pmatrix} \mathbf{q}_{31} & \mathbf{q}_{32} & \mathbf{q}_{33} \end{pmatrix} \end{Bmatrix} = \begin{Bmatrix} \begin{pmatrix} \mathbf{p}_{11}^T \\ \mathbf{p}_{12}^T \\ \mathbf{p}_{13}^T \end{pmatrix} \\ \begin{pmatrix} \mathbf{p}_{21}^T \\ \mathbf{p}_{22}^T \\ \mathbf{p}_{23}^T \end{pmatrix} \\ \begin{pmatrix} \mathbf{p}_{31}^T \\ \mathbf{p}_{32}^T \\ \mathbf{p}_{33}^T \end{pmatrix} \end{Bmatrix}.
\end{aligned}
\tag{S7.1}
$$

The proofs for the higher dimensional cases are the same with the following proofs.

##### Proof of Eq. (S3.19)

$$
\begin{aligned}
\mathbf{u}^T \langle \mathbf{z}^T, \mathsf{Q} \rangle &= \mathbf{u}^T \left\langle \begin{pmatrix} z_1 & z_2 & z_3 \end{pmatrix}, \begin{Bmatrix} \mathbf{Q}^1 \\ \mathbf{Q}^2 \\ \mathbf{Q}^3 \end{Bmatrix} \right\rangle \\
&= \mathbf{u}^T [z_1 \mathbf{Q}^1 + z_2 \mathbf{Q}^2 + z_3 \mathbf{Q}^3] \\
&= z_1 \mathbf{u}^T \mathbf{Q}^1 + z_2 \mathbf{u}^T \mathbf{Q}^2 + z_3 \mathbf{u}^T \mathbf{Q}^3 \\
&= \begin{pmatrix} z_1 & z_2 & z_3 \end{pmatrix} \begin{pmatrix} \mathbf{u}^T \mathbf{Q}^1 \\ \mathbf{u}^T \mathbf{Q}^2 \\ \mathbf{u}^T \mathbf{Q}^3 \end{pmatrix} = \mathbf{z}^T \mathbf{u}^T \mathsf{Q}.
\end{aligned}
\tag{S7.2}
$$

##### Proof of Eq. (S3.20)

By using Eq. (S7.2),

$$\begin{aligned}
\mathbf{u}^T\langle\mathbf{B}^T,\mathsf{Q}\rangle &= \mathbf{u}^T\left\langle\begin{pmatrix}\mathbf{b}_1^T\\ \mathbf{b}_2^T\\ \mathbf{b}_3^T\end{pmatrix},\mathsf{Q}\right\rangle = \mathbf{u}^T\begin{Bmatrix}\langle\mathbf{b}_1^T,\mathsf{Q}\rangle\\ \langle\mathbf{b}_2^T,\mathsf{Q}\rangle\\ \langle\mathbf{b}_3^T,\mathsf{Q}\rangle\end{Bmatrix} = \begin{pmatrix}\mathbf{u}^T\langle\mathbf{b}_1^T,\mathsf{Q}\rangle\\ \mathbf{u}^T\langle\mathbf{b}_2^T,\mathsf{Q}\rangle\\ \mathbf{u}^T\langle\mathbf{b}_3^T,\mathsf{Q}\rangle\end{pmatrix}\\
&= \begin{pmatrix}\mathbf{b}_1^T\mathbf{u}^T\mathsf{Q}\\ \mathbf{b}_2^T\mathbf{u}^T\mathsf{Q}\\ \mathbf{b}_3^T\mathbf{u}^T\mathsf{Q}\end{pmatrix} = \begin{pmatrix}\mathbf{b}_1^T\\ \mathbf{b}_2^T\\ \mathbf{b}_3^T\end{pmatrix}\mathbf{u}^T\mathsf{Q} = \mathbf{B}^T\mathbf{u}^T\mathsf{Q}. \qquad \text{(S7.3)}
\end{aligned}$$

#### Proof of Eq. (S3.21)

$$\begin{aligned}
\mathbf{u}^T\mathsf{Q}^c &= \mathbf{u}^T\begin{Bmatrix}(\ \mathbf{q}_{11}\ \ \mathbf{q}_{12}\ \ \mathbf{q}_{13}\ )\\ (\ \mathbf{q}_{21}\ \ \mathbf{q}_{22}\ \ \mathbf{q}_{23}\ )\\ (\ \mathbf{q}_{31}\ \ \mathbf{q}_{32}\ \ \mathbf{q}_{33}\ )\end{Bmatrix}^c = \mathbf{u}^T\begin{Bmatrix}(\ \mathbf{q}_{11}\ \ \mathbf{q}_{21}\ \ \mathbf{q}_{31}\ )\\ (\ \mathbf{q}_{12}\ \ \mathbf{q}_{22}\ \ \mathbf{q}_{32}\ )\\ (\ \mathbf{q}_{13}\ \ \mathbf{q}_{23}\ \ \mathbf{q}_{33}\ )\end{Bmatrix}\\
&= \begin{pmatrix}\mathbf{u}^T(\ \mathbf{q}_{11}\ \ \mathbf{q}_{21}\ \ \mathbf{q}_{31}\ )\\ \mathbf{u}^T(\ \mathbf{q}_{12}\ \ \mathbf{q}_{22}\ \ \mathbf{q}_{32}\ )\\ \mathbf{u}^T(\ \mathbf{q}_{13}\ \ \mathbf{q}_{23}\ \ \mathbf{q}_{33}\ )\end{pmatrix} = \begin{pmatrix}\mathbf{u}^T\mathbf{q}_{11} & \mathbf{u}^T\mathbf{q}_{21} & \mathbf{u}^T\mathbf{q}_{31}\\ \mathbf{u}^T\mathbf{q}_{12} & \mathbf{u}^T\mathbf{q}_{22} & \mathbf{u}^T\mathbf{q}_{32}\\ \mathbf{u}^T\mathbf{q}_{13} & \mathbf{u}^T\mathbf{q}_{23} & \mathbf{u}^T\mathbf{q}_{33}\end{pmatrix}\\
&= \begin{pmatrix}\mathbf{u}^T\mathbf{q}_{11} & \mathbf{u}^T\mathbf{q}_{12} & \mathbf{u}^T\mathbf{q}_{13}\\ \mathbf{u}^T\mathbf{q}_{21} & \mathbf{u}^T\mathbf{q}_{22} & \mathbf{u}^T\mathbf{q}_{23}\\ \mathbf{u}^T\mathbf{q}_{31} & \mathbf{u}^T\mathbf{q}_{32} & \mathbf{u}^T\mathbf{q}_{33}\end{pmatrix}^T\\
&= \left[\mathbf{u}^T\begin{Bmatrix}(\ \mathbf{q}_{11}\ \ \mathbf{q}_{12}\ \ \mathbf{q}_{13}\ )\\ (\ \mathbf{q}_{21}\ \ \mathbf{q}_{22}\ \ \mathbf{q}_{23}\ )\\ (\ \mathbf{q}_{31}\ \ \mathbf{q}_{32}\ \ \mathbf{q}_{33}\ )\end{Bmatrix}\right]^T = [\mathbf{s}^T\mathsf{Q}]^T \qquad \text{(S7.4)}
\end{aligned}$$

#### Proof of Eq. (S3.22)

$$\begin{aligned}
\mathbf{u}^T\mathsf{Q}^r &= \mathbf{u}^T\begin{Bmatrix}\begin{pmatrix}\mathbf{p}_{11}^T\\ \mathbf{p}_{12}^T\\ \mathbf{p}_{13}^T\end{pmatrix}\\ \begin{pmatrix}\mathbf{p}_{21}^T\\ \mathbf{p}_{22}^T\\ \mathbf{p}_{23}^T\end{pmatrix}\\ \begin{pmatrix}\mathbf{p}_{31}^T\\ \mathbf{p}_{32}^T\\ \mathbf{p}_{33}^T\end{pmatrix}\end{Bmatrix}^r = \mathbf{u}^T\begin{Bmatrix}\begin{pmatrix}\mathbf{p}_{11}^T\\ \mathbf{p}_{21}^T\\ \mathbf{p}_{31}^T\end{pmatrix}\\ \begin{pmatrix}\mathbf{p}_{12}^T\\ \mathbf{p}_{22}^T\\ \mathbf{p}_{32}^T\end{pmatrix}\\ \begin{pmatrix}\mathbf{p}_{13}^T\\ \mathbf{p}_{23}^T\\ \mathbf{p}_{33}^T\end{pmatrix}\end{Bmatrix} = \begin{pmatrix}\mathbf{u}^T\begin{pmatrix}\mathbf{p}_{11}^T\\ \mathbf{p}_{21}^T\\ \mathbf{p}_{31}^T\end{pmatrix}\\ \mathbf{u}^T\begin{pmatrix}\mathbf{p}_{12}^T\\ \mathbf{p}_{22}^T\\ \mathbf{p}_{32}^T\end{pmatrix}\\ \mathbf{u}^T\begin{pmatrix}\mathbf{p}_{13}^T\\ \mathbf{p}_{23}^T\\ \mathbf{p}_{33}^T\end{pmatrix}\end{pmatrix}\\
&= \begin{pmatrix}u_1\mathbf{p}_{11}^T + u_2\mathbf{p}_{21}^T + u_3\mathbf{p}_{31}^T\\ u_1\mathbf{p}_{12}^T + u_2\mathbf{p}_{22}^T + u_3\mathbf{p}_{32}^T\\ u_1\mathbf{p}_{13}^T + u_2\mathbf{p}_{23}^T + u_3\mathbf{p}_{33}^T\end{pmatrix}\\
&= u_1\begin{pmatrix}\mathbf{p}_{11}^T\\ \mathbf{p}_{12}^T\\ \mathbf{p}_{13}^T\end{pmatrix} + u_2\begin{pmatrix}\mathbf{p}_{21}^T\\ \mathbf{p}_{22}^T\\ \mathbf{p}_{23}^T\end{pmatrix} + u_3\begin{pmatrix}\mathbf{p}_{31}^T\\ \mathbf{p}_{32}^T\\ \mathbf{p}_{33}^T\end{pmatrix}\\
&= \left\langle(\ u_1\ \ u_2\ \ u_3\ ),\begin{Bmatrix}\begin{pmatrix}\mathbf{p}_{11}^T\\ \mathbf{p}_{12}^T\\ \mathbf{p}_{13}^T\end{pmatrix}\\ \begin{pmatrix}\mathbf{p}_{21}^T\\ \mathbf{p}_{22}^T\\ \mathbf{p}_{23}^T\end{pmatrix}\\ \begin{pmatrix}\mathbf{p}_{31}^T\\ \mathbf{p}_{32}^T\\ \mathbf{p}_{33}^T\end{pmatrix}\end{Bmatrix}\right\rangle = \langle\mathbf{u}^T,\mathsf{Q}\rangle. \qquad \text{(S7.5)}
\end{aligned}$$

#### Proof of Eq. (S3.23)

$$\begin{aligned}\mathbf{B}\mathbf{Q}^c &= \mathbf{B}\begin{Bmatrix}(\,\mathbf{q}_{11}\;\;\mathbf{q}_{12}\;\;\mathbf{q}_{13}\,)\\(\,\mathbf{q}_{21}\;\;\mathbf{q}_{22}\;\;\mathbf{q}_{23}\,)\\(\,\mathbf{q}_{31}\;\;\mathbf{q}_{32}\;\;\mathbf{q}_{33}\,)\end{Bmatrix}^c = \mathbf{B}\begin{Bmatrix}(\,\mathbf{q}_{11}\;\;\mathbf{q}_{21}\;\;\mathbf{q}_{31}\,)\\(\,\mathbf{q}_{12}\;\;\mathbf{q}_{22}\;\;\mathbf{q}_{32}\,)\\(\,\mathbf{q}_{13}\;\;\mathbf{q}_{23}\;\;\mathbf{q}_{33}\,)\end{Bmatrix} = \begin{Bmatrix}(\,\mathbf{B}\mathbf{q}_{11}\;\;\mathbf{B}\mathbf{q}_{21}\;\;\mathbf{B}\mathbf{q}_{31}\,)\\(\,\mathbf{B}\mathbf{q}_{12}\;\;\mathbf{B}\mathbf{q}_{22}\;\;\mathbf{B}\mathbf{q}_{32}\,)\\(\,\mathbf{B}\mathbf{q}_{13}\;\;\mathbf{B}\mathbf{q}_{23}\;\;\mathbf{B}\mathbf{q}_{33}\,)\end{Bmatrix}\\ &= \begin{Bmatrix}(\,\mathbf{B}\mathbf{q}_{11}\;\;\mathbf{B}\mathbf{q}_{12}\;\;\mathbf{B}\mathbf{q}_{13}\,)\\(\,\mathbf{B}\mathbf{q}_{21}\;\;\mathbf{B}\mathbf{q}_{22}\;\;\mathbf{B}\mathbf{q}_{23}\,)\\(\,\mathbf{B}\mathbf{q}_{31}\;\;\mathbf{B}\mathbf{q}_{32}\;\;\mathbf{B}\mathbf{q}_{33}\,)\end{Bmatrix}^c = \left[\mathbf{B}\begin{Bmatrix}(\,\mathbf{q}_{11}\;\;\mathbf{q}_{12}\;\;\mathbf{q}_{13}\,)\\(\,\mathbf{q}_{21}\;\;\mathbf{q}_{22}\;\;\mathbf{q}_{23}\,)\\(\,\mathbf{q}_{31}\;\;\mathbf{q}_{32}\;\;\mathbf{q}_{33}\,)\end{Bmatrix}\right]^c = (\mathbf{B}\mathbf{Q})^c \qquad \text{(S7.6)}\end{aligned}$$

#### Proof of Eq. (S3.24)

$$\begin{aligned}\mathbf{Q}^r\mathbf{B} &= \begin{Bmatrix}\begin{pmatrix}\mathbf{p}_{11}^T\\\mathbf{p}_{12}^T\\\mathbf{p}_{13}^T\end{pmatrix}\\\begin{pmatrix}\mathbf{p}_{21}^T\\\mathbf{p}_{22}^T\\\mathbf{p}_{23}^T\end{pmatrix}\\\begin{pmatrix}\mathbf{p}_{31}^T\\\mathbf{p}_{32}^T\\\mathbf{p}_{33}^T\end{pmatrix}\end{Bmatrix}^r\mathbf{B} = \begin{Bmatrix}\begin{pmatrix}\mathbf{p}_{11}^T\\\mathbf{p}_{21}^T\\\mathbf{p}_{31}^T\end{pmatrix}\\\begin{pmatrix}\mathbf{p}_{12}^T\\\mathbf{p}_{22}^T\\\mathbf{p}_{32}^T\end{pmatrix}\\\begin{pmatrix}\mathbf{p}_{13}^T\\\mathbf{p}_{23}^T\\\mathbf{p}_{33}^T\end{pmatrix}\end{Bmatrix}\mathbf{B} = \begin{Bmatrix}\begin{pmatrix}\mathbf{p}_{11}^T\\\mathbf{p}_{21}^T\\\mathbf{p}_{31}^T\end{pmatrix}\mathbf{B}\\\begin{pmatrix}\mathbf{p}_{12}^T\\\mathbf{p}_{22}^T\\\mathbf{p}_{32}^T\end{pmatrix}\mathbf{B}\\\begin{pmatrix}\mathbf{p}_{13}^T\\\mathbf{p}_{23}^T\\\mathbf{p}_{33}^T\end{pmatrix}\mathbf{B}\end{Bmatrix}\\ &= \begin{Bmatrix}\begin{pmatrix}\mathbf{p}_{11}^T\mathbf{B}\\\mathbf{p}_{21}^T\mathbf{B}\\\mathbf{p}_{31}^T\mathbf{B}\end{pmatrix}\\\begin{pmatrix}\mathbf{p}_{12}^T\mathbf{B}\\\mathbf{p}_{22}^T\mathbf{B}\\\mathbf{p}_{32}^T\mathbf{B}\end{pmatrix}\\\begin{pmatrix}\mathbf{p}_{13}^T\mathbf{B}\\\mathbf{p}_{23}^T\mathbf{B}\\\mathbf{p}_{33}^T\mathbf{B}\end{pmatrix}\end{Bmatrix} = \begin{Bmatrix}\begin{pmatrix}\mathbf{p}_{11}^T\mathbf{B}\\\mathbf{p}_{12}^T\mathbf{B}\\\mathbf{p}_{13}^T\mathbf{B}\end{pmatrix}\\\begin{pmatrix}\mathbf{p}_{21}^T\mathbf{B}\\\mathbf{p}_{22}^T\mathbf{B}\\\mathbf{p}_{23}^T\mathbf{B}\end{pmatrix}\\\begin{pmatrix}\mathbf{p}_{31}^T\mathbf{B}\\\mathbf{p}_{32}^T\mathbf{B}\\\mathbf{p}_{33}^T\mathbf{B}\end{pmatrix}\end{Bmatrix}^r = (\mathbf{Q}\mathbf{B})^r \qquad \text{(S7.7)}\end{aligned}$$

#### Proof of Eq. (S3.25)

$$\begin{aligned}\mathbf{B}\mathbf{Q}^r &= \mathbf{B}\begin{Bmatrix}\begin{pmatrix}\mathbf{p}_{11}^T\\\mathbf{p}_{12}^T\\\mathbf{p}_{13}^T\end{pmatrix}\\\begin{pmatrix}\mathbf{p}_{21}^T\\\mathbf{p}_{22}^T\\\mathbf{p}_{23}^T\end{pmatrix}\\\begin{pmatrix}\mathbf{p}_{31}^T\\\mathbf{p}_{32}^T\\\mathbf{p}_{33}^T\end{pmatrix}\end{Bmatrix}^r = \mathbf{B}\begin{Bmatrix}\begin{pmatrix}\mathbf{p}_{11}^T\\\mathbf{p}_{21}^T\\\mathbf{p}_{31}^T\end{pmatrix}\\\begin{pmatrix}\mathbf{p}_{12}^T\\\mathbf{p}_{22}^T\\\mathbf{p}_{32}^T\end{pmatrix}\\\begin{pmatrix}\mathbf{p}_{13}^T\\\mathbf{p}_{23}^T\\\mathbf{p}_{33}^T\end{pmatrix}\end{Bmatrix} = \begin{Bmatrix}\begin{pmatrix}B_{11}&B_{12}&B_{13}\\B_{21}&B_{22}&B_{23}\\B_{31}&B_{32}&B_{33}\end{pmatrix}\begin{pmatrix}\mathbf{p}_{11}^T\\\mathbf{p}_{21}^T\\\mathbf{p}_{31}^T\end{pmatrix}\\\begin{pmatrix}B_{11}&B_{12}&B_{13}\\B_{21}&B_{22}&B_{23}\\B_{31}&B_{32}&B_{33}\end{pmatrix}\begin{pmatrix}\mathbf{p}_{12}^T\\\mathbf{p}_{22}^T\\\mathbf{p}_{32}^T\end{pmatrix}\\\begin{pmatrix}B_{11}&B_{12}&B_{13}\\B_{21}&B_{22}&B_{23}\\B_{31}&B_{32}&B_{33}\end{pmatrix}\begin{pmatrix}\mathbf{p}_{13}^T\\\mathbf{p}_{23}^T\\\mathbf{p}_{33}^T\end{pmatrix}\end{Bmatrix}\\ &= \begin{Bmatrix}\begin{pmatrix}B_{11}\mathbf{p}_{11}^T+B_{12}\mathbf{p}_{21}^T+B_{13}\mathbf{p}_{31}^T\\B_{21}\mathbf{p}_{11}^T+B_{22}\mathbf{p}_{21}^T+B_{23}\mathbf{p}_{31}^T\\B_{31}\mathbf{p}_{11}^T+B_{32}\mathbf{p}_{21}^T+B_{33}\mathbf{p}_{31}^T\end{pmatrix}\\\begin{pmatrix}B_{11}\mathbf{p}_{12}^T+B_{12}\mathbf{p}_{22}^T+B_{13}\mathbf{p}_{32}^T\\B_{21}\mathbf{p}_{12}^T+B_{22}\mathbf{p}_{22}^T+B_{23}\mathbf{p}_{32}^T\\B_{31}\mathbf{p}_{12}^T+B_{32}\mathbf{p}_{22}^T+B_{33}\mathbf{p}_{32}^T\end{pmatrix}\\\begin{pmatrix}B_{11}\mathbf{p}_{13}^T+B_{12}\mathbf{p}_{23}^T+B_{13}\mathbf{p}_{33}^T\\B_{21}\mathbf{p}_{13}^T+B_{22}\mathbf{p}_{23}^T+B_{23}\mathbf{p}_{33}^T\\B_{31}\mathbf{p}_{13}^T+B_{32}\mathbf{p}_{23}^T+B_{33}\mathbf{p}_{33}^T\end{pmatrix}\end{Bmatrix} = \begin{Bmatrix}\begin{pmatrix}B_{11}\mathbf{p}_{11}^T+B_{12}\mathbf{p}_{21}^T+B_{13}\mathbf{p}_{31}^T\\B_{11}\mathbf{p}_{12}^T+B_{12}\mathbf{p}_{22}^T+B_{13}\mathbf{p}_{32}^T\\B_{11}\mathbf{p}_{13}^T+B_{12}\mathbf{p}_{23}^T+B_{13}\mathbf{p}_{33}^T\end{pmatrix}\\\begin{pmatrix}B_{21}\mathbf{p}_{11}^T+B_{22}\mathbf{p}_{21}^T+B_{23}\mathbf{p}_{31}^T\\B_{21}\mathbf{p}_{12}^T+B_{22}\mathbf{p}_{22}^T+B_{23}\mathbf{p}_{32}^T\\B_{21}\mathbf{p}_{13}^T+B_{22}\mathbf{p}_{23}^T+B_{23}\mathbf{p}_{33}^T\end{pmatrix}\\\begin{pmatrix}B_{31}\mathbf{p}_{11}^T+B_{32}\mathbf{p}_{21}^T+B_{33}\mathbf{p}_{31}^T\\B_{31}\mathbf{p}_{12}^T+B_{32}\mathbf{p}_{22}^T+B_{33}\mathbf{p}_{32}^T\\B_{31}\mathbf{p}_{13}^T+B_{32}\mathbf{p}_{23}^T+B_{33}\mathbf{p}_{33}^T\end{pmatrix}\end{Bmatrix}^r\end{aligned}$$

$$
= \left\{ \begin{array}{c}
B_{11}\begin{pmatrix} \mathbf{p}_{11}^T \\ \mathbf{p}_{12}^T \\ \mathbf{p}_{13}^T \end{pmatrix} + B_{12}\begin{pmatrix} \mathbf{p}_{21}^T \\ \mathbf{p}_{22}^T \\ \mathbf{p}_{23}^T \end{pmatrix} + B_{13}\begin{pmatrix} \mathbf{p}_{31}^T \\ \mathbf{p}_{32}^T \\ \mathbf{p}_{33}^T \end{pmatrix} \\
B_{21}\begin{pmatrix} \mathbf{p}_{11}^T \\ \mathbf{p}_{12}^T \\ \mathbf{p}_{13}^T \end{pmatrix} + B_{22}\begin{pmatrix} \mathbf{p}_{21}^T \\ \mathbf{p}_{22}^T \\ \mathbf{p}_{23}^T \end{pmatrix} + B_{23}\begin{pmatrix} \mathbf{p}_{31}^T \\ \mathbf{p}_{32}^T \\ \mathbf{p}_{33}^T \end{pmatrix} \\
B_{31}\begin{pmatrix} \mathbf{p}_{11}^T \\ \mathbf{p}_{12}^T \\ \mathbf{p}_{13}^T \end{pmatrix} + B_{32}\begin{pmatrix} \mathbf{p}_{21}^T \\ \mathbf{p}_{22}^T \\ \mathbf{p}_{23}^T \end{pmatrix} + B_{33}\begin{pmatrix} \mathbf{p}_{31}^T \\ \mathbf{p}_{32}^T \\ \mathbf{p}_{33}^T \end{pmatrix}
\end{array} \right\}^r
$$

$$
= \left\{ \begin{array}{c}
\langle (\ B_{11} \ \ B_{12} \ \ B_{13}\ ), \mathsf{Q} \rangle \\
\langle (\ B_{21} \ \ B_{22} \ \ B_{23}\ ), \mathsf{Q} \rangle \\
\langle (\ B_{31} \ \ B_{32} \ \ B_{33}\ ), \mathsf{Q} \rangle
\end{array} \right\}^r = \left\langle \begin{pmatrix} B_{11} & B_{12} & B_{13} \\ B_{21} & B_{22} & B_{23} \\ B_{31} & B_{32} & B_{33} \end{pmatrix}, \mathsf{Q} \right\rangle^r = \langle \mathbf{B}, \mathsf{Q} \rangle^r \qquad \text{(S7.8)}
$$

#### Proof of Eq. (S3.26)

$$
\begin{aligned}
\langle \mathbf{B}^T, \mathsf{Q}^c \rangle &= \left\langle \begin{pmatrix} B_{11} & B_{21} & B_{31} \\ B_{12} & B_{22} & B_{32} \\ B_{13} & B_{23} & B_{33} \end{pmatrix}, \mathsf{Q}^c \right\rangle \\
&= \left\{ \begin{array}{c}
\langle (\ B_{11} \ \ B_{21} \ \ B_{31}\ ), \mathsf{Q}^c \rangle \\
\langle (\ B_{12} \ \ B_{22} \ \ B_{32}\ ), \mathsf{Q}^c \rangle \\
\langle (\ B_{13} \ \ B_{23} \ \ B_{33}\ ), \mathsf{Q}^c \rangle
\end{array} \right\} \\
&= \left\{ \begin{array}{c}
\left\langle (\ B_{11} \ \ B_{21} \ \ B_{31}\ ), \left\{ \begin{array}{c} (\ \mathbf{q}_{11} \ \ \mathbf{q}_{21} \ \ \mathbf{q}_{31}\ ) \\ (\ \mathbf{q}_{12} \ \ \mathbf{q}_{22} \ \ \mathbf{q}_{32}\ ) \\ (\ \mathbf{q}_{13} \ \ \mathbf{q}_{23} \ \ \mathbf{q}_{33}\ ) \end{array} \right\} \right\rangle \\
\left\langle (\ B_{12} \ \ B_{22} \ \ B_{32}\ ), \left\{ \begin{array}{c} (\ \mathbf{q}_{11} \ \ \mathbf{q}_{21} \ \ \mathbf{q}_{31}\ ) \\ (\ \mathbf{q}_{12} \ \ \mathbf{q}_{22} \ \ \mathbf{q}_{32}\ ) \\ (\ \mathbf{q}_{13} \ \ \mathbf{q}_{23} \ \ \mathbf{q}_{33}\ ) \end{array} \right\} \right\rangle \\
\left\langle (\ B_{13} \ \ B_{23} \ \ B_{33}\ ), \left\{ \begin{array}{c} (\ \mathbf{q}_{11} \ \ \mathbf{q}_{21} \ \ \mathbf{q}_{31}\ ) \\ (\ \mathbf{q}_{12} \ \ \mathbf{q}_{22} \ \ \mathbf{q}_{32}\ ) \\ (\ \mathbf{q}_{13} \ \ \mathbf{q}_{23} \ \ \mathbf{q}_{33}\ ) \end{array} \right\} \right\rangle
\end{array} \right\} \\
&= \left\{ \begin{array}{l}
B_{11}(\ \mathbf{q}_{11} \ \ \mathbf{q}_{21} \ \ \mathbf{q}_{31}\ ) + B_{21}(\ \mathbf{q}_{12} \ \ \mathbf{q}_{22} \ \ \mathbf{q}_{32}\ ) + B_{31}(\ \mathbf{q}_{13} \ \ \mathbf{q}_{23} \ \ \mathbf{q}_{33}\ ) \\
B_{12}(\ \mathbf{q}_{11} \ \ \mathbf{q}_{21} \ \ \mathbf{q}_{31}\ ) + B_{22}(\ \mathbf{q}_{12} \ \ \mathbf{q}_{22} \ \ \mathbf{q}_{32}\ ) + B_{32}(\ \mathbf{q}_{13} \ \ \mathbf{q}_{23} \ \ \mathbf{q}_{33}\ ) \\
B_{13}(\ \mathbf{q}_{11} \ \ \mathbf{q}_{21} \ \ \mathbf{q}_{31}\ ) + B_{23}(\ \mathbf{q}_{12} \ \ \mathbf{q}_{22} \ \ \mathbf{q}_{32}\ ) + B_{33}(\ \mathbf{q}_{13} \ \ \mathbf{q}_{23} \ \ \mathbf{q}_{33}\ )
\end{array} \right\} \\
&= \left\{ \begin{array}{l}
(\ B_{11}\mathbf{q}_{11} + B_{21}\mathbf{q}_{12} + B_{31}\mathbf{q}_{13}, \ \ B_{11}\mathbf{q}_{21} + B_{21}\mathbf{q}_{22} + B_{31}\mathbf{q}_{23}, \ \ B_{11}\mathbf{q}_{31} + B_{21}\mathbf{q}_{32} + B_{31}\mathbf{q}_{33}\ ) \\
(\ B_{12}\mathbf{q}_{11} + B_{22}\mathbf{q}_{12} + B_{32}\mathbf{q}_{13}, \ \ B_{12}\mathbf{q}_{21} + B_{22}\mathbf{q}_{22} + B_{32}\mathbf{q}_{23}, \ \ B_{12}\mathbf{q}_{31} + B_{22}\mathbf{q}_{32} + B_{32}\mathbf{q}_{33}\ ) \\
(\ B_{13}\mathbf{q}_{11} + B_{23}\mathbf{q}_{12} + B_{33}\mathbf{q}_{13}, \ \ B_{13}\mathbf{q}_{21} + B_{23}\mathbf{q}_{22} + B_{33}\mathbf{q}_{23}, \ \ B_{13}\mathbf{q}_{31} + B_{23}\mathbf{q}_{32} + B_{33}\mathbf{q}_{33}\ )
\end{array} \right\} \\
&= \left\{ \begin{array}{l}
(\ B_{11}\mathbf{q}_{11} + B_{21}\mathbf{q}_{12} + B_{31}\mathbf{q}_{13}, \ \ B_{12}\mathbf{q}_{11} + B_{22}\mathbf{q}_{12} + B_{32}\mathbf{q}_{13}, \ \ B_{13}\mathbf{q}_{11} + B_{23}\mathbf{q}_{12} + B_{33}\mathbf{q}_{13},\ ) \\
(\ B_{11}\mathbf{q}_{21} + B_{21}\mathbf{q}_{22} + B_{31}\mathbf{q}_{23}, \ \ B_{12}\mathbf{q}_{21} + B_{22}\mathbf{q}_{22} + B_{32}\mathbf{q}_{23}, \ \ B_{13}\mathbf{q}_{21} + B_{23}\mathbf{q}_{22} + B_{33}\mathbf{q}_{23}\ ) \\
(\ B_{11}\mathbf{q}_{31} + B_{21}\mathbf{q}_{32} + B_{31}\mathbf{q}_{23}, \ \ B_{12}\mathbf{q}_{31} + B_{22}\mathbf{q}_{32} + B_{32}\mathbf{q}_{33} \ \ B_{13}\mathbf{q}_{31} + B_{23}\mathbf{q}_{32} + B_{33}\mathbf{q}_{33}\ )
\end{array} \right\}^c \\
&= \left\{ \begin{array}{c}
(\ \mathbf{q}_{11} \ \ \mathbf{q}_{12} \ \ \mathbf{q}_{13}\ )\begin{pmatrix} B_{11} & B_{12} & B_{13} \\ B_{21} & B_{22} & B_{23} \\ B_{31} & B_{32} & B_{33} \end{pmatrix} \\
(\ \mathbf{q}_{21} \ \ \mathbf{q}_{22} \ \ \mathbf{q}_{23}\ )\begin{pmatrix} B_{11} & B_{12} & B_{13} \\ B_{21} & B_{22} & B_{23} \\ B_{31} & B_{32} & B_{33} \end{pmatrix} \\
(\ \mathbf{q}_{31} \ \ \mathbf{q}_{32} \ \ \mathbf{q}_{33}\ )\begin{pmatrix} B_{11} & B_{12} & B_{13} \\ B_{21} & B_{22} & B_{23} \\ B_{31} & B_{32} & B_{33} \end{pmatrix}
\end{array} \right\}^c \\
&= \left[ \left\{ \begin{array}{c} (\ \mathbf{q}_{11} \ \ \mathbf{q}_{12} \ \ \mathbf{q}_{13}\ ) \\ (\ \mathbf{q}_{21} \ \ \mathbf{q}_{22} \ \ \mathbf{q}_{23}\ ) \\ (\ \mathbf{q}_{31} \ \ \mathbf{q}_{32} \ \ \mathbf{q}_{33}\ ) \end{array} \right\} \begin{pmatrix} B_{11} & B_{12} & B_{13} \\ B_{21} & B_{22} & B_{23} \\ B_{31} & B_{32} & B_{33} \end{pmatrix} \right]^c = (\mathsf{Q}\mathbf{B})^c \qquad \text{(S7.9)}
\end{aligned}
$$

**Proof of Eq. (S3.27)**

$$\begin{aligned}
\mathbf{A}^T\langle \mathbf{u}^T, \mathsf{Q}\rangle &= \begin{pmatrix} \mathbf{a}_1^T \\ \mathbf{a}_2^T \\ \mathbf{a}_3^T \end{pmatrix} \left\langle \begin{pmatrix} u_1 & u_2 & u_3 \end{pmatrix}, \begin{pmatrix} \mathbf{Q}^1 \\ \mathbf{Q}^2 \\ \mathbf{Q}^3 \end{pmatrix} \right\rangle \\
&= \begin{pmatrix} \mathbf{a}_1^T \\ \mathbf{a}_2^T \\ \mathbf{a}_3^T \end{pmatrix} (u_1\mathbf{Q}^1 + u_2\mathbf{Q}^2 + u_3\mathbf{Q}^3) \\
&= \begin{pmatrix} \mathbf{a}_1^T(u_1\mathbf{Q}^1 + u_2\mathbf{Q}^2 + u_3\mathbf{Q}^3) \\ \mathbf{a}_2^T(u_1\mathbf{Q}^1 + u_2\mathbf{Q}^2 + u_3\mathbf{Q}^3) \\ \mathbf{a}_3^T(u_1\mathbf{Q}^1 + u_2\mathbf{Q}^2 + u_3\mathbf{Q}^3) \end{pmatrix} \\
&= u_1\begin{pmatrix} \mathbf{a}_1^T\mathbf{Q}^1 \\ \mathbf{a}_2^T\mathbf{Q}^1 \\ \mathbf{a}_3^T\mathbf{Q}^1 \end{pmatrix} + u_2\begin{pmatrix} \mathbf{a}_1^T\mathbf{Q}^2 \\ \mathbf{a}_2^T\mathbf{Q}^2 \\ \mathbf{a}_3^T\mathbf{Q}^2 \end{pmatrix} + u_3\begin{pmatrix} \mathbf{a}_1^T\mathbf{Q}^3 \\ \mathbf{a}_2^T\mathbf{Q}^3 \\ \mathbf{a}_3^T\mathbf{Q}^3 \end{pmatrix} \\
&= \left\langle \mathbf{u}^T, \left\{ \begin{matrix} \begin{pmatrix} \mathbf{a}_1^T\mathbf{Q}^1 \\ \mathbf{a}_2^T\mathbf{Q}^1 \\ \mathbf{a}_3^T\mathbf{Q}^1 \end{pmatrix} \\ \begin{pmatrix} \mathbf{a}_1^T\mathbf{Q}^2 \\ \mathbf{a}_2^T\mathbf{Q}^2 \\ \mathbf{a}_3^T\mathbf{Q}^2 \end{pmatrix} \\ \begin{pmatrix} \mathbf{a}_1^T\mathbf{Q}^3 \\ \mathbf{a}_2^T\mathbf{Q}^3 \\ \mathbf{a}_3^T\mathbf{Q}^3 \end{pmatrix} \end{matrix} \right\} \right\rangle = \left\langle \mathbf{u}^T, \left\{ \begin{matrix} \begin{pmatrix} \mathbf{a}_1^T \\ \mathbf{a}_2^T \\ \mathbf{a}_3^T \end{pmatrix}\mathbf{Q}^1 \\ \begin{pmatrix} \mathbf{a}_1^T \\ \mathbf{a}_2^T \\ \mathbf{a}_3^T \end{pmatrix}\mathbf{Q}^2 \\ \begin{pmatrix} \mathbf{a}_1^T \\ \mathbf{a}_2^T \\ \mathbf{a}_3^T \end{pmatrix}\mathbf{Q}^3 \end{matrix} \right\} \right\rangle \\
&= \left\langle \mathbf{u}^T, \begin{pmatrix} \mathbf{a}_1^T \\ \mathbf{a}_2^T \\ \mathbf{a}_3^T \end{pmatrix} \begin{pmatrix} \mathbf{Q}^1 \\ \mathbf{Q}^2 \\ \mathbf{Q}^3 \end{pmatrix} \right\rangle \\
&= \langle \mathbf{u}^T, \mathbf{A}^T\mathsf{Q}\rangle \qquad \text{(S7.10)}
\end{aligned}$$

and

$$\begin{aligned}
\langle \mathbf{u}^T, \mathsf{Q}\rangle\mathbf{A} &= \left\langle \begin{pmatrix} u_1 & u_2 & u_3 \end{pmatrix}, \begin{pmatrix} \mathbf{Q}_1 \\ \mathbf{Q}_2 \\ \mathbf{Q}_3 \end{pmatrix} \right\rangle \mathbf{A} \\
&= (u_1\mathbf{Q}_1 + u_2\mathbf{Q}_2 + u_3\mathbf{Q}_3)\mathbf{A} \\
&= u_1\mathbf{Q}_1\mathbf{A} + u_2\mathbf{Q}_2\mathbf{A} + u_3\mathbf{Q}_3\mathbf{A} \\
&= \left\langle \begin{pmatrix} u_1 & u_2 & u_3 \end{pmatrix}, \begin{pmatrix} \mathbf{Q}_1\mathbf{A} \\ \mathbf{Q}_2\mathbf{A} \\ \mathbf{Q}_3\mathbf{A} \end{pmatrix} \right\rangle \\
&= \langle \mathbf{u}^T, \mathsf{Q}\mathbf{A}\rangle. \qquad \text{(S7.11)}
\end{aligned}$$

**Proof of Eq. (S3.28)**

By using Eqs. (S7.10) and (S7.11),

$$\mathbf{A}\langle \mathbf{B}, \mathsf{Q}\rangle = \mathbf{A}\left\langle \begin{pmatrix} \mathbf{b}_1^T \\ \mathbf{b}_2^T \\ \mathbf{b}_3^T \end{pmatrix}, \mathsf{Q} \right\rangle = \mathbf{A}\left\{ \begin{matrix} \langle \mathbf{b}_1^T, \mathsf{Q}\rangle \\ \langle \mathbf{b}_2^T, \mathsf{Q}\rangle \\ \langle \mathbf{b}_3^T, \mathsf{Q}\rangle \end{matrix} \right\} = \left\{ \begin{matrix} \mathbf{A}\langle \mathbf{b}_1^T, \mathsf{Q}\rangle \\ \mathbf{A}\langle \mathbf{b}_2^T, \mathsf{Q}\rangle \\ \mathbf{A}\langle \mathbf{b}_3^T, \mathsf{Q}\rangle \end{matrix} \right\}$$

$$= \left\{ \begin{array}{c} \langle \mathbf{b}_1^T, \mathbf{A}\mathrm{Q} \rangle \\ \langle \mathbf{b}_2^T, \mathbf{A}\mathrm{Q} \rangle \\ \langle \mathbf{b}_3^T, \mathbf{A}\mathrm{Q} \rangle \end{array} \right\} = \left\langle \begin{pmatrix} \mathbf{b}_1^T \\ \mathbf{b}_2^T \\ \mathbf{b}_3^T \end{pmatrix}, \mathbf{A}\mathrm{Q} \right\rangle = \langle \mathbf{B}, \mathbf{A}\mathrm{Q} \rangle. \quad \text{(S7.12)}$$

$$\begin{aligned} \langle \mathbf{B}, \mathrm{Q} \rangle \mathbf{A} &= \left\langle \begin{pmatrix} \mathbf{b}_1^T \\ \mathbf{b}_2^T \\ \mathbf{b}_3^T \end{pmatrix}, \mathrm{Q} \right\rangle \mathbf{A} = \left\{ \begin{array}{c} \langle \mathbf{b}_1^T, \mathrm{Q} \rangle \\ \langle \mathbf{b}_2^T, \mathrm{Q} \rangle \\ \langle \mathbf{b}_3^T, \mathrm{Q} \rangle \end{array} \right\} \mathbf{A} = \left\{ \begin{array}{c} \langle \mathbf{b}_1^T, \mathrm{Q} \rangle \mathbf{A} \\ \langle \mathbf{b}_2^T, \mathrm{Q} \rangle \mathbf{A} \\ \langle \mathbf{b}_3^T, \mathrm{Q} \rangle \mathbf{A} \end{array} \right\} \\ &= \left\{ \begin{array}{c} \langle \mathbf{b}_1^T, \mathrm{Q}\mathbf{A} \rangle \\ \langle \mathbf{b}_2^T, \mathrm{Q}\mathbf{A} \rangle \\ \langle \mathbf{b}_3^T, \mathrm{Q}\mathbf{A} \rangle \end{array} \right\} = \left\langle \begin{pmatrix} \mathbf{b}_1^T \\ \mathbf{b}_2^T \\ \mathbf{b}_3^T \end{pmatrix}, \mathrm{Q}\mathbf{A} \right\rangle = \langle \mathbf{B}, \mathrm{Q}\mathbf{A} \rangle. \end{aligned} \quad \text{(S7.13)}$$

**Proof of Eq. (S3.29)**

First we see

$$\begin{aligned} \langle \mathbf{u}^T, \langle \mathbf{A}, \mathrm{Q} \rangle \rangle &= \left\langle \begin{pmatrix} u_1 & u_2 & u_3 \end{pmatrix}, \left\langle \begin{pmatrix} \mathbf{a}_1^T \\ \mathbf{a}_2^T \\ \mathbf{a}_3^T \end{pmatrix}, \mathrm{Q} \right\rangle \right\rangle \\ &= \left\langle \begin{pmatrix} u_1 & u_2 & u_3 \end{pmatrix}, \left\{ \begin{array}{c} \langle \mathbf{a}_1^T, \mathrm{Q} \rangle \\ \langle \mathbf{a}_2^T, \mathrm{Q} \rangle \\ \langle \mathbf{a}_3^T, \mathrm{Q} \rangle \end{array} \right\} \right\rangle \\ &= u_1 \langle \mathbf{a}_1^T, \mathrm{Q} \rangle + u_2 \langle \mathbf{a}_2^T, \mathrm{Q} \rangle + u_3 \langle \mathbf{a}_3^T, \mathrm{Q} \rangle \\ &= \langle u_1 \mathbf{a}_1^T, \mathrm{Q} \rangle + \langle u_2 \mathbf{a}_2^T, \mathrm{Q} \rangle + \langle u_3 \mathbf{a}_3^T, \mathrm{Q} \rangle \\ &= \langle u_1 \mathbf{a}_1^T + u_2 \mathbf{a}_2^T + u_3 \mathbf{a}_3^T, \mathrm{Q} \rangle \\ &= \left\langle \begin{pmatrix} u_1 & u_2 & u_3 \end{pmatrix} \begin{pmatrix} \mathbf{a}_1^T \\ \mathbf{a}_2^T \\ \mathbf{a}_3^T \end{pmatrix}, \mathrm{Q} \right\rangle \\ &= \langle \mathbf{u}^T \mathbf{A}, \mathrm{Q} \rangle, \end{aligned} \quad \text{(S7.14)}$$

and thus we see

$$\begin{aligned} \langle \mathbf{B}, \langle \mathbf{A}, \mathrm{Q} \rangle \rangle &= \left\langle \begin{pmatrix} \mathbf{b}_1^T \\ \mathbf{b}_2^T \\ \mathbf{b}_3^T \end{pmatrix}, \langle \mathbf{A}, \mathrm{Q} \rangle \right\rangle \\ &= \left\{ \begin{array}{c} \langle \mathbf{b}_1^T, \langle \mathbf{A}, \mathrm{Q} \rangle \rangle \\ \langle \mathbf{b}_2^T, \langle \mathbf{A}, \mathrm{Q} \rangle \rangle \\ \langle \mathbf{b}_3^T, \langle \mathbf{A}, \mathrm{Q} \rangle \rangle \end{array} \right\} = \left\{ \begin{array}{c} \langle \mathbf{b}_1^T \mathbf{A}, \mathrm{Q} \rangle \\ \langle \mathbf{b}_2^T \mathbf{A}, \mathrm{Q} \rangle \\ \langle \mathbf{b}_3^T \mathbf{A}, \mathrm{Q} \rangle \end{array} \right\} \\ &= \left\langle \begin{pmatrix} \mathbf{b}_1^T \mathbf{A} \\ \mathbf{b}_2^T \mathbf{A} \\ \mathbf{b}_3^T \mathbf{A} \end{pmatrix}, \mathrm{Q} \right\rangle = \left\langle \begin{pmatrix} \mathbf{b}_1^T \\ \mathbf{b}_2^T \\ \mathbf{b}_3^T \end{pmatrix} \mathbf{A}, \mathrm{Q} \right\rangle \\ &= \langle \mathbf{B}\mathbf{A}, \mathrm{Q} \rangle. \end{aligned} \quad \text{(S7.15)}$$
